# Scaffold Affinity Tunes Biomolecular Condensate Function

**DOI:** 10.64898/2026.08.30.748146

**Authors:** Andres Reyna, Madyson O. Briggs, Alexander F. Russell, Tien M. Phan, Ryan J. Wang, Reuben Allen, Thomas R. Hinds, Ning Zheng, Jeetain Mittal, Champak Chatterjee

## Abstract

Biomolecular condensates (BMCs) organize cellular biochemistry by concentrating selected molecules into dynamic membrane-free compartments. Yet the molecular parameters that determine not only whether condensates form, but also how they behave and what they do, remain poorly defined. Here we show that scaffold binding affinity (K_d_) is a quantitative determinant of condensate phase behavior, internal dynamics and biochemical output. Using a modular SUMO-SIM system in which scaffold valency was held constant while binding affinity was systematically varied, we found that affinity governs the phase boundary, resistance to chemical perturbation, and molecular mobility of condensates in vitro and in human cells. In multicomponent mixtures, the highest-affinity scaffold dominated dense-phase composition and dynamics, revealing a hierarchical rule for condensate organization. Finally, affinity-dependent changes in condensate dynamics translated into tunable enzyme activity, establishing binding energetics as an engineerable parameter for programming condensate biochemistry.

## Main

Biomolecular condensates (BMCs) are cellular bodies that compartmentalize macromolecules without an encapsulating membrane.^1^ Condensates form through liquid-liquid phase separation (LLPS) of multivalent macromolecular components, producing a dense phase that coexists with a dilute surrounding environment.^2^ This offers a route to localize and concentrate proteins and nucleic acids, while still allowing rapid material transfer with the surrounding milieu. The biochemical functions of BMCs arise from the enrichment or exclusion of specific components, such that composition, sequence-encoded interactions, and molecular mobility become a regulatory layer.^3,4,5^ A central unresolved question is how microscopic interaction parameters determine the material properties and biochemical outputs of condensates.

The emergent properties of condensates vary widely, ranging from liquid-like assemblies in early stress granules^6^ and transcriptional condensates^7^ to gel-like states found in neurodegenerative diseases.^8^ Valency, charge patterning, client identity and sequence composition, all influence phase separation, but it remains unclear if a simple and broadly useful parameter, such as scaffold affinity (K_d_), can predict condensate behavior in complex environments.^9–11^ Indeed, the heterogeneity of native condensates complicate efforts to isolate the contribution of discrete interaction parameters and have precluded a quantitative understanding of how K_d_ may dictate condensate stability, internal dynamics, multicomponent organization and biochemical output.^12,13^ Engineering synthetic condensates for metabolic control, signaling rewiring, or therapeutic applications requires design rules that connect molecular interactions to function. Therefore, understanding the specific role of scaffold affinity in the emergent properties of condensates can provide a principled route to tune condensates across both synthetic and biological contexts.

To address this problem, we used a modular small ubiquitin-like modifier isoform 3 (SUMO3) and SUMO-interacting motif (SIM) condensate platform in which scaffold architecture and valency are held constant while the SIM sequence can be systematically varied to span a range of SUMO-binding affinities ranging from ∼1 to ∼100 µM.^14–18^ In this system, polyvalent interactions between asymmetric multimers of SUMO3 and SIM peptides drive condensation; modeling the extensive SUMO-SIM interaction networks found in the SUMO-rich promyelocytic leukemia (PML) nuclear bodies.^10,19^

Our studies revealed that scaffold affinity quantitatively controls phase separation thresholds, chemical stability and molecular mobility in vitro and in living cells. Engineering additional electrostatic interactions further strengthened affinity-dependent effects. Computational simulation established that affinity-dependent condensate behavior arises from differences in molecular contact lifetimes and interaction-network connectivity. Scaffold affinity was also predictive for the dense-phase composition and emergent dynamics in complex multi-component condensates. Finally, by recruiting the cellular oxidative stress response enzyme pyroglutamyl peptidase 1 (PGP1)^20^ to condensates, we showed that scaffold affinity correlates with enzyme activity. Collectively, our results established K_d_ as a fundamental engineerable parameter for programming the phase behavior, material properties, and enzymatic activity in condensates.

## Results

### Scaffold affinity predicts phase boundaries

Inspired by Rosen and coworkers, we employed the 91-residue protein SUMO3, herein referred to as SUMO, as a scaffold for BMC formation.^10^ The binary scaffolding platform consists of a SIM that binds SUMO with well-defined 1:1 stoichiometry at a highly conserved surface in SUMO, the SIM-binding groove that binds diverse SIMs through primarily hydrophobic interactions (Extended Fig. 1). This enabled us to systematically change the SIM component and interrogate the relationship between K_d_ and condensate properties. Our studies began with an *in trans* SUMO-SIM system consisting of *N*-terminal mEGFP fused with five copies of SUMO, mEGFP-(SUMO)_5_, and three copies of the SIM sequence from the Protein Inhibitor of Activated STAT x or PIASx, mEGFP-(PIASx)_3_ (Extended Table 1). To generate condensates where the effects of 10-fold changes in SUMO-SIM affinity (∼7 µM to ∼70 µM) are readily observable, we envisioned that a (SUMO)_5_:(SIM)_3_ system (apparent K_d ∼_70 nM^10^) may provide a good dynamic range of measurable properties at mid-nanomolar to low micromolar protein concentrations.

The phase boundary (c_sat_) observed for each SIM tested in the 5:3 asymmetric system correlated well with K_d_ measured by isothermal titration calorimetry (ITC) (Extended Table 1), with c_sat_ increasing to higher concentrations for weaker binders (Fig 1a). In contrast, condensate size and number showed no linear correlation with either K_d_, molecular weight, isoelectric point, total charged residues or hydrophobicity (Extended Fig. 2 and Extended Table 2). This is consistent with size/number being influenced by nucleation, growth, and coalescence kinetics rather than by scaffold interaction affinity alone.

**Figure 1.**
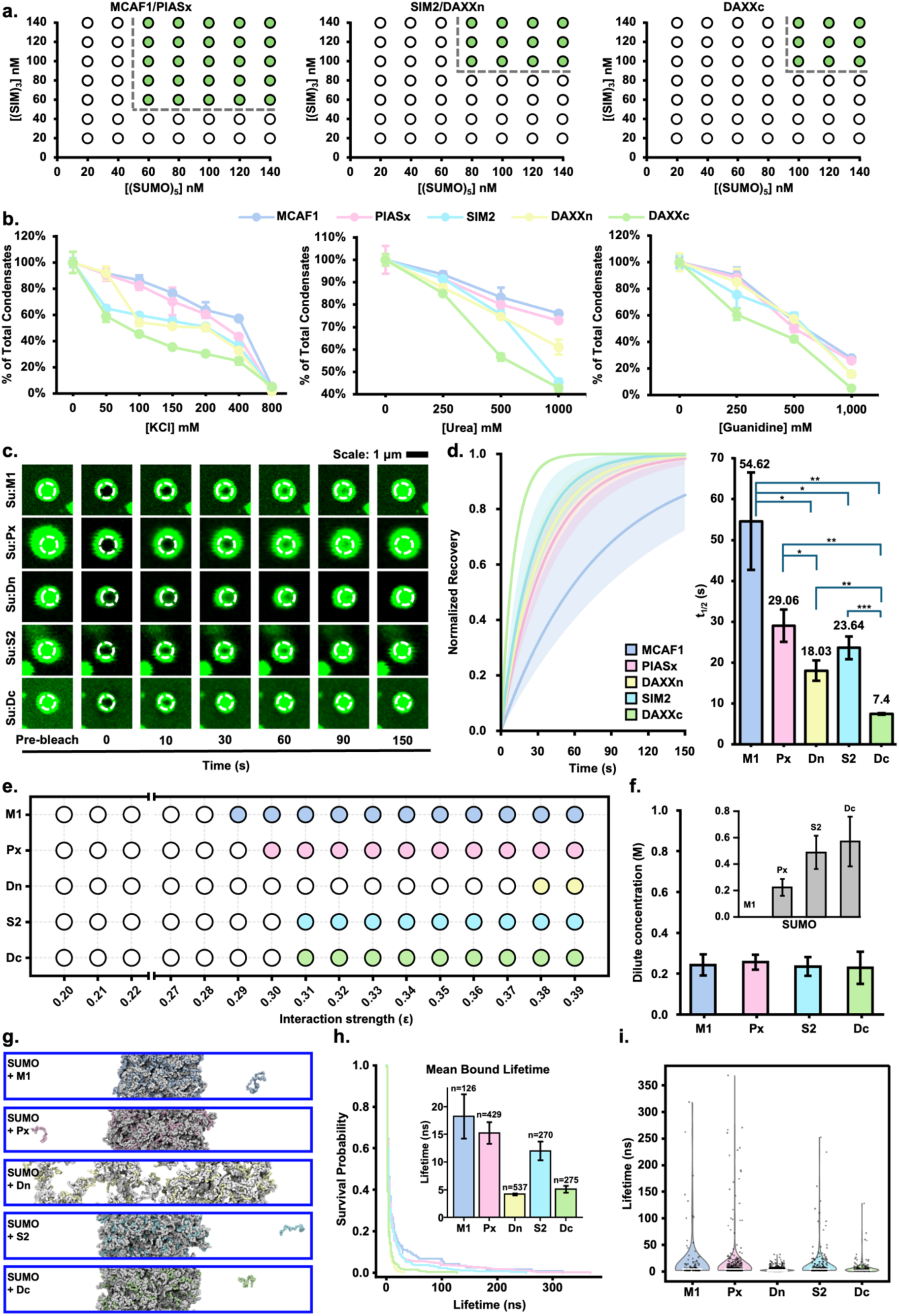
Scaffold affinity dictates the formation, stability and internal dynamics of condensates. **a**, Phase diagrams for condensate formation by mEGFP-(SUMO)_5_ and either mEGFP-(MCAF1)_3_ or mEGFP-(PIASx)_3_ (left), mEGFP-(SIM2)_3_ or mEGFP-(DAXXn)_3_ (center), and mEGFP-(DAXXc)_3_ (left). Unfilled circles denote protein concentrations at which phase separation was not observed and filled circles denote concentrations at which condensates were observed. Dashed lines indicate approximate phase boundaries. **b,** Condensates formed by various mEGFP-(SIM)_3_ constructs were treated with increasing concentrations of KCl (left), urea (center) and guanidinium chloride (right). The number of condensates within the same field-of-view per well was normalized to the maximum number of condensates observed in the absence of salt or chaotropes. n≥6 independent replicates, and error bars show s.e.m. **c,** Time-lapse images of fluorescence recovery after photobleaching (FRAP) experiments in single condensates formed by MCAF1 (M1), PIASx (Px), DAXXn (Dn), SIM2 (S2) and DAXXc (Dc). Rows are arranged from top to bottom in order of decreasing FRAP half-time (t_1/2_). Scale bar is 1 µm. **d,** Solid lines represent a single-exponential recovery curve generated from the mean of individually fitted FRAP t_1/2_ values. Shaded areas indicate recovery curves generated from the 95% confidence interval of the mean t_1/2_ for each SUMO-SIM pair (left). Average FRAP t_1/2_ for the indicated condensates calculated from fitting all normalized recovery curves (right). SIMs are arranged from left to right in order of increasing K_d_ for SUMO binding. n= 6-11 independent replicates, and error bars show s.e.m. Statistical significance was calculated using Welch’s two-tailed t-test that does not assume homoscedasticity. *= p< 0.05, **= p< 0.01, ***= p<0.001. **e,** Coarse-grained (CG) simulation phase diagrams for co-condensation of (SUMO)₅ with each (SIM)₃ construct as a function of SUMO-SIM interaction strength (ε). Open circles denote ε values at which stable phase separation was not observed, and filled circles denote ε values at which a stable condensate formed. **f,** Dilute-phase concentrations of each (SIM)₃ construct and corresponding (SUMO)₅ (inset) from 5 µs CG slab simulations at ε = 0.33. Error bars are estimated using block averages with five blocks. **g,** Representative snapshots at ε = 0.33 show phase separation for M1, Px, S2 and Dc, whereas Dn does not form a stable condensate. (SUMO)₅ is shown in gray; (SIM)₃ constructs are colored as in panel e. **h,** Survival probability curves for the SUMO-SIM bound state from AAMD simulations per SIM peptide. Inset shows mean bound lifetime (τ) with the number of complete unbinding events (n) per SIM. Error bars show s.e.m. **i,** Distributions of individual bound-state lifetimes across all simulation replicas. The median of the lifetimes is shown as a white horizontal line within each violin.

### Scaffold affinity sets condensate stability

Having observed a clear correlation between K_d_ and phase separation boundaries, we asked if stronger interactions stabilize the dense-phase network against chemical perturbation by cellular hydrotropes (ATP), electrolytes (KCl), and denaturants (urea and guanidine). To answer this, each condensate was produced at concentrations above c_sat_ and incubated with increasing amounts of ATP, KCl, urea or guanidinium chloride (Fig. 1b). Interestingly, all concentrations of ATP tested, up to 10 mM, showed no significant reduction in condensates over 24 h (Supplementary Fig. S1). This suggested that physiological concentrations of ATP, which can reach up to 8 mM,^21^ should not inhibit condensate formation by trivalent SUMO-SIM scaffolds in cells. In contrast, all condensates were sensitive to increasing concentrations of KCl, urea and guanidine (Fig. 1b), including those generated by tight-binding SIMs, such as MCAF1 (K_d_ ∼4 µM) and PIASx (K_d_ ∼8 µM). This was seen by the presence of less condensates as the concentration of salt/denaturants increased (Extended Fig. 3). As little as 50 mM KCl or 250 mM urea/guanidine led to a smaller number of condensates for weaker binding SIMs such as SIM2 (K_d_ ∼64 µM) and DAXXc (K_d_ ∼70 μM), however, PIASx and MCAF1 condensates were less sensitive to low concentrations of salt/denaturant (Fig. 1b). Overall, SUMO-SIM binding affinity correlated well with the number of detectable condensates in the presence of salt/chaotrope, indicating that stability is governed by the structured SUMO-SIM interaction network.

### Scaffold affinity predicts intra-condensate dynamics

Cellular condensates contain many different protein and/or nucleic acid components. This poses a steep challenge toward understanding how binding affinities of the scaffolds influence emergent dynamics. We tackled this question by applying fluorescence recovery after photobleaching (FRAP) to measure the intra-condensate dynamics of dense phases formed by a series of SUMO-SIM pairs. In this platform, the time required to recover fluorescence in a bleached spot is an indicator of the dynamic nature, or mobility, of condensate components. For each FRAP experiment, a circular area corresponding to ∼50% of the visible condensate was photobleached and the time required to recover fluorescence was measured (Supplementary Video 1). Condensates formed by the tighter binding MCAF1 and PIASx SIMs recovered fluorescence slower and had longer recovery half-times (t_1/2_) than those containing weaker binders such as SIM2 or DAXXc (Fig. 1c-d and Supplementary Fig. S2). The faster FRAP recovery suggests that condensates with weaker binding scaffolds exhibit more rapid molecular redistribution. Consistent with internal reorganization being a major contributor to fluorescence recovery, halving the initial concentration of weaker-binding SIMs did not significantly change t_1/2_ values (Extended Fig. 4). Thus, affinity-dependent FRAP trends primarily reflect protein redistribution within condensates rather than mass transport from the bulk solution.

### Molecular-scale contact lifetimes underlie affinity-dependent condensate dynamics

To identify a molecular basis for the observed correlations between condensate behaviors and scaffold affinity, we performed coarse-grained (CG) coexistence simulations of the (SUMO)₅-(SIM)₃ systems, using the HPS-Urry model, which has previously been shown to faithfully recapitulate phase behavior of intrinsically disordered and multidomain proteins in silico (see Methods for details).^22,23^ Phase diagrams constructed by varying a global scaling parameter for SUMO-SIM interaction strength (ε) revealed a clear hierarchy in the critical ε required for phase separation, following the trend MCAF1 < PIASx < SIM2 ≈ DAXXc << DAXXn (Fig. 1e), where lower values of ε correspond to a greater propensity for condensation. The increasing order closely recapitulated the experimentally observed variation in phase separation across SIM variants, with DAXXn being the sole outlier. At a representative interaction strength of ε = 0.33, the dilute-phase concentration of SUMO increased systematically as SIM binding affinity weakened, with SUMO fully partitioned into the dense phase for MCAF1 and progressively lesser partitioning into the dense phase for PIASx, SIM2 and DAXXc (Fig. 1f, inset). The dilute-phase SIM concentrations remained comparable across all variants (Fig. 1f). Simulation snapshots at ε = 0.33 showed stable condensate formation for MCAF1, PIASx, SIM2 and DAXXc, whereas DAXXn did not exhibit a stable condensate (Fig. 1g). The unusually elevated ε threshold for DAXXn is likely due to its concentrated negative charge (q = −7, DDDDEDE acidic cluster) introducing electrostatic repulsion with the negatively charged SUMO surface. Additionally, CG simulations do not fully capture the specific SUMO-SIM binding interactions that govern affinity at the atomic level, which may underlie the deviation from trends observed for this single variant. Intermolecular contact maps within the dense phase further revealed that the interaction network is dominated by distributed electrostatic contacts between oppositely charged residues along the scaffold sequences, with localized enrichment of contacts involving the SIM core motif and SUMO (Extended Fig. 5).

To further dissect the molecular basis of our observations, we performed all-atom (AA) molecular dynamics simulations initialized from Boltz2-predicted SUMO-SIM complex structures, using the Amber99SBwsc-STQ’ force field with multiple independent replicas per SIM peptide (see Methods for experimental details).^24^ Survival probability analysis of SUMO-SIM bound states revealed that mean bound lifetimes (τ) followed the rank order MCAF1 > PIASx > SIM2 > DAXXc > DAXXn (Fig. 1h), which was consistent with the measured trends in FRAP half-times (Fig. 1d), with only DAXXn showing a significant difference. This deviation likely arises because the single-pair AA simulations do not explicitly capture multivalent rebinding dynamics within the condensed phase, where multiple transient interactions between DAXXn and neighboring SUMO molecules may collectively extend effective residence times beyond those predicted from individual binding events. Distributions of individual bound-state lifetimes further showed that tighter-binding SIMs such as MCAF1 and PIASx exhibited broad distributions with long-lived binding events extending beyond 300 ns, whereas the weaker-binding SIM, DAXXc, was characterized by predominantly short-lived, transient contacts (Fig. 1i). Residue-level contact probability maps identified the region spanning the first α-helix and second β-strand of SUMO (residues 28-48) as the primary binding interface for the SIM hydrophobic core (Extended Fig. 6). Collectively, these results established that the measured K_d_-dependent hierarchy in condensate dynamics reflects differences in the lifetime and specificity of contacts at the SUMO binding groove, and that these molecular-scale kinetic differences also propagate to mesoscale properties.

### Hydrophobic contacts in the scaffold dictate condensate properties

SIMs vary not only in SUMO-binding affinity but also in their length, pI, charged and hydrophobic residues. Therefore, to confirm that K_d_ is the primary determinant of condensate properties, a Val in the MCAF1/PIASx hydrophobic core (*N*-<u>V</u>IDL-*C*) was mutated to Ala and the effects on affinity and phase separation were measured. Consistent with the key role of the hydrophobic core in SUMO-SIM interactions,^25^ ITC revealed a 4– and 7-fold increase in K_d_ for MCAF1(V34A) and PIASx(V4A), respectively (Fig. 2a). As expected from their weaker binding, the mutant scaffolds mEGFP-(PIASx(V4A))_3_ and mEGFP-(MCAF1(V34A))_3_ exhibited higher c_sat_ for phase separation than their wild-type (WT) counterparts (Fig. 2b). Their condensates were also significantly more sensitive to salt and denaturant (Fig. 2c) and more dynamic than WT SIMs in FRAP studies (Figure 2d-e and Supplementary Fig. S3). Altogether these tests confirmed that SUMO-SIM binding affinity, dictated by hydrophobic residues in the SIM core, primarily determines condensate stability and dynamics.

**Figure 2.**
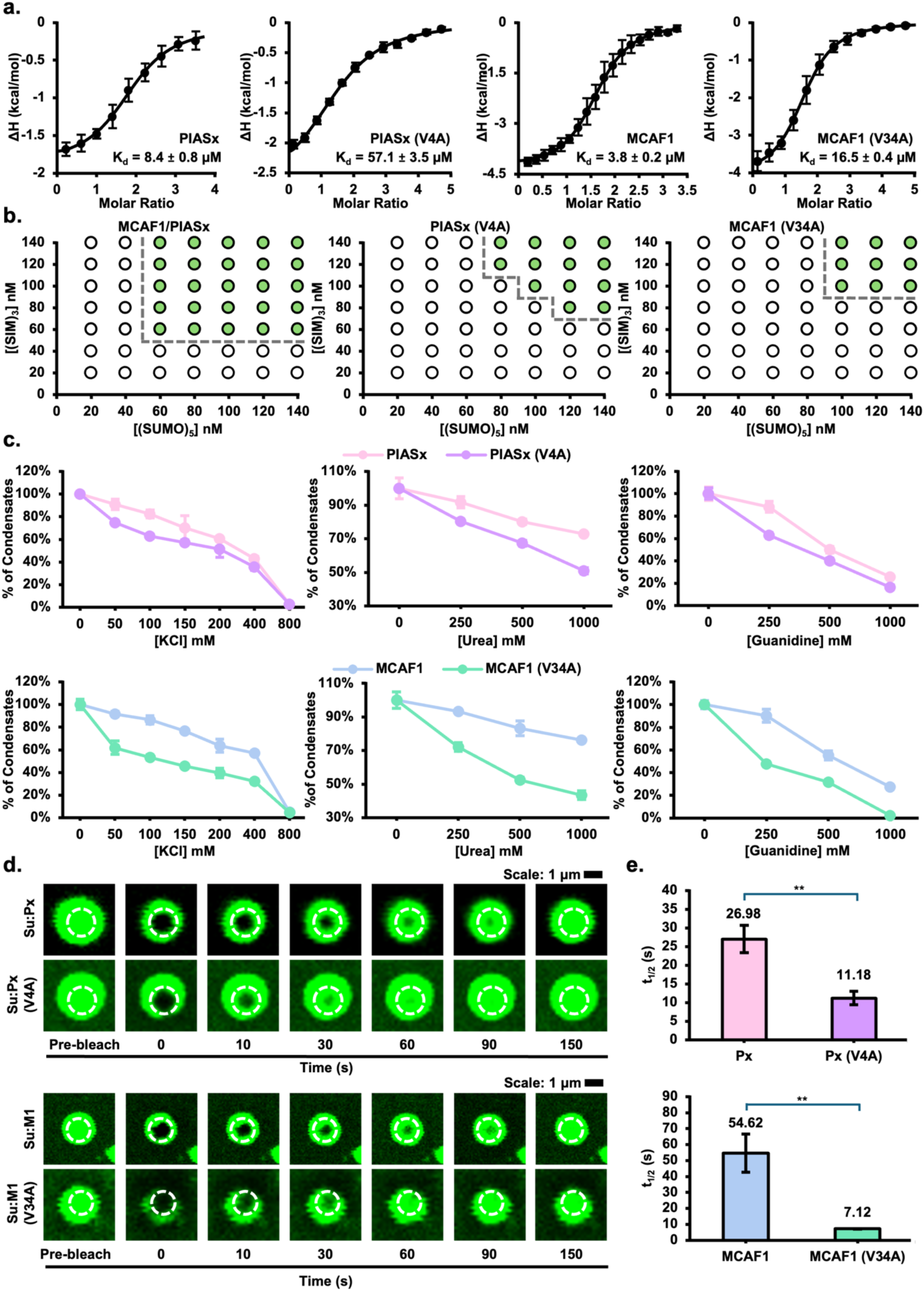
Hydrophobic core residues in SIM sequences dictate the formation, stability and internal dynamics of condensates. **a**, Isothermal titration calorimetry (ITC) binding isotherms for SUMO-binding by the indicated synthetic SIM peptides. n≥3 and error bars show s.d. **b,** Phase diagrams for condensate formation by mEGFP-(SUMO)_5_ and either mEGFP-(MCAF1)_3_ or mEGFP– (PIASx)_3_ (left), mEGFP-(PIASx(V4A))_3_ (center) or mEGFP-(MCAF1(V34A))_3_ (right). Unfilled circles denote protein concentrations at which phase separation was not observed and filled circles denote concentrations at which condensates were observed. Dashed lines indicate approximate phase boundaries. **c,** Salt– and chaotrope-dependent dissolution of preformed condensates assembled with PIASx or PIASx(V4A) (top row) and MCAF1 or MCAF1(V34A) (bottom row). Condensates formed by various mEGFP-(SIM)_3_ constructs were treated with increasing concentrations of KCl (left), urea (center) and guanidinium chloride (right). The number of condensates within the same field-of-view per well was normalized to the maximum number of condensates observed in the absence of salt or chaotropes. n=6 independent replicates, and error bars show s.e.m. **d,** Time-lapse images of FRAP experiments in single condensates formed by PIASX (Px), PIASx(V4A), MCAF1 (M1) and MCAF1(V34A). Scale bar is 1 µm. **e,** Average FRAP t_1/2_ for the indicated condensates calculated from fitting the normalized recovery curves. SIMs are arranged from left to right in order of increasing K_d_ for SUMO binding. n=7-11 independent replicates, and error bars show s.e.m. Statistical significance was calculated using Welch’s two-tailed t-test. **= p< 0.01.

### Electrostatic interactions enhance condensate stability

In addition to core hydrophobic interactions, we asked if scaffold affinity could be enhanced by engineered electrostatic interactions to further modulate condensate properties. An acidic patch on the surface of SUMO (Fig. 3a) was proposed to mediate inter-SUMO interactions at pH<7.0 when surface histidines in SUMO are protonated.^26^ To avoid the pH-dependence of this interaction, we appended a positively charged sequence, RKRKRKRKRK or (RK)5, to the C-terminus of mEGFP-(PIASx)_3_.^10^ Confirming the additive role of hydrophobic and engineered electrostatic interactions, the mEGFP-(PIASx)_3_-(RK)_5_ construct shifted to lower c_sat_ than mEGFP-(PIASx)_3_ and yielded more and larger condensates (Fig. 3b-d). To test the contribution of the positive patch, it was halved to generate mEGFP-(PIASx)_3_-RKRKR, which gratifyingly yielded condensates with properties in-between PIASx and PIASx-(RK)_5_ (Fig. 3b-d). The positive patch also rendered condensates more stable to salts and chaotropes (Fig. 3e). FRAP experiments revealed increasing t_1/2_ with increasing positive charge (Fig. 3f-g). Our efforts to quantitate SUMO-binding affinity of the synthetic 50-mer PIASx-(RK)_5_ peptide, generated by native chemical ligation^27^ of two fragments due to its large size (Extended Fig. 7a-g), were limited by poor solubility of the PIASx-(RK)_5_ peptide at pH 7.5. At pH 6.5 where PIASx-(RK)_5_ is more soluble, robust SUMO-binding was observed whereas the PIASx peptide alone did not bind SUMO (Extended Fig. 7h-k). Thus, we show that scaffold affinity can be further enhanced by additional electrostatic interactions to tune the properties of SUMO-scaffolded condensates.

**Figure 3.**
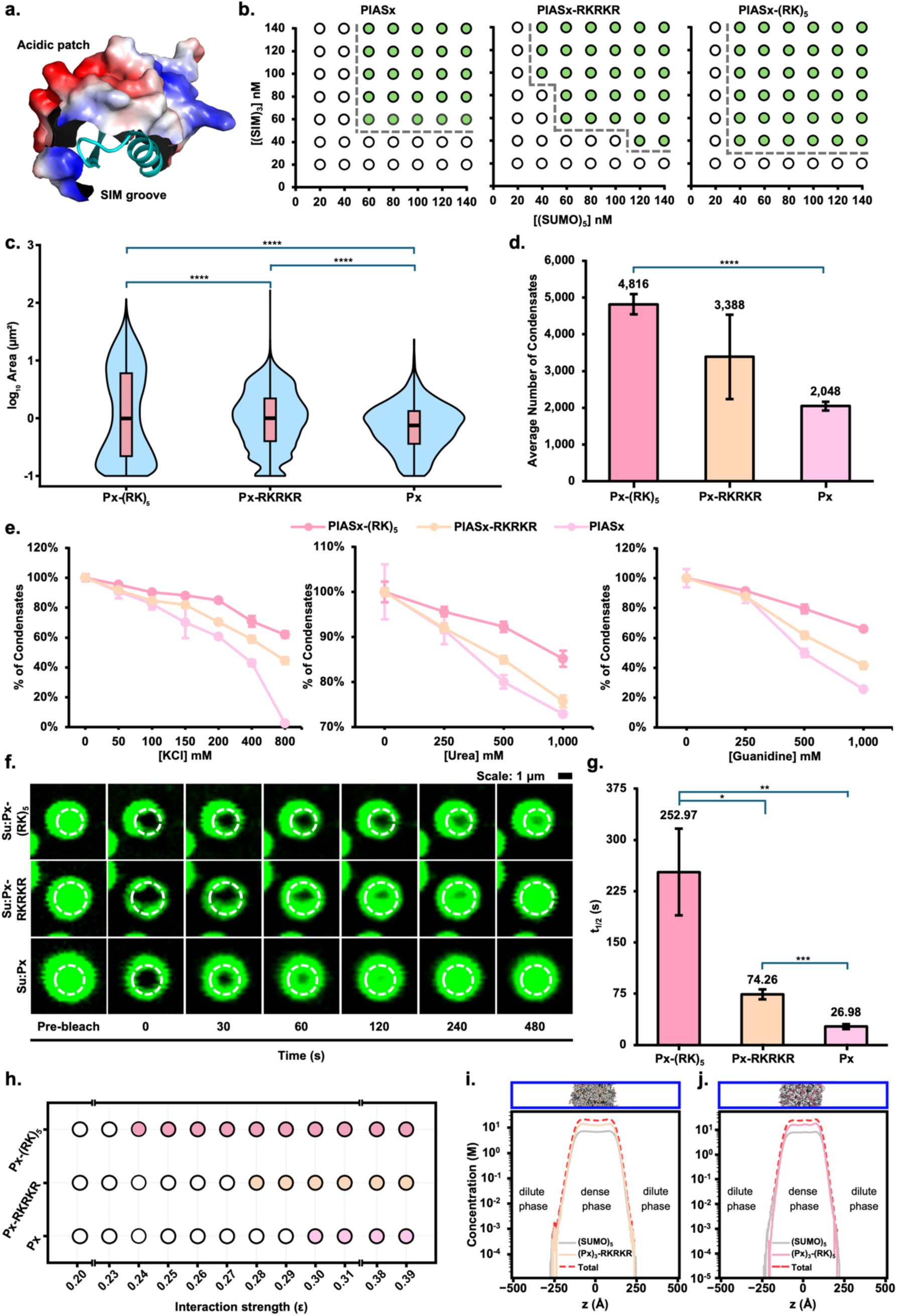
Electrostatic interactions fine-tune intra-condensate dynamics. **a**, Electrostatic surface of SUMO-3 (PDB code 2I01) showing the α-helix and β-strand (cyan) lining the SIM-binding groove and the distal acidic patch (red). **b,** Phase diagrams for condensate formation by mEGFP-(SUMO)_5_ with mEGFP-(PIASx)_3_ (left), mEGFP-(PIASx)_3_-RKRKR (center) and mEGFP-(PIASx)_3_-(RK)_5_ (right). Unfilled circles denote protein concentrations at which phase separation was not observed and filled circles denote concentrations at which condensates were observed. Dashed lines indicate approximate phase boundaries. **c,** Size-distributions of condensates formed by mEGFP-(SUMO)_5_ and the indicated PIASx constructs shown as violin plots. Box-and-whisker overlay indicates the median area (center line), interquartile range (box), and the full range of sizes (whiskers). ****= p<0.0001. **d,** Average number of condensates observed in the same field-of-view for condensates formed with mEGFP-(SUMO)_5_ and the indicated PIASx constructs. n=6 independent replicates, and error bars show s.e.m. Statistical significance was calculated using Welch’s two-tailed t-test. ****= p< 0.0001. **e,** Condensates formed by various mEGFP-(SIM)_3_ constructs were treated with increasing concentrations of KCl (left), urea (center) and guanidinium chloride (right). The number of condensates within the same field-of-view per well was normalized to the maximum number of condensates observed in the absence of salt or chaotropes. n=6 independent replicates, and error bars show s.e.m. **f,** Time-lapse images of FRAP experiments in single condensates formed by PIASx-(RK)_5_, PIASx-RKRKR and PIASx (Px). Rows are arranged from top to bottom in order of decreasing FRAP half-time (t_1/2_). Scale bar is 1 µm. **g,** Average FRAP t_1/2_ for the indicated condensates calculated from fitting the normalized recovery curves. n=7-13 independent replicates, and error bars show s.e.m. Statistical significance was calculated using Welch’s two-tailed t-test. *= p< 0.05, **= p< 0.01, ***= p<0.001. **h,** CG simulation phase diagrams for co-phase separation of (SUMO)₅ with (Px)₃-(RK)₅, (Px)₃-RKRKR and (Px)₃ as a function of SUMO–SIM interaction strength (ε). Open circles denote ε values at which stable phase separation was not observed, and filled circles denote ε values at which a stable condensate formed. Constructs are arranged from top to bottom in order of decreasing phase separation propensity. **i**-**j**, Representative snapshot and concentration profiles along the slab axis (z) from a 5 µs CG slab simulation of (SUMO)₅ with (Px)₃-RKRKR (i) and (Px)₃-(RK)₅ (j) at ε = 0.33. (SUMO)₅ is shown in gray and RK-constructs colored as in panel h. The dashed red line indicates the total concentration profile. Dilute and dense phase regions are labeled.

To elucidate the molecular basis by which polybasic extensions modulate condensate properties, we performed CG coexistence simulations of (SUMO)₅ with (PIASx)₃-RKRKR and (PIASx)₃-(RK)₅. Phase diagrams revealed an RK-length-dependent propensity for phase separation, with PIASx-(RK)₅ condensing at the lowest ε, followed by PIASx-RKRKR and then PIASx (Fig. 3h), perfectly recapitulating the measured hierarchy in c_sat_. Simulations at ε = 0.33 showed that both RK-extended constructs formed stable condensates with complete partitioning of SUMO and SIM components into the dense phase (Fig. 3i–j). Intermolecular contact maps within the dense phase revealed that, while the canonical SUMO-SIM binding interface is preserved, polybasic extensions introduce additional distributed contacts with negatively charged regions on the SUMO surface (Extended Fig. 8). These electrostatic interactions are more pronounced in PIASx-(RK)₅ than in PIASx-RKRKR, resulting in a broader and more interconnected interaction network. These results demonstrate that electrostatic interactions stabilize the dense phase by increasing interaction multiplicity and promoting a more interconnected interaction network.

### Intra-condensate dynamics of complex condensates are dictated by the highest affinity scaffold

Nuclear PML bodies contain several proteins that bind SUMO, and it remains unclear how proteins with varying affinities contribute to the biophysical properties of the heterogeneous PML body.^28^ To address this question, we investigated the properties of ternary and quaternary condensates formed by a mixture of SIMs. Condensates were initially generated from the weakest affinity mEGFP-(SUMO)_5_ and mEGFP-(DAXXc)_3_ pair, followed by adding the tighter binding mEGFP-(MCAF1)_3_ protein (Fig. 4a). This led to the formation of well-defined puncta on glass slides (Fig. 4b). Control condensates for direct comparison were generated with mEGFP-(SUMO)_5_ and the corresponding mEGFP-(SIM)_3_ (Supplementary Fig. S4). FRAP experiments with the ternary mixtures revealed that t_1/2_ resembled the less dynamic MCAF1-condensates (Fig. 4c). Consistent with this, SDS-PAGE analysis of the separated condensates revealed an excess of the tighter binding MCAF1 in the dense phase (Fig. 4d). An increase in t_1/2_ was also observed when condensates initially generated with mEGFP-(DAXXc)_3_ were mixed with free mEGFP-(PIASx)_3_-(RK)_5_ (Extended Fig. 9). More complex condensates formed by quaternary mixtures of mEGFP-(SUMO)_5_, mEGFP-(DAXXc)_3_, mEGFP-(MCAF1)_3_ and mEGFP-(PIASx)_3_-(RK)_5_ also revealed t_1/2_ that closely resembled the tightest SUMO-binding SIM (Fig. 4b and Fig. 4e-f and Supplementary Fig. S5), with the dense phase enriched in mEGFP-(PIASx)_3_-(RK)_5_ (Fig. 4f). This suggests that SUMO-scaffolded condensates are not kinetically trapped and can undergo thermodynamic equilibration at room temperature, leading to their properties being dictated by the strongest binding partners.

**Figure 4.**
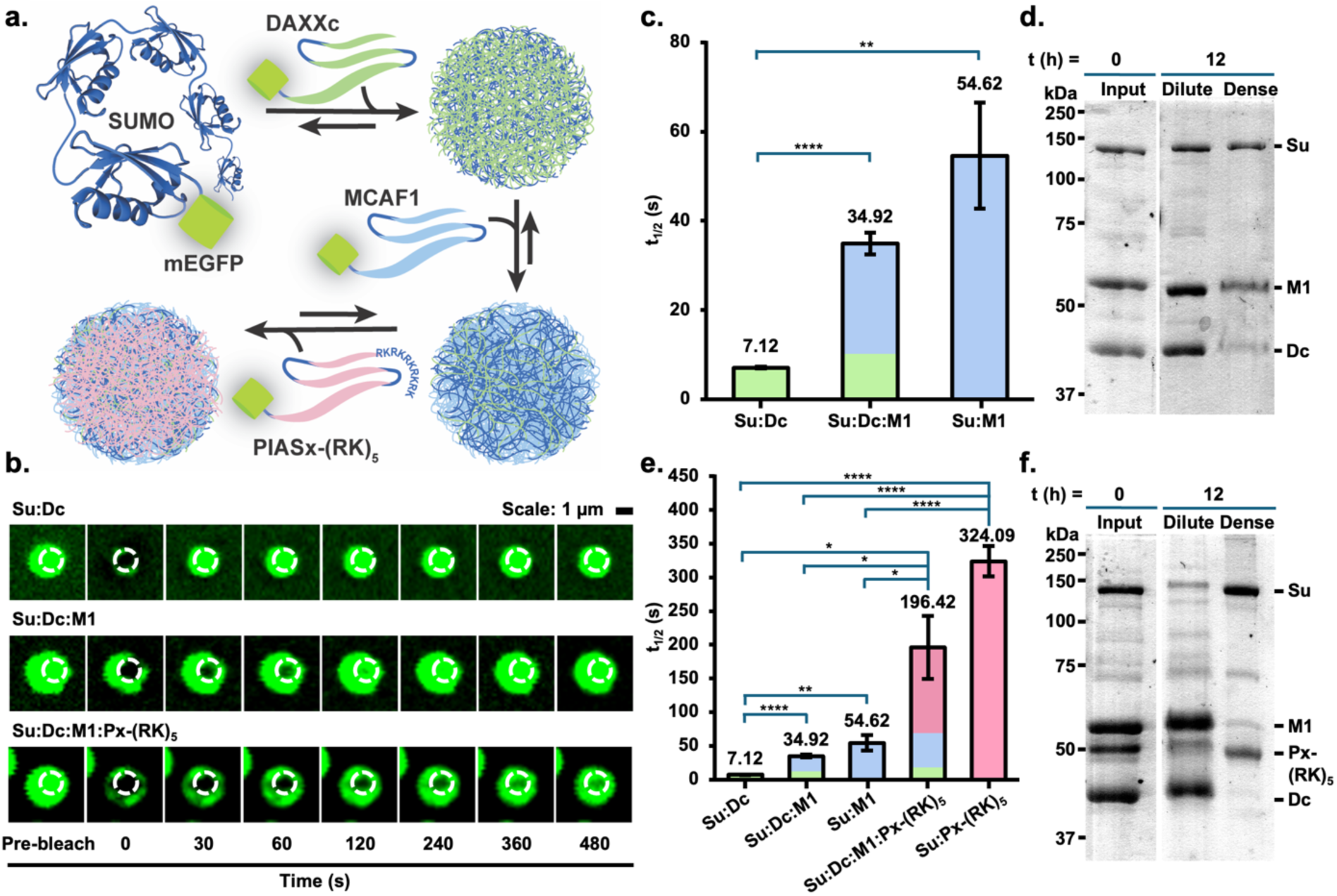
Intra-condensate dynamics of complex condensates are dictated by the highest affinity scaffold. **a**, Schematic depiction of the stepwise assembly of multi-component condensates. Phase separation was initiated by the mEGFP-(SUMO)_5_ scaffold and the weakest binding SIM, followed by subsequent additions of increasingly stronger binding scaffolds to yield complex condensates. **b,** Time-lapse images of FRAP experiments in single condensates formed by a binary mixture of mEGFP-(SUMO)_3_ (Su) and mEGFP-(DAXXc)_3_ (Dc) (top row), a ternary mixture of Su, Dc and mEGFP-(MCAF1)_3_ (M1) (middle row) and a quaternary mixture containing Su, Dc, M1 and mEGFP-(PIASx)_3_-(RK)_5_ (bottom row). Rows are arranged from top to bottom in order of increasing FRAP half-time (t_1/2_). Scale bar is 1 µm. **c,** Average FRAP t_1/2_ for the dense phase formed by ternary mixtures calculated from fitting the normalized recovery curves. n= 7-11 independent replicates, and error bars show s.e.m. Statistical significance was calculated using Welch’s two-tailed t-test. ****= p< 0.0001 and **= p< 0.01. **d,** Analysis of the dilute and dense phases in a ternary mixture of scaffolds by 12% SDS-PAGE showing enrichment of the tighter binding scaffold M1 in the dense phase. **e,** Average FRAP t_1/2_ for the dense phase formed by ternary and quaternary mixtures, calculated from fitting the normalized recovery curves. n=6 to 11 independent replicates, and error bars show s.e.m. Statistical significance was calculated using Welch’s two-tailed t-test. ****= p< 0.0001, **= p< 0.01, *= p<0.05. **f,** Analysis of the dilute and dense phases in a quaternary mixture of scaffolds by 10% SDS-PAGE showing enrichment of the tightest binding scaffold PIASx-(RK)_5_ in the dense phase.

### Scaffold affinities predict intra-condensate dynamics in human cells

Controlling condensate composition, location and function in living cells is desirable for engineering metabolic pathways,^29^ increasing therapeutic efficacy,^30^ and regulating gene transcription.^31^ To test the portability and conservation of affinity-driven condensate properties in human cells, we generated *in cis* linear fusions containing mCherry-(SUMO)_5_-(SIM)_3_. Three SIMs with varying affinities for SUMO–PIASx (∼8 µM), SIM2 (∼64 µM), and DAXXc (∼70 µM)– formed well-defined puncta in HeLa cells (Fig. 5a). The puncta were primarily cytoplasmic with sizes ranging from 1-6 µm^2^ that did not correlate with SUMO-SIM K_d_ (Fig. 5b), which was consistent with our in vitro observations. The internal dynamics of puncta were measured by FRAP and found to match the affinity-driven in vitro trends (Fig. 5c-d) such that mCherry-(SUMO)_5_-(PIASx)_3_ had the longest t_1/2_ while mCherry-(SUMO)_5_-(DAXXc)_3_ had the shortest t_1/2_ (Fig. 5d). The affinity-dependent ordering of t_1/2_ was surprisingly also conserved in HEK293T and H1299 cells, demonstrating that K_d_ robustly predicts condensate dynamics in complex cellular environments (Fig. 5e).

**Figure 5.**
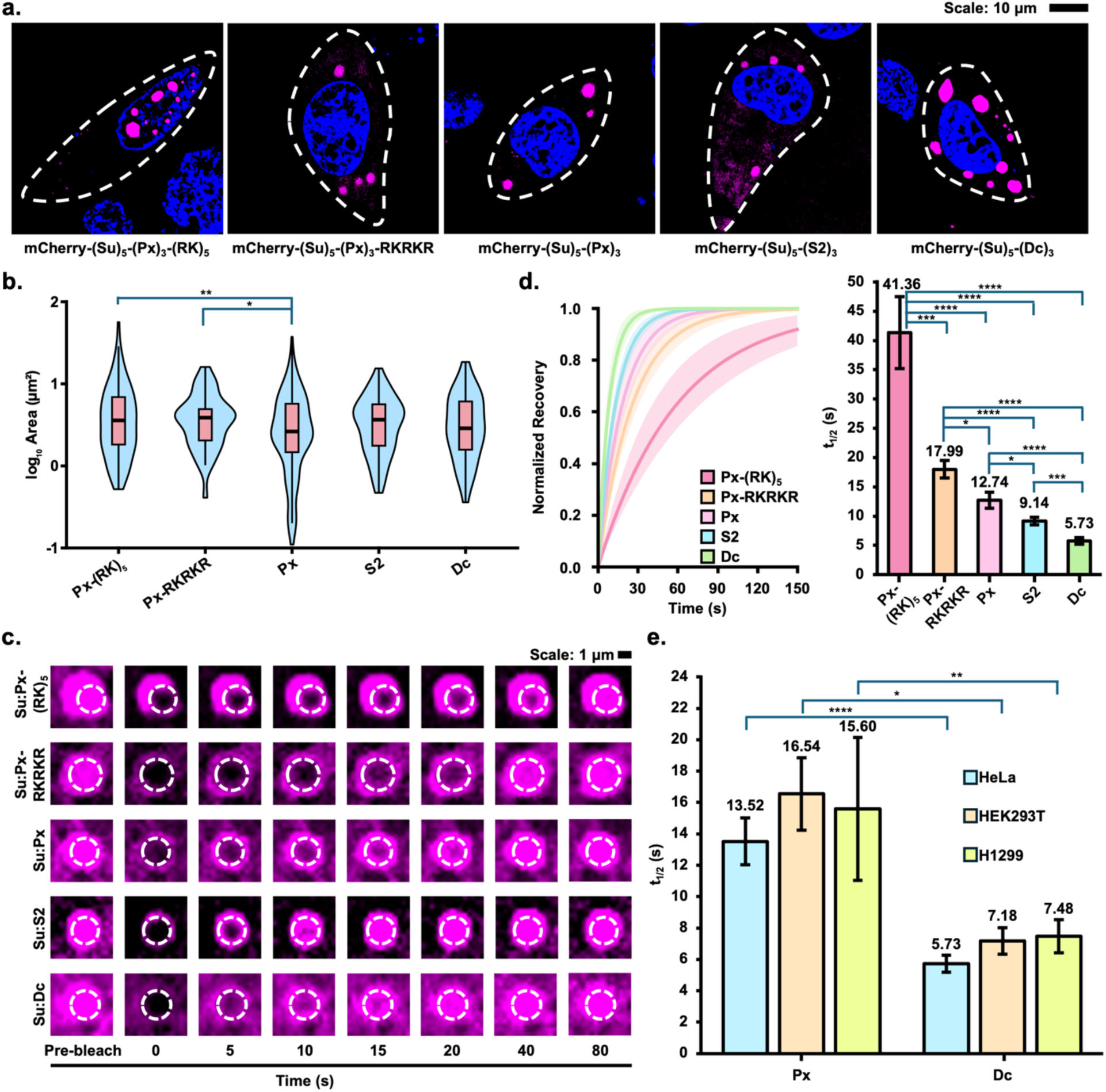
Intra-condensate dynamics in human cells reflect scaffold affinities. **a**, HeLa cells expressing the indicated mCherry-(SUMO)_5_-(SIM)_3_ variants were imaged at 48 h post-transfection. The punctate mCherry fluorescence (magenta) at 610 nm indicates condensate formation. Nuclei were stained with Hoechst 33342 dye (blue), and the cell membrane (dotted line) was labelled with wheat germ agglutinin conjugated with Alexa Fluor 647 (WGA647). Scale bar is 10 µm. **b,** Size-distributions of condensates formed by mEGFP-(SUMO)_5_-(SIM)_3_ fusion constructs shown as violin plots. Box-and-whisker overlay indicates the median area (center line), interquartile range (box), and the full range of sizes (whiskers). Statistical significance was calculated using Welch’s two-tailed t-test. *= p<0.05 and **= p<0.01. **c,** Time-lapse images of FRAP experiments in single condensates formed by the indicated mCherry-(SUMO)_5_-(SIM)_3_ constructs in live HeLa cells. Rows are arranged from top to bottom in order of decreasing FRAP half-time (t_1/2_). Scale bar is 1 µm. **d,** Solid lines represent single-exponential recovery curves generated from the mean of all individually fitted FRAP half-times (t_1/2_) for condensates formed by the indicated mCherry-(SUMO)_5_-(SIM)_3_ constructs in live HeLa cells (Left). Shaded areas indicate recovery curves generated from the 95% confidence interval of the mean t_1/2_ for each construct. Average FRAP t_1/2_ for condensates formed by the indicated mCherry-(SUMO)_5_-(SIM)_3_ constructs in live HeLa cells, calculated from fitting the normalized recovery curves (Right). n= 7-32 independent replicates and error bars show s.e.m. Statistical significance was calculated using Welch’s two-tailed t-test. *= p<0.05, ***= p<0.001, ****= p<0.0001. **e,** Average FRAP t_1/2_ for condensates formed by mCherry-(SUMO)_5_-(PIASx)_3_ or mCherry-(SUMO)_5_-(DAXXc)_3_ in live HeLa, HEK293T and H1299 cells, calculated from fitting the normalized recovery curves. n= 6-23 independent replicates and error bars show s.e.m. Statistical significance was calculated using Welch’s two-tailed t-test. *= p<0.05, **= p<0.01, ****= p<0.0001.

Based on this success, we asked if the introduction of additional electrostatic interactions could influence the internal dynamics of condensates inside cells. Interestingly, when the mCherry-(SUMO)_5_-(PIASx)_3_-(RK)_5_ protein was expressed in HeLa cells, large nuclear puncta were observed with much longer t_1/2_ in FRAP experiments than mCherry-(SUMO)_5_-(PIASx)_3_ alone (Fig. 5a and Fig. 5c-d). The atypical location of these puncta was attributed to the (RK)_5_ sequence acting as a monopartite classical nuclear localization sequence.^32^ Indeed, truncating the RK repeats in mCherry-(SUMO3)_5_-(PIASx)_3_-RKRKR was sufficient to regain cytoplasmic localization (Fig. 5a) and condensates exhibited the expected intermediate FRAP t_1/2_ times (Fig. 5c-d). Collectively, our results show that the effect of scaffold affinity on molecular interaction networks is conserved in complex environments, which enables tailoring condensate properties.

### Scaffold affinities regulate enzyme activity in condensates

Having established scaffold affinity as a predictive parameter for emergent properties in living cells, we asked if it also correlates with enzyme function, which could provide new strategies for fine-tuning the biochemical output of condensates. Previous studies demonstrated increased or decreased enzymatic activity based on the scaffold,^33^ active site conformation^34^ and substrate accumulation.^35^ However, the relationship between scaffold affinity and enzymatic activity has not been systematically explored. Due to its widespread tissue distribution and important roles in regulating a range of human hormones including neurotensin and gastrin, we focused on the enzyme pyroglutamyl peptidase 1 (PGP1) that hydrolyzes a pyroglutamic acid residue from the *N*-termini of peptides.^36^

To visualize and quantify PGP1 activity within condensates we generated two AlexaFluor568 (AF568)-labeled PGP1 fusions bearing three SIM repeats– AF568-PGP1-(PIASx)_3_ and AF568-PGP1-(DAXXc)_3_ (Supplementary Fig. S6). A fluorogenic substrate, pGlu-HCA, in which L-pyroglutamic acid is conjugated to a hemicyanine (HCA) fluorophore, was synthesized (see Methods and Supplementary Fig. S7 for details).^37^ PGP1 activity was measured by near-IR fluorescence of the hydrolyzed HCA product at 700 nm. Steady-state kinetic analysis in solution showed the two fusions had similar catalytic efficiencies, with k_cat_/K_M_ of 8.5 ± 1.0 x 10^4^ M^−1^s^−1^ for AF568-PGP1-(PIASx)_3_ and 6.2 ± 2.8 x 10^4^ M^−1^s^−1^ for AF568-PGP1-(DAXXc)_3_ (Fig. 6a).

**Figure 6.**
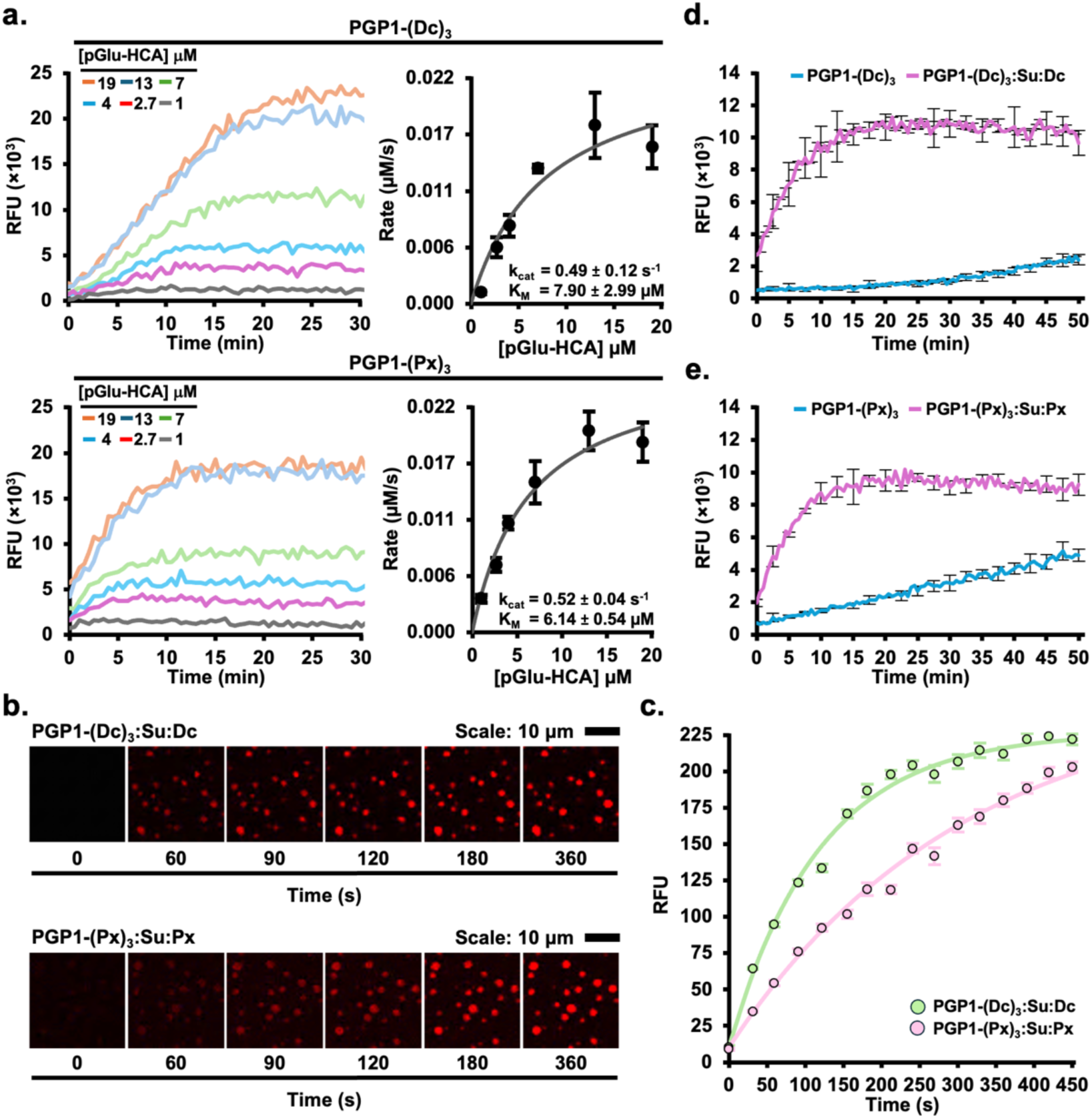
Scaffold affinities regulate enzyme activity in condensates. **a**, Enzyme progress curves showing HCA generation at λ_em_ 700 nm after incubating 50 nM of either PGP1-(DAXXc)_3_ or PGP1-(PIASx)_3_ with increasing concentrations of the probe pGlu-HCA (Left). Michaelis-Menten plots of initial reaction velocities versus pGlu-HCA concentrations for PGP1-(DAXXc)_3_ and PGP1-(PIASx)_3_ (Right). **b,** Time-lapse images of the appearance of HCA fluorescence following the addition of pGlu-HCA to mEGFP-(SUMO)_5_+mEGFP-(DAXXc)_3_ condensates containing AF568-PGP1-(DAXXc)3 (Top row) and mEGFP-(SUMO)_5_+mEGFP-(PIASx)_3_ condensates containing AF568-PGP1-(PIASx)_3_ (Bottom row). Images shown at the indicated time points show product accumulation (red puncta) within condensates. Scale bar, 10 µm. **c,** Increase in HCA fluorescence over time in 60 discrete droplets from time-lapse images of AF568-PGP1-(DAXXc)_3_ activity in mEGFP-(SUMO)_5_:mEGFP-(DAXXc)_3_ condensates (green trace) and AF568-PGP1-(PIASx)_3_ activity in mEGFP-(SUMO)_5_:mEGFP-(PIASx)_3_ condensates (pink trace). Error bars show s.e.m. **d,** Increase in HCA fluorescence over time in a 96-well plate-based assay of AF568-PGP1-(DAXXc)_3_ activity in a mixture of dilute and dense phases containing mEGFP-(SUMO)_5_:mEGFP-(DAXXc)_3_ condensates (pink trace) or in buffer alone (blue trace). n=3 independent replicates and error bars show s.d. **e,** Increase in HCA fluorescence over time in a 96-well plate-based assay of AF568-PGP1-(PIASx)_3_ activity in a mixture of dilute and dense phases containing mEGFP-(SUMO)_5_:mEGFP-(PIASx)_3_ condensates (pink trace) or in buffer alone (blue trace). n=3 independent replicates and error bars show s.d.

We next asked if recruitment to compositionally distinct condensates alters PGP1 activity. Matched AF568-PGP1-SIM fusions were recruited to pre-assembled SUMO:PIASx or SUMO:DAXXc condensates, and pGlu-HCA hydrolysis was monitored by the appearance of HCA fluorescence. The substrate partitioned efficiently and similarly into both condensate types, with encapsulation efficiencies of 89.5 ± 0.7% for SUMO:PIASx and 88.4 ± 0.5% for SUMO:DAXXc, indicating that differences in product formation were not attributable to differential substrate loading. Condensates lacking PGP1 showed no increase in HCA fluorescence over time, confirming that pGlu-HCA was stable under the assay conditions (Supplementary Fig. S8). HCA fluorescence appeared more rapidly in AF568-PGP1-(DAXXc)_3_ containing SUMO:DAXXc condensates than in the corresponding SUMO:PIASx system (Fig. 6b). Quantification of individual condensates over time confirmed faster product accumulation in the more dynamic SUMO:DAXXc condensates (Fig. 6c). PGP1 activity did not correlate with condensate size (Extended Fig. 10), arguing that the observed differences arise from condensate physicochemical properties rather than from variation in the total amount of enzyme within individual droplets.

Finally, we measured PGP1 activity in bulk biphasic mixtures containing both dense and dilute phases. Recruitment of AF568-PGP1-(DAXXc)_3_ to SUMO:DAXXc condensates produced a >10-fold apparent rate enhancement relative to enzyme alone (Fig. 6d). AF568-PGP1-(PIASx)_3_ recruited to SUMO:PIASx condensates also showed enhanced activity, although the magnitude of activation was ∼10-fold relative to enzyme alone (Fig. 6e). Together, these data show that condensate recruitment enhances PGP1 activity and that the magnitude of activation is tuned by scaffold identity, with less dynamic condensates supporting reduced enzymatic activation relative to more dynamic condensates.

## Discussion

A central challenge in condensate biology is to predict how defined molecular interactions give rise to emergent phase behavior, material state and biochemical function. Here, we identify scaffold binding affinity, K_d_, as a quantitative and engineerable parameter that dictates these properties. By systematically varying SUMO-SIM affinity while preserving scaffold valency and architecture, we isolated interaction strength as an independent variable and established design rules for tuning condensate formation, stability, internal dynamics and enzymatic output.

A central finding is that scaffold affinity quantitatively tunes the phase boundary for condensation. The measured and computed relationship between K_d_ and c_sat_ showed that weaker interactions require higher scaffold concentrations to phase separate, consistent with the behavior of multivalent associative polymers.^38^ Coarse-grained and all-atom simulations provided a molecular basis for this relationship, revealing that stronger SUMO-SIM interactions lower the threshold for phase separation and promote more complete dense-phase partitioning. By contrast, condensate size and number did not scale with K_d_, separating the thermodynamic threshold for phase separation from kinetic processes that determine mesoscale morphology, including nucleation barriers, diffusion and local concentration fluctuations.^39^

Scaffold affinity also controlled condensate material properties. Tighter-binding scaffolds formed condensates that were more resistant to salt and chaotropic perturbation and exhibited slower internal rearrangement by FRAP. Mutations in the SIM hydrophobic core increased molecular mobility, supporting binding affinity, rather than sequence identity, as a dominant determinant of dynamics in this scaffold context. Thus, K_d_ provides a molecular handle for tuning condensates across a continuum from dynamic liquid-like assemblies to more slowly exchanging, viscoelastic states. Our engineered electrostatic interactions provided an additional and modular layer of control. Introducing positively charged motifs into the SIM scaffold enhanced condensate stability, lowered phase boundaries and slowed internal dynamics. Simulations suggest that these effects arise from additional distributed contacts between the positive patch and negatively charged regions on SUMO, increasing interaction multiplicity and network connectivity. The additivity of hydrophobic affinity and electrostatic complementarity indicates that scaffold properties can be rationally combined to tune condensate behavior with greater precision than either parameter alone.

The affinity framework also extended to multicomponent condensates. In ternary and quaternary mixtures, the highest-affinity scaffold disproportionately shaped dense-phase composition and dynamics, indicating that condensate properties are not simple averages of the constituent interactions. This hierarchical behavior suggests a mechanism by which cells could bias condensate identity in complex environments containing competing interaction motifs. It also provides a design principle for synthetic systems: incorporation of a high-affinity component can reprogram the composition and dynamics of a multicomponent condensate. Importantly, the affinity-dependent trends observed in vitro were retained in living cells, where K_d_ predicted intra-condensate dynamics across multiple human cell lines. This portability indicates that affinity-driven control remains robust despite the complexity of the intracellular environment, including crowding, active transport and heterogeneous molecular composition. The ability to rationally tune condensate dynamics in cells using defined interaction modules provides a foundation for engineering intracellular organization and function.

Affinity-dependent material changes also had direct biochemical consequences. Recruitment of PGP1 into condensates showed that more dynamic condensates supported higher enzymatic activity, whereas less dynamic condensates produced comparatively reduced activity. Because substrate partitioning was similar across condensates, the differences in activity likely arise from changes in molecular mobility and encounter frequency within the dense phase. Thus, scaffold affinity regulates catalysis not only by concentrating reaction components but also by defining the physical environment in which reactions occur.

Collectively, our results establish scaffold affinity as a molecular control parameter that connects microscopic binding interactions to condensate phase behavior, material state and biochemical function. In this engineered SUMO-SIM system, K_d_ predicts the concentration threshold for condensation, dense-phase stability, internal dynamics, hierarchical behavior in multicomponent mixtures and intracellular mobility. Although endogenous condensates will include additional regulatory features such as RNA, DNA, metabolites, post-translational modifications and active cellular processes, the quantitative relationships described here provide a framework for interpreting natural condensates and engineering synthetic assemblies with programmable biochemical outputs.

## Supporting information

Movie of Fluorescence Recovery after Photobleaching

## Acknowledgments

This work was supported by NSF Grant 2107525 and NIH grant R35GM149228 to C.C. and NIH grants R35GM153388 to J.M. and K99GM159055 to T.MP. Ning Zheng is an Investigator of the Howard Hughes Medical Institute. We are grateful for the computational resources provided by Texas A&M High Performance Research Computing (HPRC) and the W.M. Keck Microscopy Center at the University of Washington for confocal imaging resources. C.C. thanks Joshua Vaughan and Dan Fu for helpful discussions on live-cell imaging.

## Author Contributions

A.R. and C.C. conceptualized the mechanistic studies of condensates; A.R. performed all scaffold-cloning, protein purification, confocal imaging and enzyme kinetics experiments; T.M.P. and J.M. conceptualized the computational model of SUMO-interactions and performed coarse-grained and all atom simulations; A.F.R. synthesized the fluorescent probe of enzyme activity; M.O.B. and R.J.W. synthesized SUMO-interacting peptides and performed isothermal titration calorimetry experiments. R.A. cloned and purified SUMO. T.R.H. and N.Z. assisted with experimental design and isothermal titration calorimetry. All authors contributed to generating figures, writing and editing the manuscript.

## Methods

### General Information

DNA synthesis was performed by Integrated DNA Technologies (Coralville, IA), and Sanger sequencing was performed by Eurofins Genomics (Louisville, KY). Plasmid miniprep, PCR purification, and gel extraction kits were from Qiagen (Germantown, MD). HisPur Ni-NTA resin was from Thermo Scientific (Waltham, MA). HeLa, HEK293T, and H1299 cells were cultured in T75 flasks or 8-well µ-Slides (Ibidi, Gräfelfing, Germany) in Dulbecco’s Modified Eagle Medium (DMEM for HeLa and HEK293T) or Roswell Park Memorial Institute Medium (RPMI 1640 for H1299) supplemented with 10% fetal bovine serum at 37 °C in a humidified incubator with 5% CO₂. Routine confocal microscopy was performed on a Leica SP8 X scanning confocal microscope using Leica LAS X. PCR used Phusion High-Fidelity DNA Polymerase in HF buffer (Thermo Scientific, Waltham, MA). All buffers were prepared with 18.2 MΩ·cm purified water (Mili-Q^®^). Protein concentrations were determined either by SDS-PAGE with Coomassie staining and compared to pure bovine serum albumin (BSA) standards,^40^ or, for constructs bearing mEGFP, by absorbance at 488 nm using ε₄₈₈ = 56,000 M⁻¹ cm⁻¹.^41^ Peptides were synthesized by Fmoc solid-phase peptide synthesis on CEM Liberty Blue 1.0 and CEM Liberty PRIME 2.0 microwave-assisted synthesizers from CEM Corporation (Matthews, NC). Reversed Phase High Performance Liquid Chromatography (RP-HPLC) used mobile phases consisting of Buffer A (0.1% TFA in H₂O) and Buffer B (90% acetonitrile, 9.9% H₂O, 0.1% TFA) on Agilent 1260 Infinity II systems from Agilent Technologies (Santa Clara, CA), with C18 or C4 columns. Analytical runs used a gradient of 0–73% B over 30 min at 1.0 mL min⁻¹, and semi-preparative or preparative runs used the indicated C18 or C4 columns at 4–20 mL min⁻¹ with UV detection at 214 and 280 nm. Preparative columns were from Waters Corporation (Milford, MA) or from W.R. Grace and Co. (now Hichrom, Hungary). Routine peptide and protein mass spectra were acquired by direct-infusion electrospray ionization on a Bruker Esquire ion-trap from Bruker (Billerica, MA) or a Finnigan LTQ ion-trap from Thermo Fisher Scientific (Waltham, MA) operated in positive-ion mode.

### Molecular cloning

All PCR amplifications and site-directed mutagenesis used Phusion polymerase (Phusion High-Fidelity PCR Master Mix with HF Buffer, Thermo Scientific). The plasmid pTEV19-His_10_-mEGFP-(SUMO3)_3_ containing three repeats of the human SUMO3 gene separated by Gly– and Ser-rich flexible linkers was obtained from Addgene (plasmid #127093).^10^ A stop codon was introduced by PCR-based site-directed mutagenesis using primers listed in (Methods Table 1) to terminate translation immediately after the final SUMO3 repeat, preventing expression of the downstream TEV site. The modified pTEV19 plasmid containing five repeats of SUMO3 with intervening Gly– and Ser-rich linkers, pTEV19-His10-mEGFP-(SUMO3)_5_, was generated by restriction-digest cloning using the pTEV19-His_10_-mEGFP-(SUMO3)_3_ plasmid as the template and AatII and XhoI restriction sites (New England Biolabs (NEB), Ipswich, MA). The restriction digested vector was treated with calf intestinal phosphatase (NEB, Ipswich, MA) to prevent self-ligation and purified by gel excision from 1% agarose using the QIAquick PCR & Gel Cleanup Kit (Qiagen, Germantown, MD). The same restriction enzymes were used to digest a synthetic gene fragment encoding two additional SUMO3 repeats with Gly– and Ser-rich linkers and flanked by 5’ AatII and 3’ XhoI restriction sites (Methods Table 2). The digested insert and dephosphorylated vector were ligated with T4 DNA ligase (NEB, Ipswich, MA).

All pTEV19-His_10_-mEGFP-(SIM)_3_ constructs used in vitro were generated by restriction-digest cloning of the plasmid GFP-(SIM)_3_ obtained from Addgene (plasmid #126955).^10^ The vector was digested with BsrGI and XhoI and treated with calf intestinal phosphatase to prevent self-ligation. The same restriction enzymes were used to digest a synthetic gene fragment corresponding to each SIM peptide, separated by Gly– and Ser-rich linkers and flanked by 5’ BsrGI and 3’ XhoI restriction sites (Methods Table 2). The digested insert and dephosphorylated vector were ligated with T4 DNA ligase.

Two PIASx variants were prepared by PCR-based site-directed mutagenesis. For mEGFP-(PIASx)_3_, a stop codon was introduced using primers listed in Methods Table 2 to terminate translation immediately after the final PIASx repeat. For mEGFP-(PIASx)_3_-RKRKR, a second stop codon was introduced further downstream using primers listed in Methods Table 1. The mEGFP-(MCAF1(V34A))_3_ construct was synthesized and purified by Genscript.

To generate PGP1 fusion constructs in pTEV19, the host vector pTEV19-His_10_-mEGFP-(SUMO3)_5_, synthesized and purified by GenScript, was linearized by PCR amplification using Phusion polymerase with primers listed in Methods Table 1 to remove the mEGFP-(SUMO3)_5_ cassette and to introduce 20-40 bp overlaps compatible with HiFi assembly for insertion of either the (PIASx)_3_ or (DAXXc)_3_ fragments. The (PIASx)_3_ and (DAXXc)_3_ inserts were generated by PCR amplification from pTEV19-His_10_-mEGFP-(PIASx)_3_ and pTEV19-His_10_-mEGFP-(DAXXc)_3_, respectively, using Phusion polymerase and primers listed in Methods Table 1 designed to provide sequence overlap with the linearized host vector. Assembly reactions were performed using NEBuilder HiFi DNA Assembly Master Mix at ∼2:1 insert:vector molar ratio and incubated at 50 °C for 60 min according to the manufacturer’s guidelines.

For mammalian expression, the plasmid RFP-(SUMO)_6_-(SIM)_10_ (pC1-mCherry-(SUMO3)_6_-(PIASx)_10_) was purchased from Addgene (plasmid #122027)^10^ and served as a positive control for condensate formation in cells. pC1-mCherry-(SUMO3)_5_-(PIASx)_3_, pC1-mCherry-(SUMO3)_5_-(SIM2)_3_, pC1-mCherry-(SUMO3)_5_-(DAXXc)_3_, and pC1-mCherry-(SUMO3)_5_-(PIASx)_3_-(RK)_5_ *in cis* fusion construct genes utilized for transient transfection of human cells were custom synthesized, purified, and cloned into the host pC1-mCherry-(SUMO3)_6_-(PIASx)_10_ plasmid between restriction sites BsrGI and SalI by GenScript (Piscataway, NJ). The plasmid pC1-mCherry-(SUMO3)_5_-(PIASx)_3_-(RK)_5_ was used to generate the plasmid pC1-mCherry-(SUMO3)_5_-(PIASx)_3_-RKRKR. Using the primers in Methods Table 1, the portion of the gene encoding for the (RK)_5_ extension was truncated to RKRKR through the introduction of a stop codon via PCR.

PCR-assembled or ligated products were transformed into XL10-Gold ultracompetent E. coli (Agilent Technologies, Santa Clara, CA). Constructs were purified by the Mini-prep plasmid purification kit, screened by agarose gel electrophoresis and verified by DNA sequencing.

**Methods Table 1.**
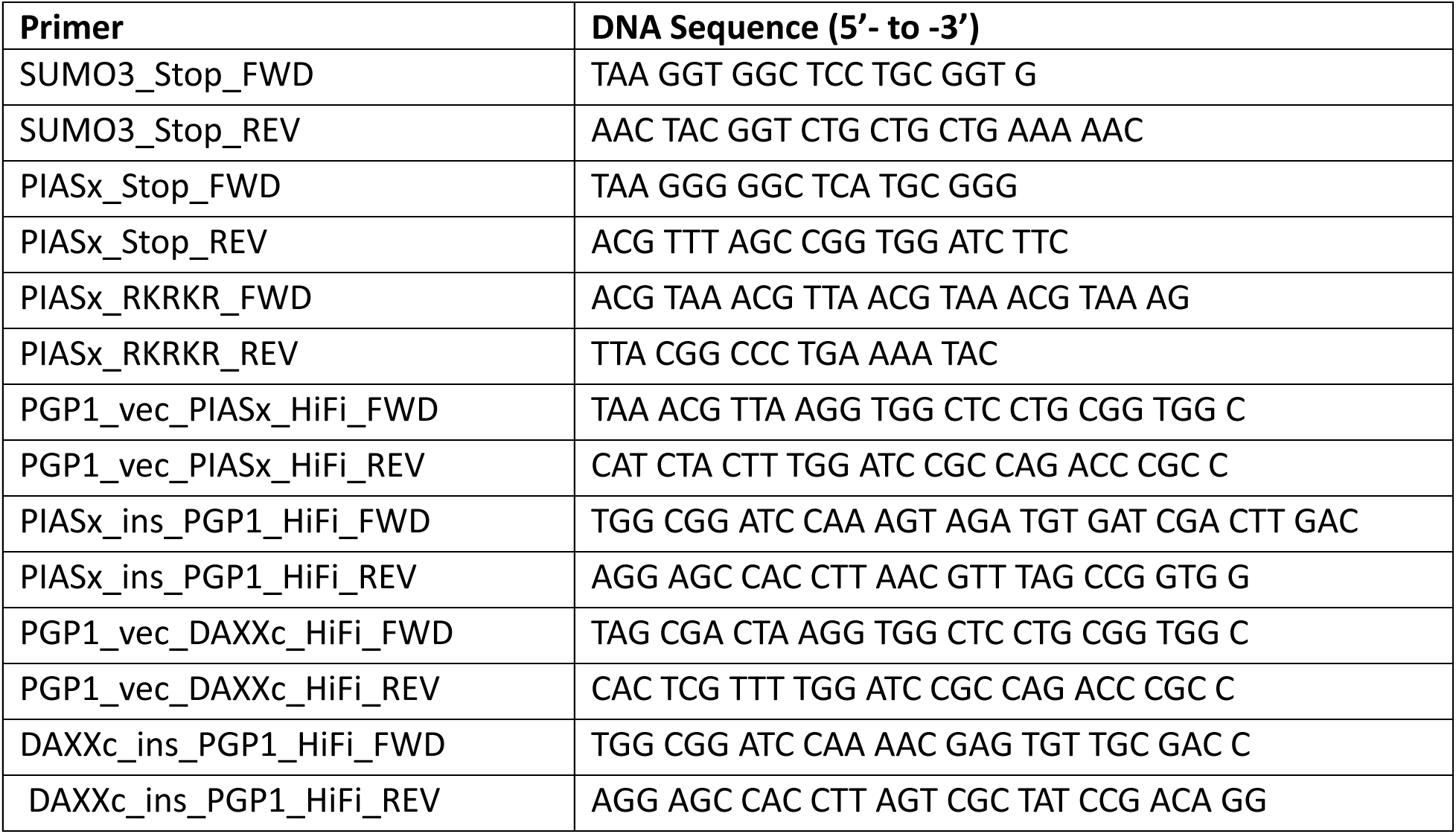
List of ssDNA primers employed in molecular cloning.

**Methods Table 2.**
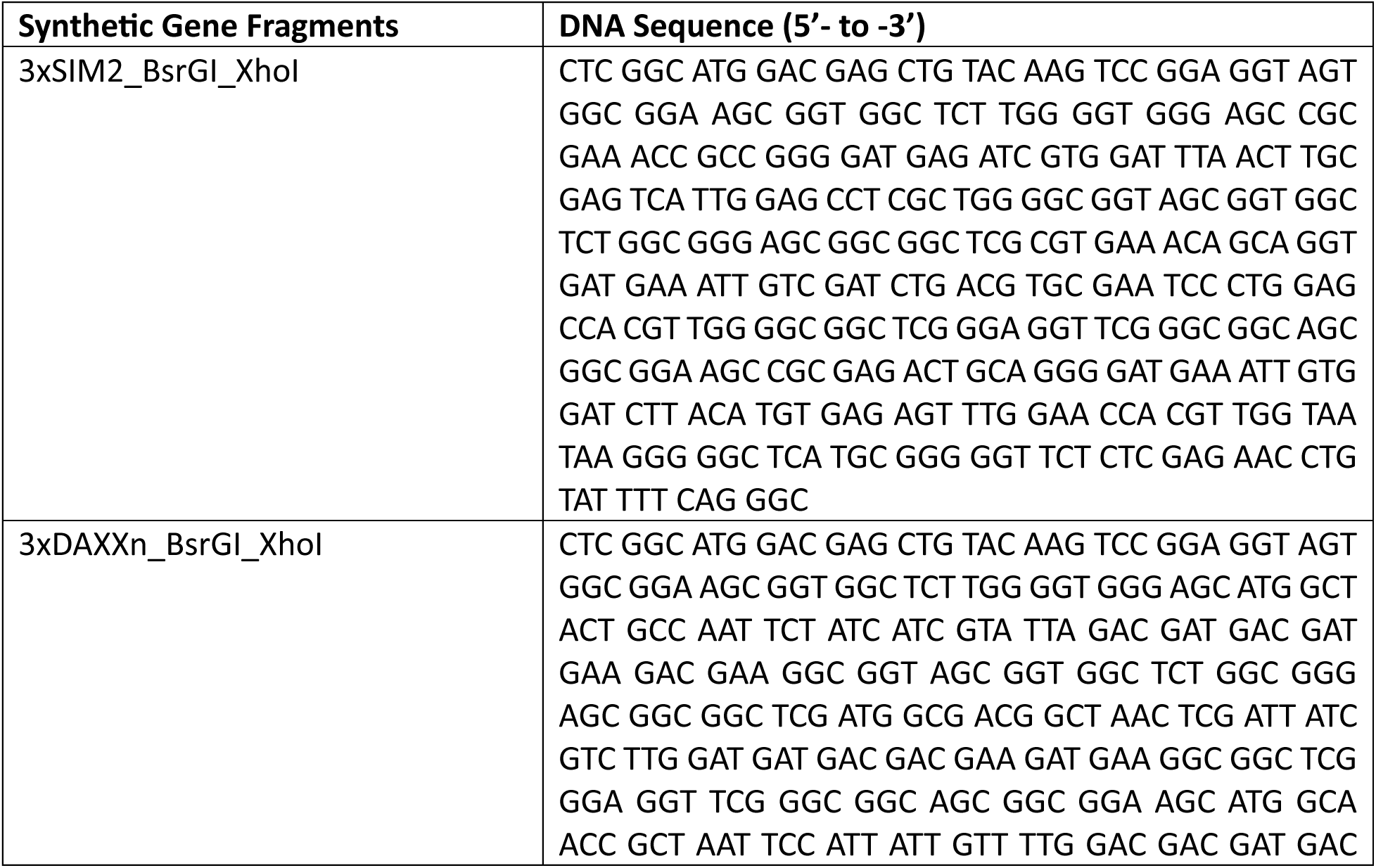
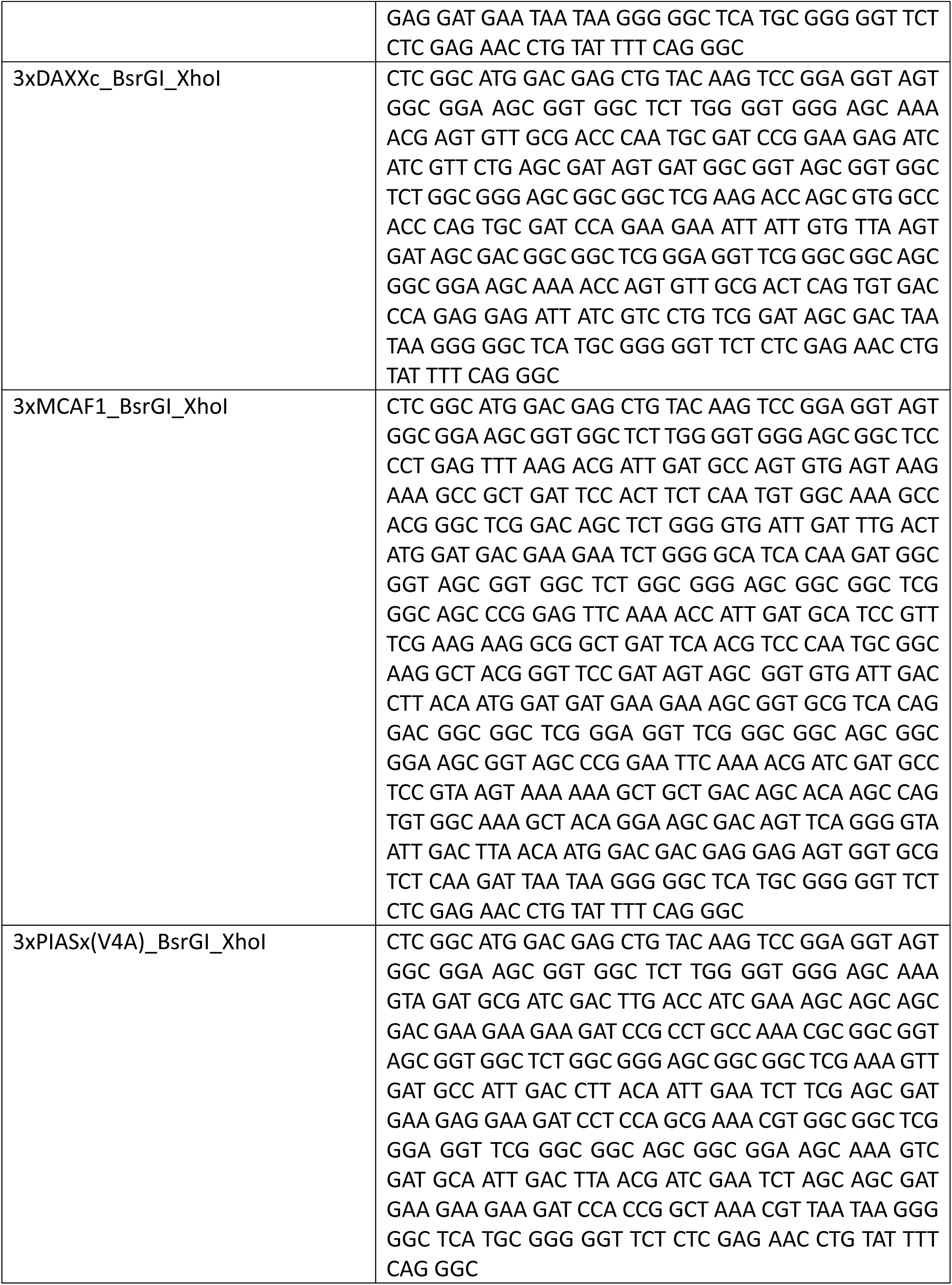
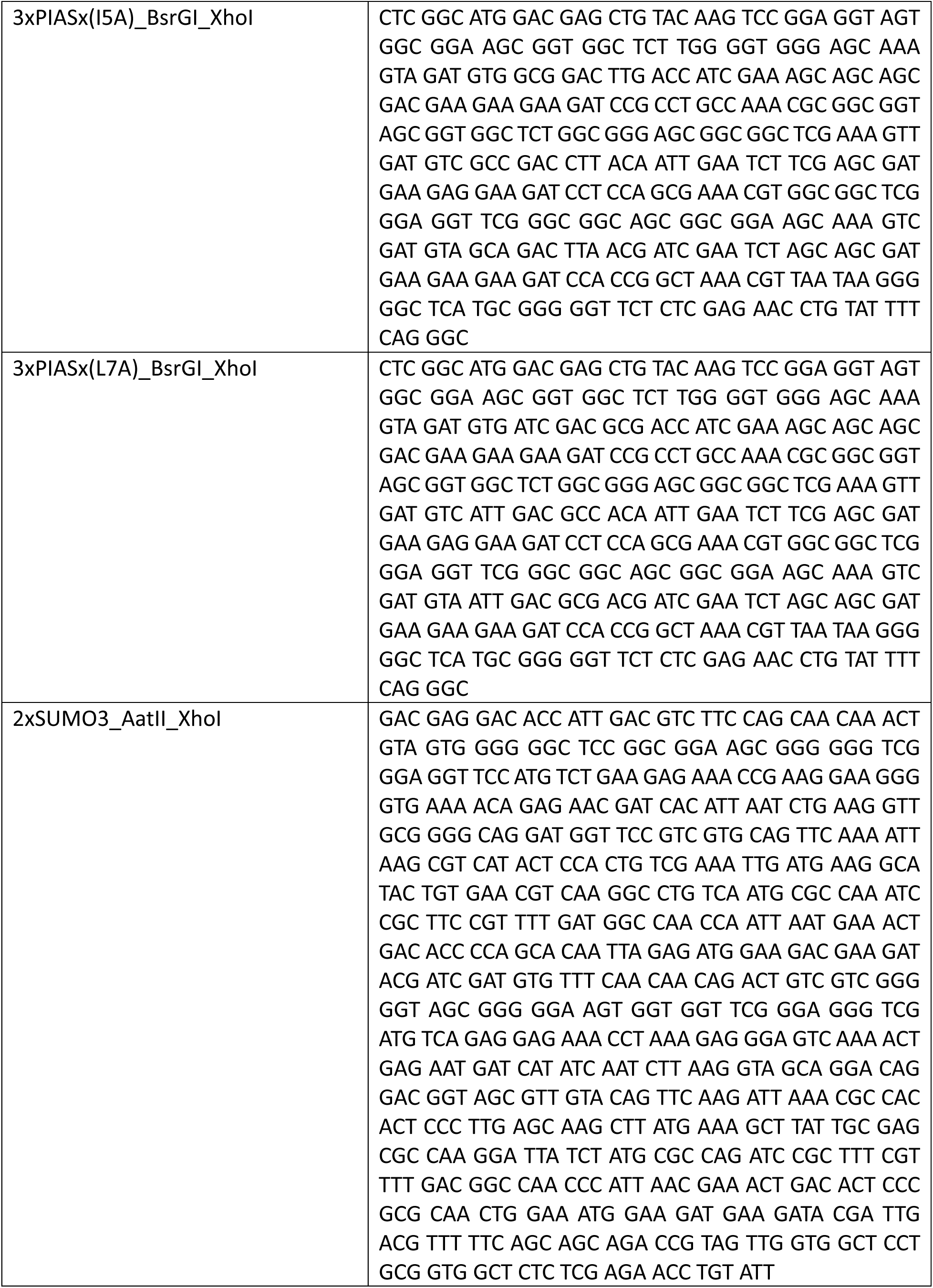
List of synthetic gene fragments employed in molecular cloning.

### Heterologous expression and purification

#### pTEV19-His_10_-mEGFP-(SUMO3)_5_ and pTEV19-His_10_-mEGFP-(SIM)_3_

Two liters of Luria-Bertani (LB) medium, supplemented with 100 µg/mL ampicillin and 34 µg/mL chloramphenicol, were inoculated with E. coli BL21(DE3)pLysS cells transformed with pTEV19-His_10_-mEGFP-(SUMO3)_5_ or pTEV19-His_10_-mEGFP-(SIM)_3_, where the SIM sequence varied. Cells were grown at 37 °C with shaking at 250 rpm to OD_600_ 0.6-0.8. Protein expression was induced with 1 mM IPTG and cultures were shifted to 16 °C for ∼18 h. Cells were harvested by centrifugation at 7,000 × g at 4 °C, and the pellet was either flash frozen in liquid nitrogen and stored at –80 °C or immediately lysed in buffer containing 50 mM sodium phosphate, pH 7.5, 300 mM NaCl, and 1 mM DTT, supplemented with 1 mM iodoacetamide, 2 mM *N*-ethylmaleimide, and 1x cOmplete protease inhibitor cocktail (Roche). Cells were lysed by French press followed by brief sonication for 10 pulses. Lysed cells were clarified by centrifugation at 20,000 × g for 30 min at 4 °C. The supernatant was passed through a 0.45 µm PES filter and loaded onto 5 mL Ni-NTA resin. Protein was incubated on resin for 1 h at 4 °C with gentle nutation after which the flow-through was collected and the resin washed with 5 column volumes (CV) of lysis buffer containing 20 mM imidazole followed by 5CV of 50 mM imidazole, and proteins were eluted with 2.5CV of 500 mM imidazole. Fractions were analyzed by 12% SDS-PAGE, and those containing the target protein were pooled and dialyzed at 4 °C twice against 2 x 4 L of 20 mM Tris, pH 7.5, and 1 mM DTT, first for 3 h then a second time overnight. The dialyzed protein was purified by anion-exchange chromatography on a SOURCE 15Q 4.6/100 PE column (GE Healthcare, Chicago, IL). A linear 0–100% gradient over 40CV was used to increase NaCl concentration up to 1 M NaCl. Buffer A: 20 mM Tris, pH 7.5, and 1 mM DTT. Buffer B: 20 mM Tris, pH 7.5, 1 M NaCl, and 1 mM DTT. Fractions were analyzed by 12% SDS-PAGE then pooled and dialyzed against 2 × 4 L of microscopy buffer (MB) containing 20 mM HEPES, pH 7.5, 150 mM KCl, 1 mM MgCl_2_, 1 mM EGTA, and 1 mM DTT. Protein concentration was determined by absorbance at 488 nm using the mEGFP extinction coefficient of 56,000 M^−1^cm^−1^.^42^ Proteins were aliquoted into single-use volumes to avoid free-thaw cycles, flash frozen in liquid nitrogen, and stored at −80 °C until use.

#### SUMO3(C47S)

Three liters of LB medium supplemented with 100 µg/mL ampicillin were inoculated with E. coli BL21(DE3) transformed with the plasmid pTXB1-SUMO3(1-91)C47S-SUMO3(2-91)C47S-His_6_ containing two head-to-tail fused copies of the SUMO3 gene. The cells were grown at 37 °C with shaking at 250 rpm to OD_600_ 0.6–0.8. Protein expression was induced with 0.8 mM IPTG and cultures were incubated for 3 h at 37 °C with shaking at 250 rpm. Cells were harvested by centrifugation at 7,000 × g for 20 min at 4 °C. The cell pellet was lysed in buffer containing 50 mM sodium phosphate (pH 7.7), 300 mM NaCl, and 5 mM imidazole supplemented with 0.2 mM phenylmethylsulfonyl fluoride (PMSF) and 1x cOmplete protease inhibitor cocktail. Cells were lysed by sonication, and lysates were clarified by centrifugation at 20,000 × g for 30 min at 4 °C. The clarified lysate was passed through a 0.45 µm filter and loaded onto 5 mL Ni-NTA resin pre-equilibrated in lysis buffer. The lysate was incubated with resin for 45 min at 4 °C with gentle nutation. The column was washed with 10CV of lysis buffer, followed by 10CV of lysis buffer containing 20 mM imidazole and then 10CV of 50 mM imidazole. Proteins were eluted with 5 x 3CV of 500 mM imidazole. Fractions were analyzed by 15% SDS-PAGE and fractions containing the diSUMO3 fusion protein were pooled and dialyzed at 4 °C against 2 × 4 L of 50 mM sodium phosphate (pH 7.7), 150 mM NaCl, and 1 mM DTT. To liberate the untagged SUMO3 from the fusion construct, the purified catalytic domain of SENP2 was added to a final concentration of 0.5 µM and the reaction was incubated at 30 °C for 2 h. The reaction mixture was diluted 1:1 with 50 mM sodium phosphate (pH 7.7), 150 mM NaCl to halve the DTT concentration and then loaded onto 5 mL Ni-NTA resin pre-equilibrated with 50 mM sodium phosphate (pH 7.7) and 150 mM NaCl to bind the His-tagged diSUMO3 fusion and His_6_-SENP2. The flow-through contained untagged monomeric SUMO3. The column was washed 4 times with 2CV of reaction buffer lacking DTT and containing 5 mM imidazole. Untagged SUMO3 collected in the flow-through and washes was dialyzed against 2 x 4 L of water. The dialysate was then collected, flash frozen, and lyophilized to dryness. The crude protein was purified via C4 preparative RP-HPLC (5-75% B over 45 min) to yield the pure monomeric SUMO3 protein. Pure fractions were characterized by ESI-MS. SUMO3 calcd. M_avg_ 10,377.6 Da. Observed 10,377.2 ± 1.6 Da.

#### pTEV19-His_10_-PGP1-(PIASx)_3_ and pTEV19-His_10_-PGP1-(DAXXc)_3_

Two liters of LB medium, supplemented with 100 µg/mL ampicillin and 34 µg/mL chloramphenicol, were inoculated with E. coli BL21(DE3)pLysS cells (Novagen, Darmstadt, Germany) transformed with pTEV19-His_10_-PGP1-(PIASx)_3_ or pTEV19-His_10_-PGP1-(DAXXc)_3_. The cells were grown at 37 °C, with shaking at 250 rpm until OD_600_ 0.6–0.8. Protein expression was induced with 1 mM IPTG and cultures were shifted to 16 °C for ∼18 h. Cells were harvested by centrifugation at 7,000 × g at 4 °C, and the pellet was either flash frozen in liquid nitrogen and stored at −80 °C or immediately lysed in buffer containing 50 mM sodium phosphate, pH 7.5, 300 mM NaCl, and 1 mM DTT. Cells were lysed by French press followed by brief sonication for 10 pulses. Lysed cells were clarified by centrifugation at 20,000 × g for 30 min at 4 °C. The supernatant was passed through a 0.45 µm PES filter and loaded onto 5 mL Ni-NTA resin. Protein was incubated on resin for 1 h at 4 °C with gentle nutation. The flow-through was collected and the resin was washed with 5CV of lysis buffer containing 20 mM imidazole followed by 5CV of 50 mM imidazole, and proteins were eluted with 2.5CV of 500 mM imidazole. Fractions were analyzed by 12% SDS-PAGE, pooled and dialyzed at 4 °C twice against 2 x 4 L of 20 mM Tris, pH 7.5, and 1 mM DTT, first for 3 h and then overnight. The dialyzed protein was purified by anion-exchange chromatography on a SOURCE 15Q 4.6/100 PE column (GE Healthcare, Chicago, IL). A linear 0-100% gradient over 40CV was used to increase NaCl concentration up to 1 M NaCl. Buffer A: 20 mM Tris, pH 7.5, and 1 mM DTT. Buffer B: 20 mM Tris, pH 7.5, 1 M NaCl, and 1 mM DTT. Fractions were analyzed by 12% SDS-PAGE, pooled and dialyzed against 2 x 4 L 10 mM Na_2_HPO_4_, 1.8 mM KH_2_PO_4_, 137 mM NaCl, 2.7 mM KCl, and 1 mM DTT, pH 7.5 (PBS). Protein concentration was determined by SDS-PAGE and Coomassie staining relative to BSA standards of known concentration. The cryoprotectant glycerol was added to 10% v/v and samples were flash frozen and stored at −80 °C until use.

### Confocal microscopy imaging of biomolecular condensates

All confocal microscopy was performed on a Leica SP8 X scanning confocal microscope from Leica (Gräfelfing, Germany).

#### *In vitro* condensates

Imaging was performed with a 40x oil-immersion objective. A tunable white-light laser (WLL) was set to 488 nm to excite mEGFP and emission observed between 497-577 nm. For imaging AF568-labeled PGP1-(PIASx)_3_ and PGP1-(DAXXc)_3_ in the presence of mEGFP alone, the WLL was set to 577 nm and emission observed between 582–677 nm. When AF568-labeled PGP1-(PIASx)_3_ and PGP1-(DAXXc)_3_ were imaged before the addition of pGlu-HCA, in anticipation of HCA formation, the emission window was narrowed by the LAS X Dye Assistant software to 582-650 nm. For imaging pGlu-HCA uptake into mEGFP-labeled condensates, the WLL was set to 600 nm and emission observed between 610-690 nm. For HCA imaging, the WLL was set to 670 nm, and emission observed from 710-799 nm. Multi-color acquisitions were performed using line-sequential scanning to minimize crosstalk between fluorophores. The z-focus was set at the glass surface, and the x-y coordinates were centered within each well.

#### *In cellulo* condensates

Imaging in live cells was performed using a 60x oil-immersion objective. A fixed-wavelength 405 nm laser was used to excite Hoechst 33342 fluorescent stain, and emission observed between 410-587 nm. For condensates containing mCherry fusions, the WLL was set to 587 nm to excite mCherry and emission observed between 587-592 nm. For Wheat Germ Agglutinin (WGA)647, the WLL was set to 650 nm to excite AF647 dye, and emission observed between 592-653 nm. Line-sequential scanning was used for multicolor acquisitions to minimize crosstalk. A z-stack was acquired for each condensate, and the middle optical section of largest diameter was used for imaging.

### SUMO-SIM phase diagrams

mEGFP-(SUMO)_5_ and mEGFP-(SIM)_3_ proteins were mixed in an experimental buffer composed of 20 mM HEPES (pH 7.5), 150 mM KCl, 1 mM MgCl_2_, 1 mM EGTA, and 1 mM DTT, at varying concentrations from 20 nM to 140 nM in a final volume of 80 µL. Mixtures were prepared in 96-well plates (Cellvis) surface-treated with bovine serum albumin (BSA) and incubated for 20-24 h at 25 °C. Wells were imaged at 40x magnification and condensate formation was scored as either present or absent to delineate the phase boundary for each SUMO-SIM pair.

### Stability of condensates in KCl, ATP, Urea and Gn-HCl

BMC mixtures were first prepared in 96-well plates (Cellvis) surface-treated with BSA by mixing 2 µM mEGFP-(SUMO)_5_ and 2 µM of the respective mEGFP-(SIM)_3_ in experimental buffer (EB) containing 20 mM HEPES (pH 7.5), 150 mM KCl, 1 mM MgCl_2_, 1 mM EGTA, and 1 mM DTT and incubating for 20-24 h at room temperature prior to imaging. To test the effect of salt, hydrotrope and chaotropes on the stability of condensates, a stock of EB containing the concentrated reagent was added to preformed condensates to reach a maximum of either 3 M KCl, 50 mM ATP, or 3 M urea/Gn-HCl in a total volume of 80 µL. Condensates were imaged after incubation for an additional 2 h at 25 °C.

### Fluorescence recovery after photobleaching (FRAP)

A Leica SP8X scanning confocal microscope with a 40x oil-immersion objective was used with the FRAP module in Leica LAS X software. Pre-bleach images were acquired for baseline normalization. A zoom factor was applied to center an individual condensate in the field-of-view, and the region of interest (ROI) was defined as approximately 50% of the visible droplet area. The ROI was photobleached with a 488 nm argon laser at 100% power. Recovery was monitored by time-lapse image acquisition until fluorescence intensity plateaued, and intensities were background-subtracted and normalized to the pre-bleach mean.

For FRAP in living cells, they were first transfected with the respective mCherry-(SUMO)_5_-(SIM)_3_ plasmid and counterstained with WGA647 and Hoechst 33342 to delineate the cell membrane and nucleus, respectively. A Leica SP8X scanning confocal microscope with a 63x oil-immersion objective was used with the FRAP module in Leica LAS X software. FRAP acquisition settings were identical to those used for in vitro assays. The ROI was photobleached with a 587 nm WLL at 100% power. Recovery was monitored by time-lapse image acquisition until fluorescence intensity reached a plateau, and intensities were background-subtracted and normalized to the pre-bleach mean.

### Statistical analysis of FRAP data

Data were analyzed in ImageJ/Fiji. Fluorescence intensities were background-subtracted and normalized to the pre-bleach mean of each ROI. To account for acquisition-induced photobleaching during the post-bleach recovery phase, intensities were corrected using an unbleached reference ROI. The resulting normalized intensities, *I(t)*, were fit by nonlinear least-squares to a single-exponential model *I(t) = I*_∞_ *+ (I*_0_ *−I*_∞_*)e^−kt^*, where *I*_0_ is the intensity in the bleached ROI immediately after bleaching, *I*_∞_ is the plateau intensity after fluorescence recovery, and *k* is the apparent first-order rate constant. The characteristic time constant *τ* was defined as 1/*k*. Half-times, t_1/2_, were calculated as (ln 2)/*k*.^43^

### Coarse-grained molecular dynamics simulations

All coarse-grained (CG) simulations used a one-bead-per-residue model with the Urry hydropathy scale.^23^ The initial structure of (SUMO)₅ was obtained from Boltz2 structure prediction.^44^ Folded SUMO domains were treated as rigid bodies and linkers as fully flexible chains. To validate the parametrization, single-chain simulations of (SUMO)₅ were performed in LAMMPS and coexistence simulations were conducted using HOOMD-Blue^45^ (version 4.0), both using NVT Langevin thermostat settings (γ = mAA/ρ with ρ = 1000 ps, timestep = 10 fs). Using the original global interactions (ε = 0.2 kcal/mol), (SUMO)₅ exhibited over-collapse in single-chain simulation and underwent phase separation in coexistence simulation, inconsistent with experimental observations. Reducing intra-domain SUMO interaction strengths by 30% yielded expanded conformational ensembles (radius of gyration Rg and end-to-end distance Ree) and eliminated artifactual phase separation (Supplementary Fig. S9a-b). This re-parameterization strategy, which compensates over-collapse artifacts arising from rigid-body constraints, has been successfully applied to full-length TDP-43 simulations to achieve agreement with in vitro phase separation observations.^46^ The 30% reduction was applied exclusively to intra-domain SUMO residue pairs; linker interactions retained the original interaction.

All (SIM)₃ constructs (PIASx, MCAF1, DAXXc, DAXXn, SIM2, PIASx-RKRKR, and PIASx-(RK)₅) were modeled as fully flexible chains. Systems were initialized in elongated rectangular boxes at a 5:3 (SUMO)₅:(SIM)₃ molar ratio, with box dimensions chosen to minimize finite-size effects following established protocols.^22,47,48^ Because SUMO-SIM phase separation involves both specific hydrophobic and non-specific electrostatic interactions that vary across SIM sequences, phase behavior was mapped as a function of a global SUMO-SIM interaction scaling parameter ε (range: 0.20–0.39 kcal/mol). This scaling was applied exclusively to inter-molecular SUMO-domain–SIM interactions; all linker interactions retained their original Urry parameters. For each ε value, 300 ns simulations were performed to construct the phase diagram. At the representative value of ε = 0.33, extended simulations of 5 μs were carried out, with the first 1 μs discarded as equilibration for all analyses. Density profiles along the box axis were computed to identify coexisting phases and estimate dilute-phase concentrations. Inter-molecular contact maps were calculated using a distance cutoff of 1.5× the arithmetic mean of residue van der Waals radii. Snapshots were rendered in VMD.^49^

### All-atom molecular dynamics simulations

Initial structures for each SUMO-SIM complex were obtained from Boltz2 structure prediction. Prediction confidence scores for all five SIM constructs (M1, Px, Dn, S2, Dc; n = 25 models each, Supplementary Fig. S9c-d). System topologies were prepared using GROMACS 2021.6,^50^ with the Amber99SBws-STQ’ force field^51^ (https://bitbucket.org/jeetain/all-atom_ff_refinements). Each complex was solvated with TIP4P/2005 water in an octahedral box with a minimum 1.5 nm buffer between the protein and box edge. Energy minimization was performed using the steepest descent algorithm, first in vacuum and again after solvation. Na⁺ and Cl⁻ ions were added to a physiological concentration of 150 mM with additional counterions for charge neutrality, using the improved ion parameters of Lou and Roux.^52^ The system was equilibrated sequentially for 100 ps in NVT (Nosé–Hoover thermostat,^53^ coupling constant 1.0 ps, 300 K) and 100 ps in NPT (Berendsen barostat,^54^ isotropic coupling constant 5.0 ps, 1 bar). Production simulations were run in Amber22. GROMACS input files were converted to Amber format using PARMED^55^ with hydrogen mass repartitioning to 1.5 amu, enabling a 4 fs timestep. After conversion, the system was energy-minimized and then heated over 5 ns from 100 K to 300 K (with protein positions restrained). Two successive NVT equilibrations at 300 K were performed (5 ns with backbone restraints, then 5 ns unrestrained), followed by 1 ns NPT equilibration (Monte Carlo barostat, 1.0 ps coupling, 1 bar). Temperature was controlled via Langevin dynamics (friction coefficient 1.0 ps⁻¹); hydrogen bonds were constrained with SHAKE.^56^ Short-range non-bonded interactions used a 0.9 nm cutoff; long-range electrostatics were treated with Particle Mesh Ewald (PME).^57,58^ Ten independent 1 μs NPT production replicas were then collected per system.

### Bound-state lifetime (τ) analysis

Trajectories were analyzed frame-by-frame using MDAnalysis 2.5.0.^59^ Because each replica was initiated from a Boltz2-predicted bound complex and preceded by a final 1 ns unrestrained equilibration, the initial bound interval was retained as a complete event in the primary analysis. For each frame, three descriptors were computed: the fraction of native contacts (Q_native_), the total number of intermolecular heavy-atom contacts, and the SUMO–peptide centroid distance. Native contacts were defined as all SUMO-SIM heavy-atom pairs within 4.5 Å in the reference bound structure. A frame was classified as bound if Q_native_ ≥ 0.3, or if at least 10 intermolecular heavy-atom contacts were present with a centroid distance ≤ 10 Å. This binary bound/unbound time series was segmented into contiguous intervals; bound lifetimes correspond to the elapsed time of each continuous bound segment. Terminal bound intervals were treated as right-censored observations. As a robustness check, analyses in which the initial interval was censored or excluded were also performed; these did not affect the qualitative trends across peptides. Bound-state survival curves were estimated using the Kaplan–Meier estimator.^60^ Mean bound lifetimes τ and standard errors were calculated from uncensored events only and visualized as violin plots. Peptides are displayed throughout as M1, Px, Dn, S2, and Dc (corresponding to MCAF1, PIASx, DAXXn, SIM2, and DAXXc, respectively).

### Sedimentation analysis of complex condensates

To obtain ternary condensates, 2 µM each of mEGFP-(SUMO)_5_ and mEGFP-(DAXXc)_3_ were mixed in phase separation buffer (PSB) composed of 20 mM HEPES pH 7.5, 150 mM KCl, 1 mM MgCl_2_, 1 mM EGTA, and 1 mM DTT, in a final volume of 500 µL in a 1.5 mL microcentrifuge tube. The mixture was gently mixed by pipetting and incubated at 25 °C for 12 h. mEGFP-(MCAF1)_3_ was then added to a final concentration of 2 µM and the mixture was further mixed by pipetting then incubated for an additional 12 h at 25 °C. For quaternary condensates, mEGFP-(SUMO)_5_ and mEGFP-(DAXXc)_3_ were first mixed and incubated at 25 °C for 12 h. mEGFP-(MCAF1)_3_ was then added to a final concentration of 2 µM, the mixture gently mixed by pipetting, and incubated for an additional 12 h at 25 °C. mEGFP-(PIASx)_3_-(RK)_5_ was subsequently added to a final concentration of 2 µM and the mixture allowed to equilibrate for 12 h at 25 °C. The mixtures of condensates were centrifuged at 17,200 x g for 30 min at 25 °C to separate the dense and dilute phases. The supernatant dilute phase was carefully pipetted away from the dense phase at the bottom of the tube, and the dense phase subsequently resuspended in a volume of PSB equal to a third of the dilute phase. Dilute and dense-phase fractions were diluted in 1x Laemmli buffer, boiled for 2 min, and analyzed by 12% and 10% SDS-PAGE for tertiary and quaternary mixtures, respectively.

### Reversed Phase High Performance Liquid Chromatography (RP-HPLC)

Analytical RP-HPLC was performed on an Agilent 1260 Infinity II quaternary system with either a Grace-Vydac C18 column (5 µm, 4.6 × 150 mm) or a Waters C4 column (5 µm, 4.6 × 150 mm) at 1.0 mL/min. A typical analytical gradient was 0-73% B over 30 min. Semi-preparative and preparative RP-HPLC were performed on an Agilent 1260 Infinity II preparative HPLC with either a Zorbax StableBond preparative C18 column (21.2 × 250 mm), a Hypersil Gold preparative C4 column (25 × 250 mm), a Zorbax StableBond semi-preparative C18 column (9.4 × 250 mm), or a Waters semi-preparative C4 column (10 × 250 mm). Preparative runs were performed at 20 mL/min and semi-preparative runs at 4 mL/min. Elution was monitored by UV-vis absorbance at 214 and 280 nm.

### Electrospray Ionization Mass Spectrometry (ESI-MS)

Peptide and protein mass spectra were acquired by direct infusion electrospray ionization on a Bruker Esquire ion trap mass spectrometer (Billerica, MA) or on a Finnigan LTQ ion trap mass spectrometer (Thermo, Waltham, MA), operated in positive ion mode.

### Synthesis, purification, and characterization of SIM peptides

SIM peptides were generated by solid-phase peptide synthesis (SPPS) using the Fmoc-protecting group strategy on either a CEM Liberty Blue 1.0 or CEM Liberty PRIME 2.0 microwave-assisted peptide synthesizer. When needed, the initial high resin loading of 0.6 to 0.8 meq g^−1^ was reduced to about 0.2 meq g^−1^ to limit aggregation of the elongating peptide chain and preclude deletion products arising from occlusion of the N-terminal amine. Crude peptides were precipitated in ice-cold diethyl ether, pelleted by centrifugation, dissolved in Buffer A with minimal amounts of Buffer B as needed, and purified by C18 preparative RP-HPLC. Pure peptide fractions were pooled and analyzed by ESI-MS. Purity was further confirmed by observing peak shapes during analytical RP-HPLC on a 0-73% B gradient. When applicable, acetyl capping of the uncoupled N-terminus was performed after difficult Asp couplings to avoid the formation of inseparable Asp-deletion peptides.

Synthesis of PIASx peptide: Ac-KVDVIDLTIESSSDEEEDPPAKR-C(O)NH₂

Rink amide (MBHA) resin (0.667 meq g^−1^) was swollen in 1:1 DMF/DCM and deprotected with 20% (v/v) piperidine in DMF for 30 min. To reduce resin-loading, the third residue (Ala) was coupled as a 1:1 mixture of Fmoc-Ala-OH and Boc-Ala-OH (0.8 mmol each) with Oxyma (0.8 mmol) and DIC (0.8 mmol) for 60 min at 25 °C. Reduced loading was verified by measuring the amount of dibenzofulvene–piperidine adduct formed from its absorbance at 301 nm after treatment with 20% (v/v) piperidine in DMF. Chain elongation proceeded on the Liberty Blue using 5 equivalents of each Fmoc-amino acid relative to the reduced loading. Fmoc deprotection used 5% (w/v) piperazine with 0.05 M HOBt in DMF for 3 min at 75 °C. Standard couplings conditions were 10 min at 75 °C with Fmoc-AA (0.5 mmol), DIC (0.49 mmol), Oxyma (0.49 mmol), and DIEA (0.98 mmol) in DMF; additional double couplings were used for Fmoc-Arg(Pbf)-OH, Fmoc-Thr(OtBu)-OH, Fmoc-Val-OH, Fmoc-Lys(Boc)-OH, Fmoc-Asp(OtBu)-OH, and Fmoc-Leu-OH as needed. The N terminus was capped with acetic anhydride (0.5 mmol) and DIEA (0.5 mmol) for 10-15 min at 25 °C. Cleavage of the final peptide from the resin was undertaken with TFA/triisopropylsilane/H₂O, 90:5:5 (v/v), for 2 h at 25 °C. The ether-precipitated peptide was dissolved in Buffer A, filtered through a 0.45 µm syringe-filter, lyophilized, and purified by C18 preparative RP-HPLC (17-48%B over 40 min). Pooled fractions were analyzed by ESI-MS. Final purity was confirmed by C18 analytical RP-HPLC and by ESI-MS. Calculated M_avg_ = 2,613.3 Da; found 2,613.3 ± 0.4 Da (Supplementary Fig. S10).

Synthesis of MCAF1 peptide: Ac-GSPEFKTIDASVSKKAADSTSQCGKATGSDSSGVIDLTMDDEESGASQ D-C(O)NH₂

Rink amide (MBHA) resin (0.667 meq g^−1^) was swollen in 1:1 DMF/DCM and deprotected with 20% piperidine in DMF (v/v). Resin-loading was reduced using a mixture of Boc-Ala-OH and Fmoc-Asp(OtBu)-OH (0.24 mmol). Chain elongation was undertaken on a CEM Liberty PRIME 2.0 synthesizer with a wash-free method.^61^ Additional double couplings were applied for Fmoc-Ile-OH, Fmoc-Thr(OtBu)-OH, Fmoc-Val-OH, Fmoc-Lys(Boc)-OH, Fmoc-Ala-OH, Fmoc-Leu-OH, Fmoc-Gly-OH, Fmoc-Cys(Trt)-OH, Fmoc-Ser(OtBu)-OH, and Fmoc-Gln(Trt)-OH. Peptide cleavage from the resin used TFA/triisopropylsilane/H₂O/anisole, 85/5/5/5 (v/v) and it was purified by C18 preparative RP-HPLC (19–43%B over 45 min). Final peptide purity was confirmed by C18 analytical RP-HPLC and by ESI-MS. Calculated M_avg_ = 4,953.2 Da; found 4,953.1 ± 0.1 Da (Supplementary Fig. S10).

Synthesis of DAXXn peptide: Ac-MATANSIIVLDDDDEDE-C(O)NH₂

Synthesized identically to the MCAF1 synthesis. Cleavage cocktail contained TFA/triisopropylsilane/H₂O/anisole/thioanisole, 85/5/5/2.5/2.5 (v/v). Purification undertaken by C18 preparative RP-HPLC (17-48% B gradient over 40 min). Pure fractions were checked by C18 analytical RP-HPLC on a gradient of 0-73% over 30 min and their identity was confirmed by ESI-MS. Calculated M_avg_ = 1,906.9 Da; found 1,907.2 ± 0.3 Da (Supplementary Fig. S10).

Synthesis of SIM2 peptide: Ac-RETAGDEIVDLTCESLEPRW-C(O)NH₂

Synthesized identically to the MCAF1 synthesis, with additional double couplings for Fmoc-Glu(OtBu)-OH and Fmoc-Arg(Pbf)-OH. Cleavage cocktail contained TFA/triisopropylsilane/H₂O/anisole/thioanisole, 85/5/5/2.5/2.5 (v/v). The crude peptide was purified by C18 preparative RP-HPLC (25-55% B over 40 min). Final peptide purity was confirmed by analytical C18 RP-HPLC on a gradient of 0-73% B over 30 min and by ESI-MS. Calculated M_avg_ = 2,360.5 Da; found 2,361.1 ± 0.4 Da (Supplementary Fig. S10).

Synthesis of DAXXc peptide: Ac-KTSVATQCDPEEIIVLSDSD-C(O)NH₂

Fmoc-Asp-Wang resin (0.57 meq g^−1^) was swollen in DMF for 20–30 min and deprotected with 20% (v/v) piperidine in DMF. Loading was reduced as in PIASx synthesis using Boc-Ser(OtBu)-OH (0.4 mmol). Elongation used a CEM Liberty PRIME 2.0 with the wash-free method and 5 equivalents per coupling. After elongation, capping and cleavage followed the method established for PIASx synthesis. Purification was undertaken by C18 preparative RP-HPLC (20-44% B over 40 min). Final pure fractions were checked by C18 RP-HPLC on a gradient of 0-73% B over 30 min and their identity was confirmed by ESI-MS. Calculated M_avg_ = 2,192.3 Da; found 2,192.2 ± 0.1 Da (Supplementary Fig. S10).

Synthesis of PIASx(V4A) peptide: Ac-KVDAIDLTIESSSDEEEDPPAKR-C(O)NH₂

Synthesized and purified identically to the wild-type PIASx peptide. Calculated M_avg_ = 2,585.7 Da; found 2,585.4 ± 0.4 Da (Supplementary Fig. S10).

Synthesis of MCAF1(V34A) peptide: Ac-GSPEFKTIDASVSKKAADSTSQCGKATGSDSSGAIDL TMDDEESGASQD-C(O)NH₂

Synthesized and purified identically to the wild-type MCAF1 peptide. Calculated M_avg_ = 4,925.1 Da; found 4,925.8 ± 0.9 Da (Supplementary Fig. S10).

### Synthesis of PIASX-(RK)_5_ peptide by native chemical ligation (NCL)

Synthesis of PIASx peptidyl hydrazide **1**: Ac-KVDVIDLTIESSSDEEEDPPAKRGGS-C(O)NHNH₂

2-Chlorotrityl hydrazide resin was prepared by reacting 2-chlorotrityl chloride resin (1.5–2.0 meq g⁻¹) with 10% (v/v) hydrazine in DMF at 25 °C for 30 min, repeated once with fresh hydrazine solution. Residual sites were capped with 10% (v/v) methanol in DMF for 10 min. SPPS was identical to the wild-type PIASx peptide, with reduced loading using Boc-Gly-OH (0.2 mmol). Calculated M_avg_ = 2,829.9 Da; found 2,829.6 ± 0.4Da (Extended Fig. 7).

Synthesis of PIASx N-terminal cysteine peptide **2**: H₂N-CGGSLENLYFQGRKRKRKRKRKG-C(O)NH₂ Synthesized identically to the wild-type PIASx peptide with additional double couplings for Fmoc-Glu(OtBu)-OH, Fmoc-Asn(Trt)-OH, and Fmoc-Cys(Trt)-OH. Calculated M_avg_ = 2,765.3 Da; found 2,765.0 ± 0.1 Da (Extended Fig. 7).

Finally, peptide **1** was dissolved in 0.2 M sodium phosphate, pH 3.0, with 6 M guanidine HCl. Sodium nitrite (10 equivalents) was added at −20 °C for 20 min to form the peptidyl azide. Peptide **2** (1.5 equivalents) was dissolved in the same buffer and mixed with 100 equivalents of 4-mercaptophenylacetic acid (MPAA). After conversion of **1** to the peptidyl azide, the solution of **2**+MPAA was added to initiate native chemical ligation at 25 °C. Aliquots were removed hourly, quenched with phosphate buffer and ∼80 equivalents TCEP, and analyzed by RP-HPLC and ESI-MS. Upon completion, the reaction was quenched with ∼80 equivalents TCEP and purified by C18 RP-HPLC (20–60% B over 45 min). Product identity was confirmed by ESI-MS. Calculated M_avg_ = 5,563.2 Da; found 5,564.3 ± 1.0 Da (Extended Fig. 7).

### Measuring SUMO-SIM affinity by ITC

ITC was performed in a buffer containing either 20 mM Tris pH 7.5 or 20 mM MES pH 6.5 and 150 mM KCl, on a MicroCal PEAQ-ITC (Malvern Panalytical, Westborough, MA). The titration program consisted of one 0.4 µL priming injection followed by 18 x 2 µL injections every 150 s at 25 °C. SUMO was dialyzed into the appropriate buffer and the post-dialysis buffer was used to dissolve each SIM peptide. Peptide solutions were adjusted to match the protein pH. Following instrument manufacturer protocols, the sample cell and syringe were washed with 20% Contrad70 (Decon Labs, PA), rinsed with water, and dried with methanol. The sample cell was then rinsed four times with Milli-Q® water followed by thrice with the working buffer. Three hundred microliters of SUMO solution were loaded into the ITC cell and 40 µL of concentrated SIM solution into the syringe. Heats of injection were integrated to generate binding isotherms, and ΔH values were fit with the instrument manufacturers binding model in MicroCal PEAQ-ITC Analysis Software to obtain binding affinities (Supplementary Fig. S11).

For the peptides SIM2, PIASx(V4A) and MCAF1(V34A), Malvern Panalytical’s concatenation workflow was used to extend the isotherm to higher ligand-to-protein ratios without increasing peptide concentration in the syringe. This was necessitated by the lower solubility of these peptides in buffer at pH 7.5. Two sequential titrations were performed per replicate measurement. After the first complete set of titrations, the syringe was refilled with peptide solution while the same protein sample remained in the cell. The two binding isotherms were concatenated in the MicroCal Concat ITC software and analyzed as a single isotherm (Supplementary Fig. S11).

### mCherry-(SUMO)_5_-(SIM)_3_ constructs for live-cell imaging

pC1-mCherry-(SUMO)_5_-(SIM)_3_ DNA for transient transfection was prepared by Miraprep of *E. coli* cells using a Qiagen (Germantown, MD) DNA miniprep kit.^62^ Briefly, transformed E. coli XL10-Gold cells (Agilent, Santa Clara, CA) were grown overnight at 37 °C in 50 mL LB medium supplemented with ampicillin (100 µg/mL). Cells were collected by centrifugation and resuspended in Buffer P1 supplemented with fresh RNase. After alkaline lysis and neutralization, the supernatant was cleared by centrifugation. The supernatant was diluted with an equal volume of 96% (v/v) ethanol prior to loading onto four Qiagen miniprep spin columns. DNA was washed and eluted according to the Qiagen protocol. Purity of the eluted DNA was checked by measuring the A_260_/A_280_ ratio on a NanoDrop 2000c spectrophotometer and by agarose gel electrophoresis. The correct gene sequences for all constructs were confirmed by DNA sequencing prior to transfection of human cells.

### Transient transfection of human cells

HeLa, HEK293T, and H1299 cells were maintained in DMEM (HeLa, HEK293T) or RPMI 1640 (H1299) supplemented with 10% fetal bovine serum at 37 °C in a humidified incubator with 5% CO₂. HeLa, HEK293T, and H1299 cells were cultured to ∼60% confluence before transient transfection with pC1-mCherry-(SUMO)_5_-(SIM)_3_ plasmids. The growth medium was replaced at least 1 h before transfection. Cells were transfected using Lipofectamine 3000 (Invitrogen) at 0.26 µg DNA per well in 8-well µ-Slides (Ibidi), or 0.13 µg for the PIASx-(RK)_5_ plasmid, and the medium was replaced 24 h later. Cells were incubated in transfection medium for an additional 24 h at 37°C in a humidified incubator with 5% CO₂ prior to counterstaining and confocal microscopy.

### Counterstaining of human cells for confocal microscopy

HeLa, HEK293T, and H1299 cells were washed 3x with warm DPBS prior to counterstaining with Hoechst 33342 (Thermo Scientific) and wheat germ agglutinin conjugated to Alexa Fluor 647 (WGA647, Invitrogen). A mixture of Hoechst 33342 at 1 µg/mL and WGA647 at 5 µg/mL was prepared in warm DPBS. This mixture was added to cells and incubated for 10 min at room temperature in the dark. Following this, cells were washed 3x with warm DPBS to remove residual stain and returned to warm growth medium. Confocal microscopy was performed immediately after counterstaining.

### Synthesis of hemicyanine amine (HCA)

HCA was synthesized as described with minor modifications.^20^ Briefly, 3-nitrophenol (461 mg, 3.75 mmol) and K_2_CO_3_ (458 mg, 3.75 mmol) were suspended in 20 mL acetonitrile under an argon atmosphere. IR-780 (1.0 g, 1.33 mmol) was added, and the reaction mixture was stirred for 4 h. The solvent was evaporated and the residue was dissolved in DCM, washed thrice with H_2_O and dried over Na_2_SO_4_. Organic layers were evaporated and the residue was dissolved in 40 mL MeOH and maintained under argon. SnCl_2_ (6.0 g, 30 mmol) in concentrated HCl (5 mL) was added via syringe and the reaction mixture gently refluxed overnight. The reaction was quenched by the addition of saturated Na_2_CO_3_ and filtered. The filtrate was diluted with DCM, washed thrice with H_2_O and dried over Na_2_SO_4_. After solvent evaporation, the crude residue was purified by silica gel chromatography using a gradient of 2-20% (v/v) MeOH/DCM to afford HCA as a green solid (162 mg, 26%).

### Synthesis of L-pyroglutamyl hemicyanine (pGlu-HCA)

Boc-L-pyroglutamic acid (71 mg, 0.3 mmol) and HATU (230 mg, 0.6 mmol) were suspended in 15 mL of dry DCM at 0 °C under an argon atmosphere. DIPEA (196 μL, 0.9 mmol) was added via syringe and the reaction mixture was stirred for 30 min on ice. HCA (162 mg, 0.3 mmol) dissolved in 2 mL of dry DCM was then added via syringe, and the reaction was brought to room temperature and stirred vigorously for 36 h under an argon atmosphere. The reaction mixture was then diluted with DCM, washed thrice with H_2_O and dried over Na_2_SO_4_ before solvent evaporation. The residual solid was suspended in DCM (10 mL) at 0 °C and TFA (5 mL) was added dropwise to remove the Boc-protecting group. The reaction mixture was then brought to room temperature and stirred for 3 h. After solvent evaporation, the crude residue was resuspended in MeOH and purified by C18 preparative RP-HPLC on a gradient of 30-90% B over 40 min. Fractions deemed pure by C18 analytical RP-HPLC and ESI-MS were combined and lyophilized to yield pGlu-HCA (12.4 mg, 6%) as a violet solid (Supplementary Figure S7).

### AlexaFluor labeling of PGP1-(PIASx)_3_ and PGP1-(DAXXc)_3_ enzymes

PGP1-(PIASx)_3_ and PGP1-(DAXXc)_3_ were labeled on reactive amines with Alexa Fluor 568 maleimide following the manufacturer’s protocol (Invitrogen) to obtain AF568-PGP1-(PIASx)_3_ and AF568-PGP1-(DAXXc)_3_. Labeled proteins were purified using the Zeba Dye and Biotin removal gel filtration column supplied with the kit. Protein concentrations were determined by absorbance at 577 nm and 280 nm using a NanoDrop 2000c UV-Vis spectrophotometer (ThermoFisher Scientific, Waltham, MA). Concentrations were calculated using the equation below (ε_280_ = 25,900 M⁻¹ cm⁻¹ for both enzymes):

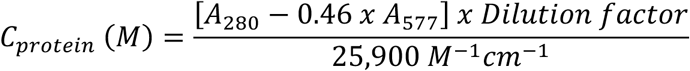

Where A_280_ and A_577_ denote absorbance at 280 nm and 577 nm, respectively.

### Michaelis-Menten kinetics of AF568-PGP1-(PIASx)_3_ and AF568-PGP1-(DAXXc)_3_

Flat-bottom 96-well plates were passivated with 0.1 mg/mL BSA in 10 mM Na_2_HPO_4_, 1.8 mM KH_2_PO_4_, 137 mM NaCl, 2.7 mM KCl, pH 7.5 (PBS) for 1 h at room temperature. Wells were washed five times with H_2_O followed by thrice with PBS to remove residual BSA. 200 microliter reactions were prepared in PBS in BSA-passivated wells at 37 °C. pGlu-HCA concentration was held at 1 µM, 2.66 µM, 4 µM, 7 µM, 13 µM or 19 µM per well. Reactions were initiated by adding either AF568-PGP1-(PIASx)_3_ or AF568-PGP1-(DAXXc)_3_ to a final concentration of 200 nM and monitored in a BioTek Synergy 2 microplate reader (Winooski, VT) at λ_ex_/λ_em_ = 670/700 nm, recording every 30 s for 2 h. Initial velocities (v_0_) were computed from the linear portion of the first 5 min of each progress curve using linear regression analysis. Background slopes corresponding to either pGlu-HCA substrate with no enzyme present and enzyme only minus substrate were subtracted from the observed slopes. Initial velocity, v_0_ values were plotted versus substrate concentration and fit by nonlinear least squares to the Michaelis–Menten equation, *v*_0_*= (V*_max_*[S])(K*_M_ *+ [S])*, in R (RStudio; packages: ggplot2 and broom) to obtain K_m_ and V_max_. The turnover number k_cat_ was calculated as V_max_/[E]_T_ using the enzyme concentration.

### Enzyme kinetics in mixtures of dense and dilute phases

Flat bottomed 96-well plates were passivated with 0.1 mg/mL BSA in PBS for 1 h at 25 °C. Wells were washed five times with H_2_O followed by three washes with PBS to remove residual BSA. 200 microliter enzyme assays were assembled in BSA-passivated wells in PBS at 37 °C. For condensate-containing samples, preformed mEGFP-(SUMO)_5_:mEGFP-(PIASx)_3_ condensates or mEGFP-(SUMO)_5_:mEGFP-(DAXXc)_3_ condensates were diluted into PBS immediately before use. Aliquots of condensate suspensions were added to the passivated wells, followed by AF568-PGP1-(PIASx)_3_ or AF568-PGP1-(DAXXc)_3_ to a final enzyme concentration of 50 nM. In matching solution only enzyme assays, the same concentration of AF568-PGP1-(SIM)_3_ construct was added to wells containing only PBS without condensates. PGP1 activity was initiated by adding pGlu-HCA to a final concentration of 4 µM. Formation of the fluorescent HCA product was monitored in a microplate reader (BioTek Synergy 2, Winooski, VT) with λ_ex_/λ_em_ = 670/700 nm. Substrate-only and condensate-only controls assays were included to confirm that changes in signal were due to enzyme-catalyzed hydrolysis of pGlu-HCA. For each biological replicate, fluorescence time courses from three technical replicates were collected. Traces were background-subtracted using the corresponding substrate-only control and normalized to the background fluorescence at t=0 s. For each replicate, the apparent initial rate was obtained from a linear fit to the initial portion of the fluorescence trace. Initial rates for each biological replicate were averaged across three technical replicates to identify differences in PGP1 activity in mixtures of dense and dilute phases from its activity in PBS alone.

### Enzyme kinetics by confocal microscopy

AF568-PGP1-(PIASx)_3_ was added to solutions of preformed mEGFP-(SUMO)_5_:mEGFP-(PIASx)_3_ condensates and AF568-PGP1-(DAXXc)_3_ was added to solutions of preformed mEGFP-(SUMO)_5_:mEGFP-(DAXXc)_3_ condensates to a final concentration of 50 nM. Enzyme recruitment to condensates was followed by microscopy for 1 h then pGlu-HCA was added to a final concentration of 2 µM. Preformed condensates were also imaged with pGlu-HCA without enzyme to ensure the absence of uncatalyzed probe hydrolysis. Enzyme activity was monitored over time for ∼30 min with λ_ex_ 670 nm and λ_em_ 710–799 nm using a Leica SP8X scanning confocal microscope. A total of 60 condensates for each enzyme were analyzed in ImageJ/FIJI to quantify mean fluorescence over time.

### Encapsulation efficiency of pGlu-HCA in condensates

pGlu-HCA fluorescence was measured on a Leica SP8X scanning confocal microscope with λ_ex_ 600 nm and λ_em_ 610-699 nm. Condensates containing either mEGFP-(SUMO)_5_:mEGFP-(PIASx)_3_ or mEGFP-(SUMO)_5_:mEGFP-(DAXXc)_3_ were prepared and 2 µM pGlu-HCA was added to each mixture. Thirty seconds after addition, fluorescence intensity was measured inside 10 discrete condensates and in 10 discrete ROI in the surrounding dilute phase. Encapsulation efficiency (EE) was calculated using the equation: 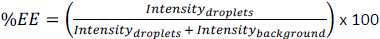

---

**Extended Figure 1.**
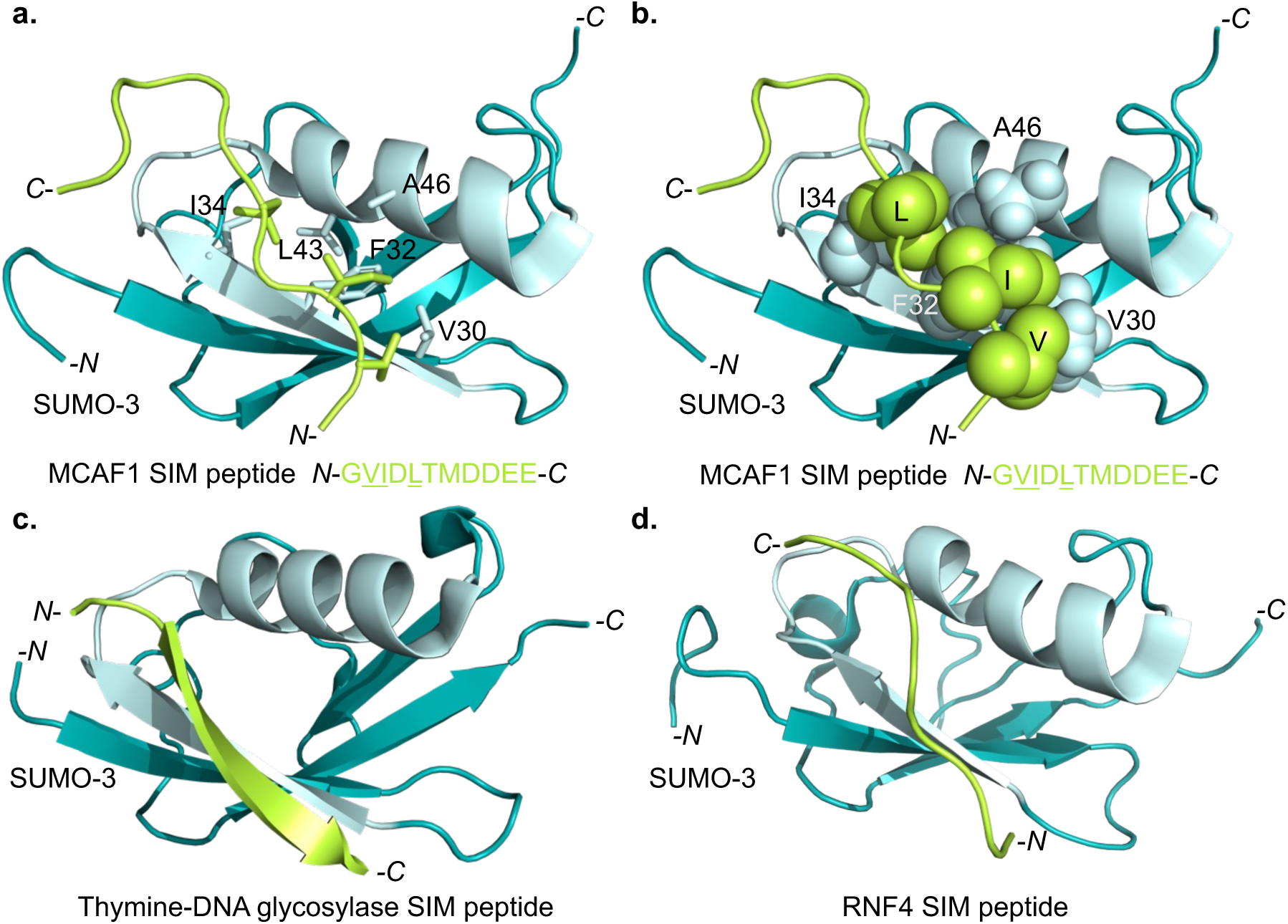
The conserved SUMO-interacting motif (SIM) binding groove in SUMO-3. **a**, NMR solution structure of SUMO-3 (teal) bound to a SIM peptide (green) from the MBD1-containing chromatin-associated factor 1 protein (MCAF1), PDB code 2RPQ.^16^ The SIM binding groove is shown in pale cyan and hydrophobic residues lining the groove are numbered based on the SUMO-3 sequence. **b,** Van der Waals interactions between hydrophobic side-chains in the SIM peptide (V, I and L) and the SUMO-binding groove are shown as spheres. **c,** X-ray crystal structure of SUMO-3 (teal) bound to a SIM peptide (green) from Thymine-DNA glycosylase, PDB code 2D07.^63^ The SIM binding groove is shown in pale cyan. **d,** NMR solution structure of SUMO-2 (teal) bound to the SIM-2 peptide (green) from RING-finger protein 4 (RNF4), PDB code 2MP2.^64^ The SIM binding groove is shown in pale cyan and SUMO-2/3 are identical in this region. Images were rendered with PyMol version 3.1.6.1.

**Extended Table 1.**
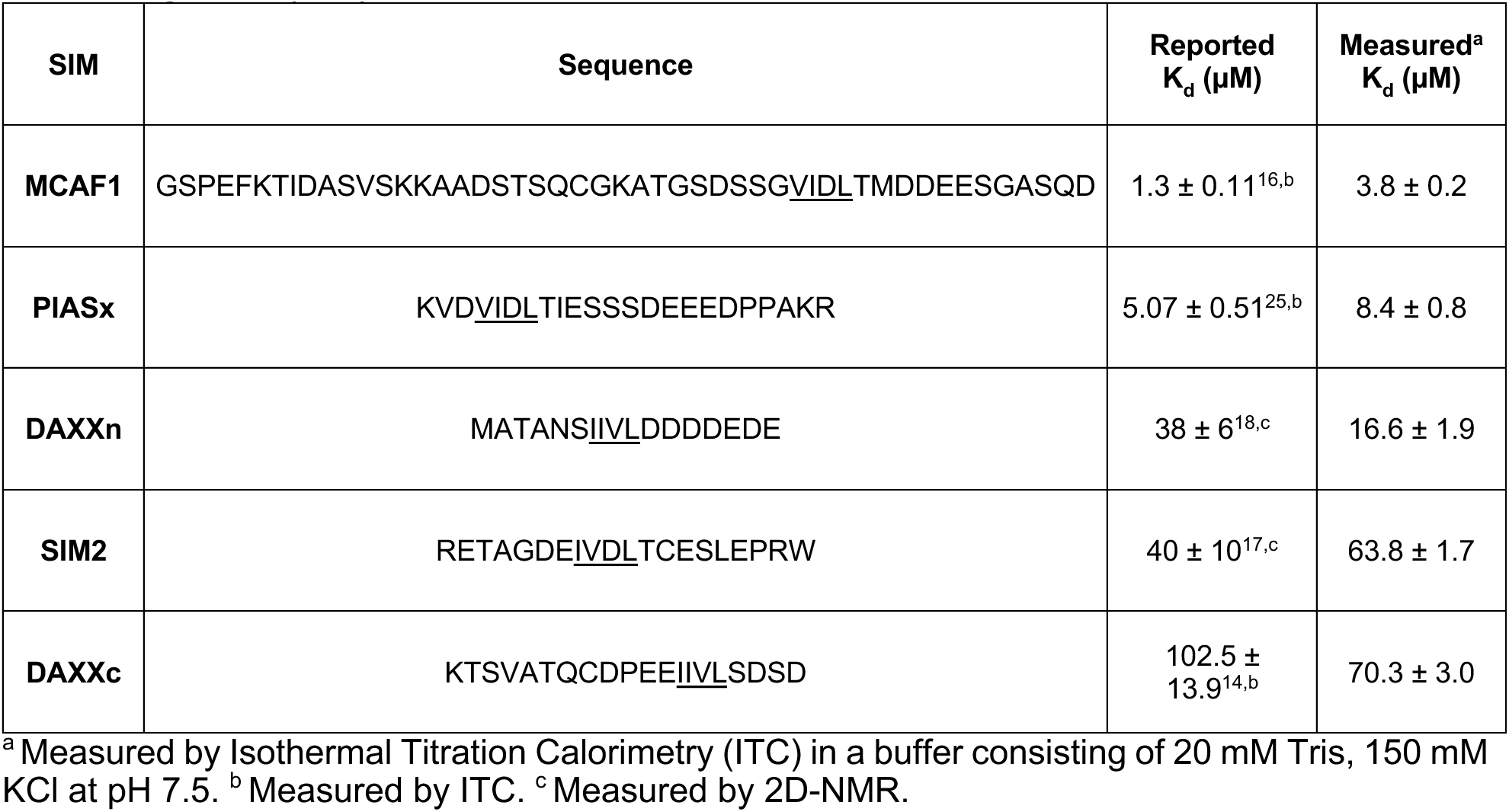
Experimentally measured K_d_ for monomeric SUMO-3 and SUMO-interacting motif (SIM) peptides.

**Extended Figure 2.**
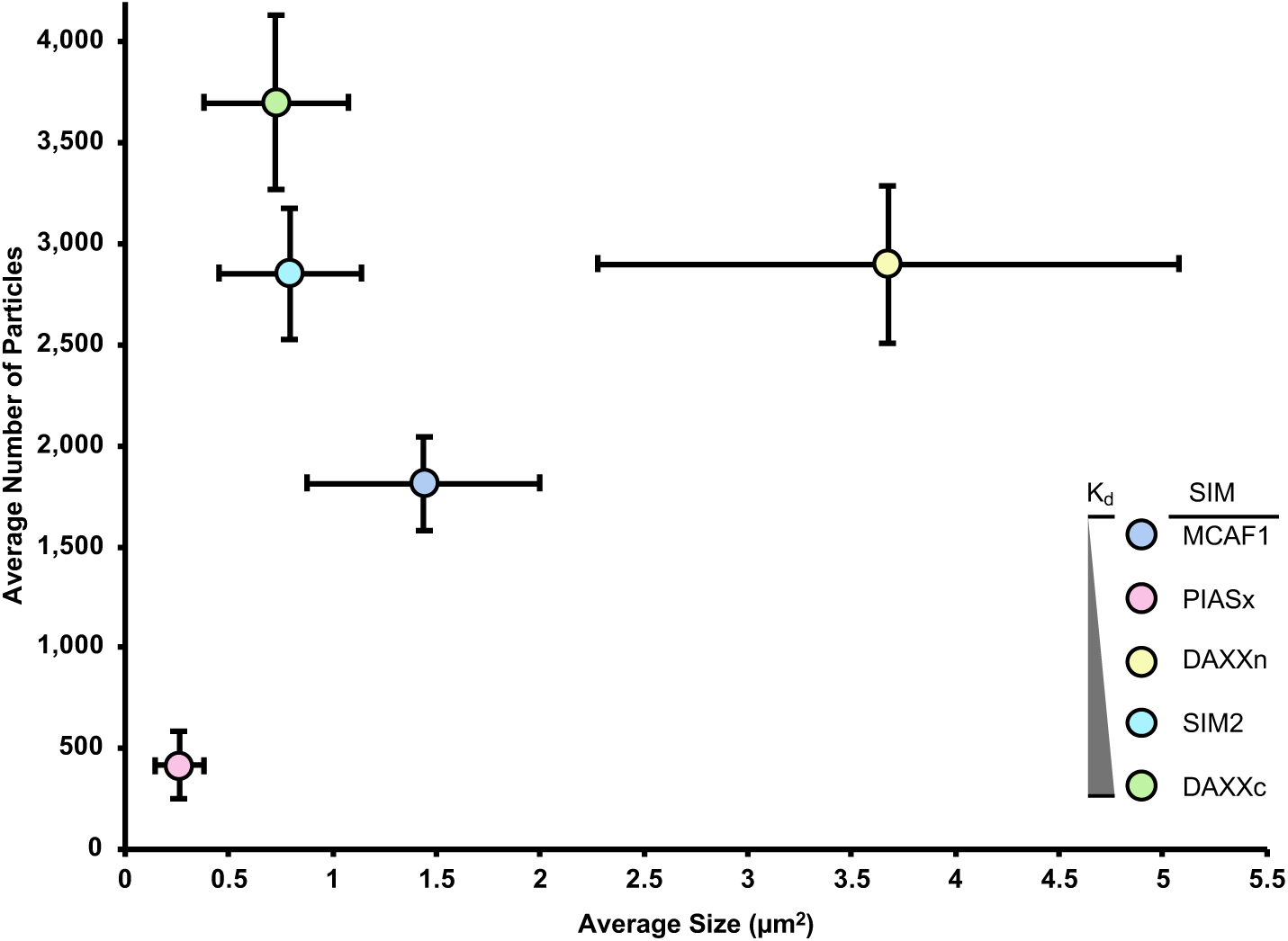
The size and number of condensates formed do not correlate with SUMO-SIM binding affinity. Condensates were assembled from mEGFP-(SUMO3)_5_ and the indicated mEGFP-(SIM)_3_ scaffolds. The number of condensates and their visible surface area was quantified from confocal fluorescence microscopy images. The average condensate surface area (µm^2^) and average number of condensates observed in the same sized field-of-view are shown for each SUMO-SIM pair. Neither the number of condensates nor their sizes were found to linearly correlate with SUMO-SIM binding affinity (K_d_). n= 3 independent replicates, filled circles indicate average number/area and error bars show s.e.m.

**Extended Table 2.** Physicochemical properties of SIM sequences.^65^.

| **SIM** | **Number of residues** | **Molecular Weight (Da)** | **Isoelectric Point (pI)** | **Grand Average of Hydropathicity (GRAVY)<sup>66</sup>** | **Positively Charged Residues** | **Negatively Charged Residues** | **Net Charge at pH 7.5<sup>a</sup>** |
| --- | --- | --- | --- | --- | --- | --- | --- |
| **MCAF1** | 49 | 4,912.16 | 4.03 | -0.706 | 4 (8%) | 10 (20%) | -6.3 |
| **PIASx** | 23 | 2,572.76 | 4.07 | -1.026 | 3 (13%) | 8 (35%) | -5.2 |
| **DAXXn** | 17 | 1,865.94 | 3.18 | -0.412 | 0% | 7 (41%) | -7.2 |
| **SIM2** | 20 | 2,319.53 | 4.08 | -0.725 | 2 (10%) | 6 (30%) | -4.3 |
| **DAXXc** | 20 | 2,150.34 | 3.77 | -0.240 | 1 (5%) | 5 (25%) | -4.3 |

<sup>a</sup> Net charge at pH 7.5 was estimated using the Henderson–Hasselbalch equation with standard model pKa values for ionizable groups and free termini.<sup>67,68</sup> Charges were treated as fractional based on protonation equilibria, yielding average solution-state values that do not account for structure or sequence-dependent pKa shifts.

**Extended Figure 3.**
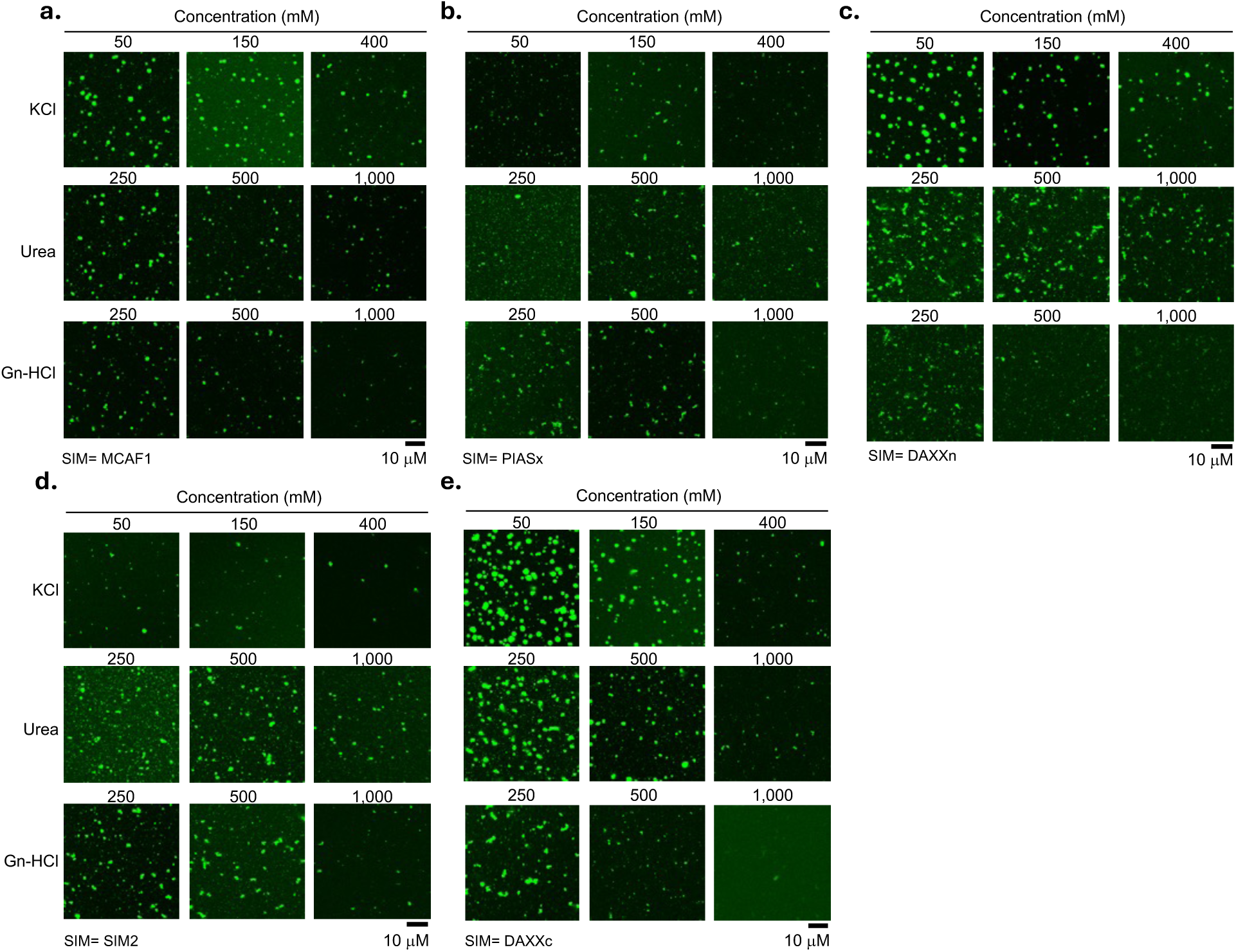
Condensate stability to salt and chaotropes correlates with scaffold affinity. Condensates assembled with mEGFP-(SUMO)_5_ and the indicated mEGFP-(SIM)_3_ scaffold at concentrations above c_sat_ for both SUMO and SIM were incubated with increasing concentrations of salt (KCl) or chemical denaturants urea and guanidinium chloride (Gn-HCl). Confocal fluorescence microscopy images for each SUMO-SIM pair are displayed as a 3×3 grid with concentrations increasing from left to right. Top row, incubation with 50 mM, 150 mM, and 400 mM KCl. Middle row, incubation with 250 mM, 500 mM, and 1,000 mM Urea. Bottom row, incubation with 250 mM, 500 mM, and 1,000 mM Gn-HCl. **a,** MCAF1 condensates. **b,** PIASx condensates. **c,** DAXXn condensates. **d,** SIM2 condensates. **e,** DAXXc condensates. Increasing KCl, urea or guanidinium chloride reduced the number of condensates, with weaker-binding SUMO-SIM pairs exhibiting loss of condensates at lower perturbant concentrations. n= 6 independent replicates and scale bar is 10 µm.

**Extended Figure 4.**
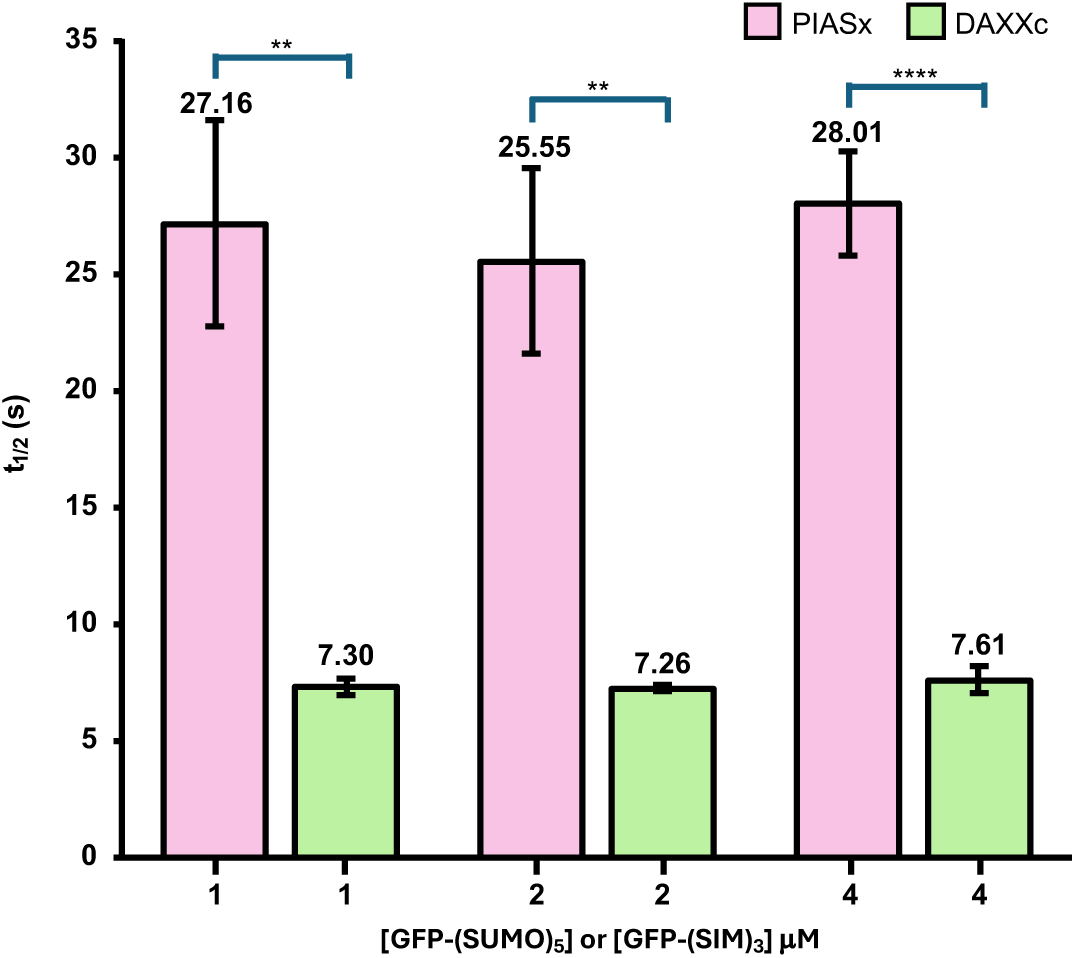
Effect of changing SIM concentrations on FRAP half-times (t_1/2_) of condensates. Fluorescence recovery half-times (t_1/2_) were calculated from fits to the normalized recovery traces for condensates formed by mEGFP-(SUMO)_5_ and mEGFP-(PIASx)_3_ (pink bars) and mEGFP-(SUMO)_5_ and mEGFP-(DAXXc)_3_ (green bars) assembled at 1, 2 or 4 µM of each scaffold. The difference in intra-condensate dynamics remained unchanged across the concentration range tested. n= 6-9 independent replicates and error bars show s.e.m. Statistical significance was calculated using Welch’s two-tailed t-test. **= p< 0.01, ****= p<0.0001.

**Extended Figure 5.**
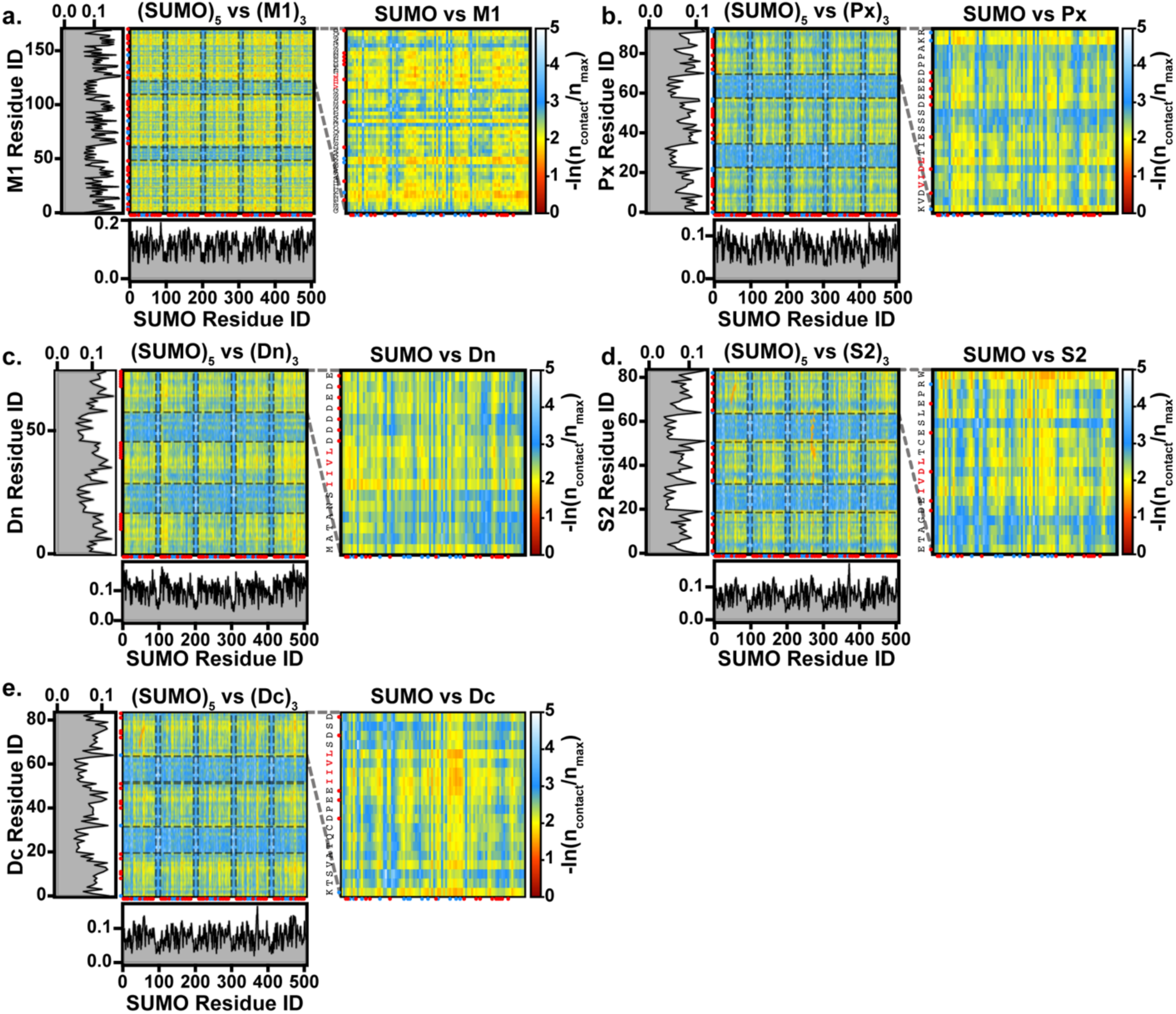
Intermolecular contact maps between (SUMO)₅ and (SIM)₃ from 5 µs CG coexistence simulations at ε = 0.33. Left panels show residue-level contact propensities between all residues of the (SUMO)₅ multimer and the corresponding (SIM)₃ construct, while right panels show the same analysis zoomed in on a single SUMO–SIM pair. Color scale represents preferential contacts, where warm colors (red) indicate high-probability contacts and cool colors (green/blue) indicate low-probability or transient contacts. Marginal histograms show the per-residue contact frequency summed over the partner chain.

**Extended Figure 6.**
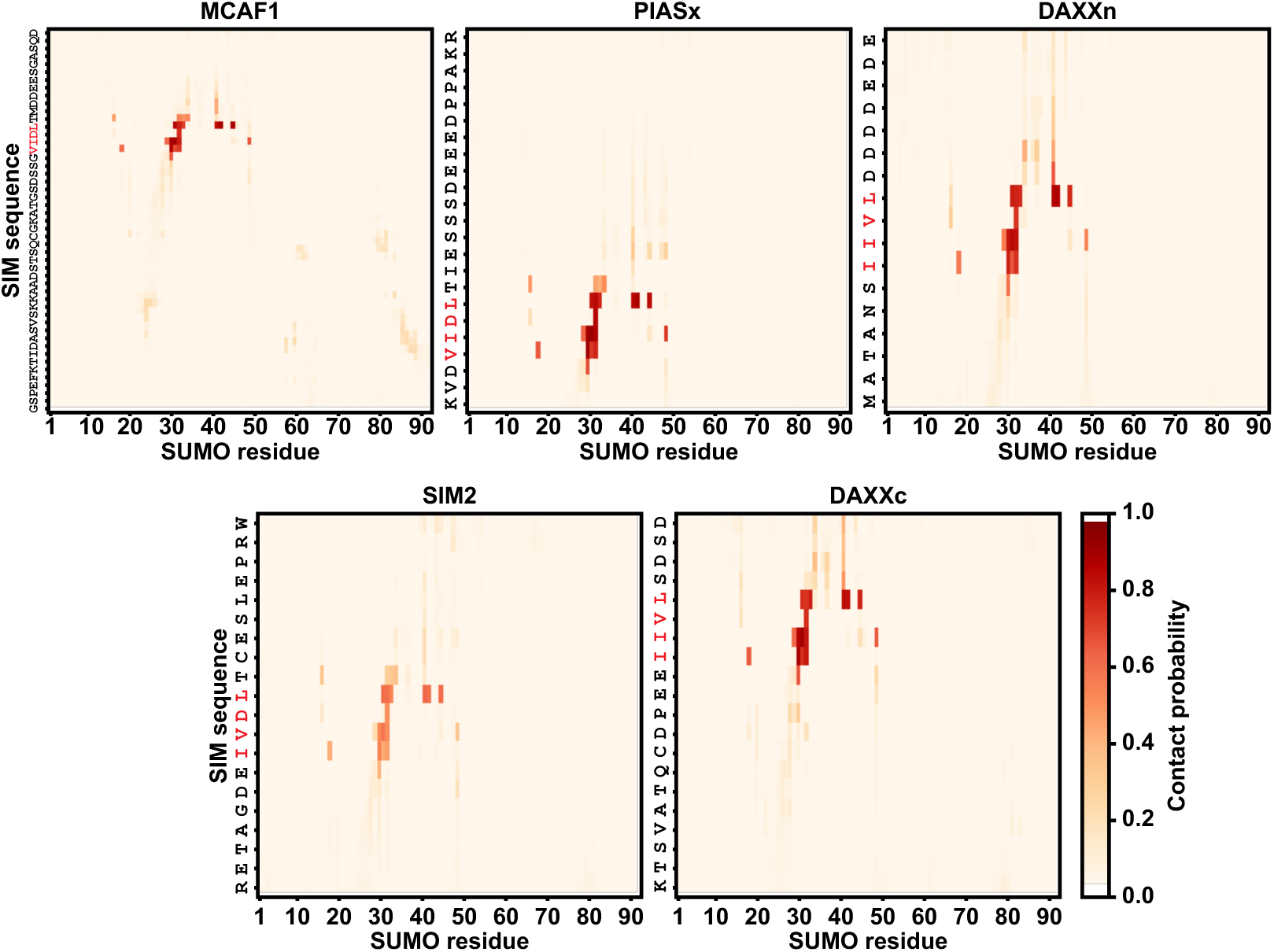
Intermolecular residue-level contact maps between SUMO and SIM variants from all-atom (AA) simulations. The color scale represents contact probability, with warm colors (red) indicating high-probability contacts and cool colors indicating low-probability or transient contacts. Sequence is annotated on the y-axis and SIM-binding motif residues are highlighted in red.

**Extended Figure 7.**
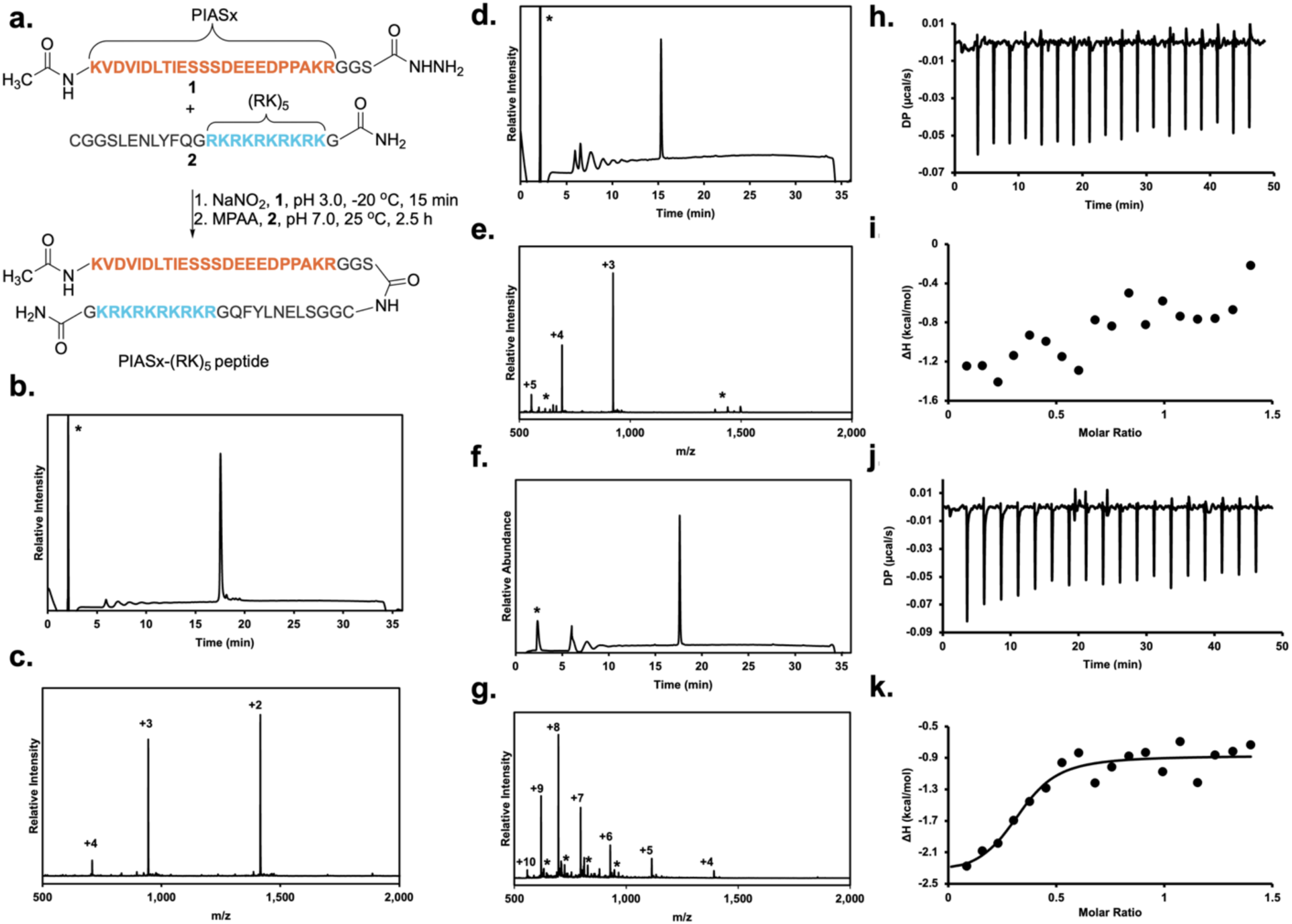
Synthesis of PIASx-(RK)_5_ by native chemical ligation and its binding to SUMO at pH 6.5. **a**, Synthetic scheme to access the 50-mer PIASx-(RK)_5_ peptide. MPAA= 4-mercaptophenylacetic acid. **b,** C18 analytical HPLC chromatogram of the pure hydrazide peptide **1** injected on a gradient of 0-73% B over 30 min. An asterisk indicates the unrelated injection peak. **c,** ESI-MS of pure hydrazide peptide **1**. Observed [M]^+^ = 2,829.6 ± 0.4 Da, calculated [M]^+^ = 2,829.9 Da. **d,** C18 analytical HPLC chromatogram of the pure cysteine peptide **2** injected on a gradient of 0-73% B over 30 min. The asterisk indicates the injection peak. **e,** ESI-MS of pure cysteine peptide **2**. Observed [M]^+^ = 2,765.0 ± 0.1 Da, calculated [M]^+^ = 2,765.3 Da. Asterisks indicate fragment ions. **f,** C18 analytical HPLC chromatogram of the PIASx-(RK)_5_ ligation product injected on a gradient of 0-73% B over 30 min. An asterisk indicates the injection peak. **g,** ESI-MS of pure PIASx-(RK)_5_. Observed [M]^+^ = 5,564.3 ± 1.0 Da, calculated [M]^+^ = 5,563.2 Da. Asterisks indicate TFA adducts during ESI-MS. **h,** Representative thermogram of SUMO titration into a solution of the PIASx peptide at pH 6.5. **i,** Integrated heat plot for the thermogram in **h** showing the lack of binding at pH 6.5. **j,** Representative thermogram of SUMO titration into a solution of the PIASx-(RK)_5_ peptide at pH 6.5. **k,** Integrated heat plot for the thermogram in **j** showing robust binding at pH 6.5.

**Extended Figure 8.**
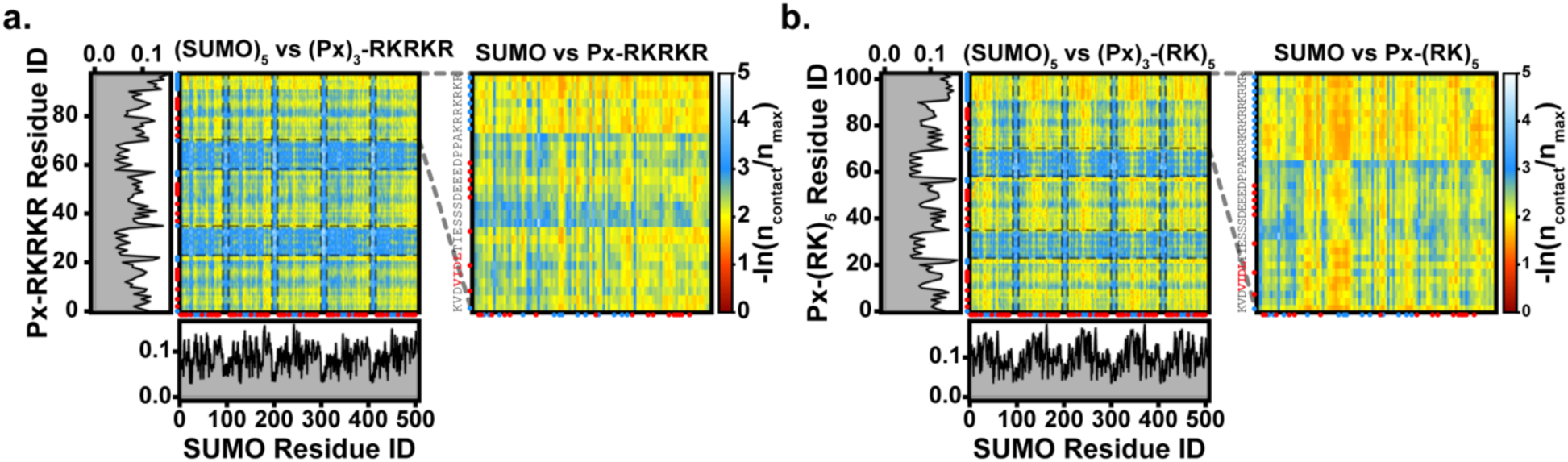
CG intermolecular contact maps within the dense phase for (SUMO)₅ with polybasic SIM constructs at ε = 0.33. **a**, Intermolecular contact maps between (SUMO)₅ and (PIASx)₃-RKRKR from a 5 µs CG slab simulation. Left: full multivalent contact map between the (SUMO)₅ and (PIASx)₃-RKRKR system. Red and blue dots on the axes indicate the positively and negatively charged residues along the sequence, respectively. The dashed gray line on the y-axis separates the PIASx SIM core sequence from the C-terminal RKRKR extension. Marginal line plots show the normalized per-residue contact frequency along the SUMO (bottom) and PIASx-RKRKR (left) chain axes. Right: zoom-in contact map for a single SUMO–PIASx-RKRKR pair, with the full PIASx-RKRKR sequence annotated on the y-axis and SIM-binding motif residues highlighted in red. Color scale indicates contact propensity where warmer colors denote higher contact frequency and cooler colors denote lower contact frequency. **b,** As in (a) for (SUMO)₅ with (PIASx)₃-(RK)₅ from a 5 µs CG slab simulation.

**Extended Figure 9.**
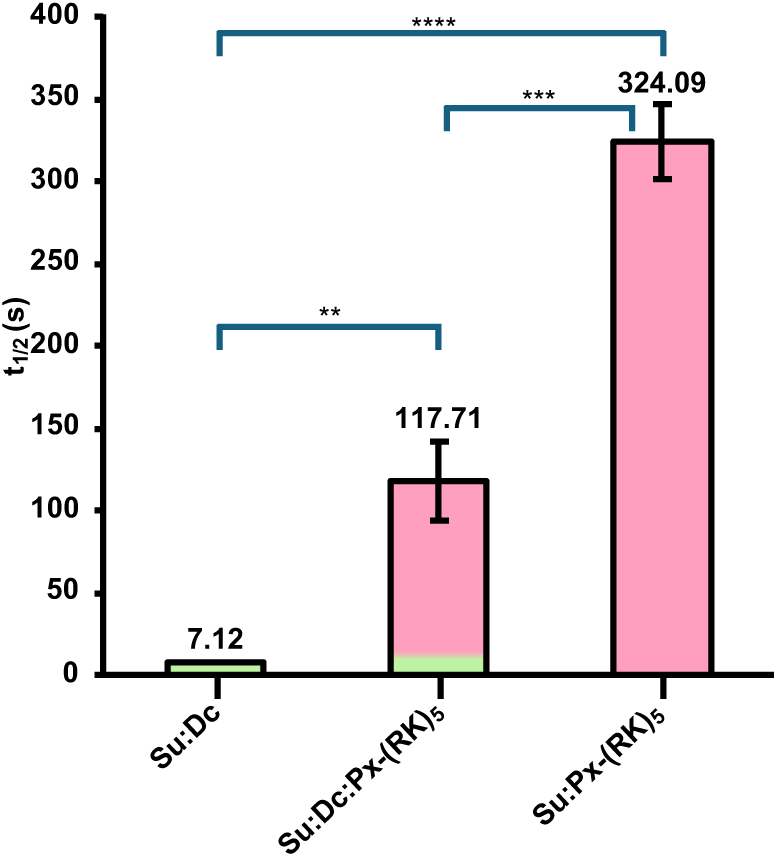
Intra-condensate dynamics of complex condensates are dictated by the highest affinity scaffold. Average FRAP t_1/2_ for the dense phase formed by binary condensates containing mEGFP-(SUMO)_5_ and either mEGFP-(DAXXc)_3_ or mEGFP-(PIASx)_3_-(RK)_5_, and a ternary mixture containing all three proteins, calculated from fitting the normalized recovery curves. n= 6-11 independent replicates and error bars show s.e.m. Su= SUMO, Dc= DAXXc, Px-(RK)_5_= PIASx-(RK)_5_. Statistical significance was calculated using Welch’s two-tailed t-test. **= p< 0.01, ***= p<0.001, ****= p<0.0001.

**Extended Figure 10.**
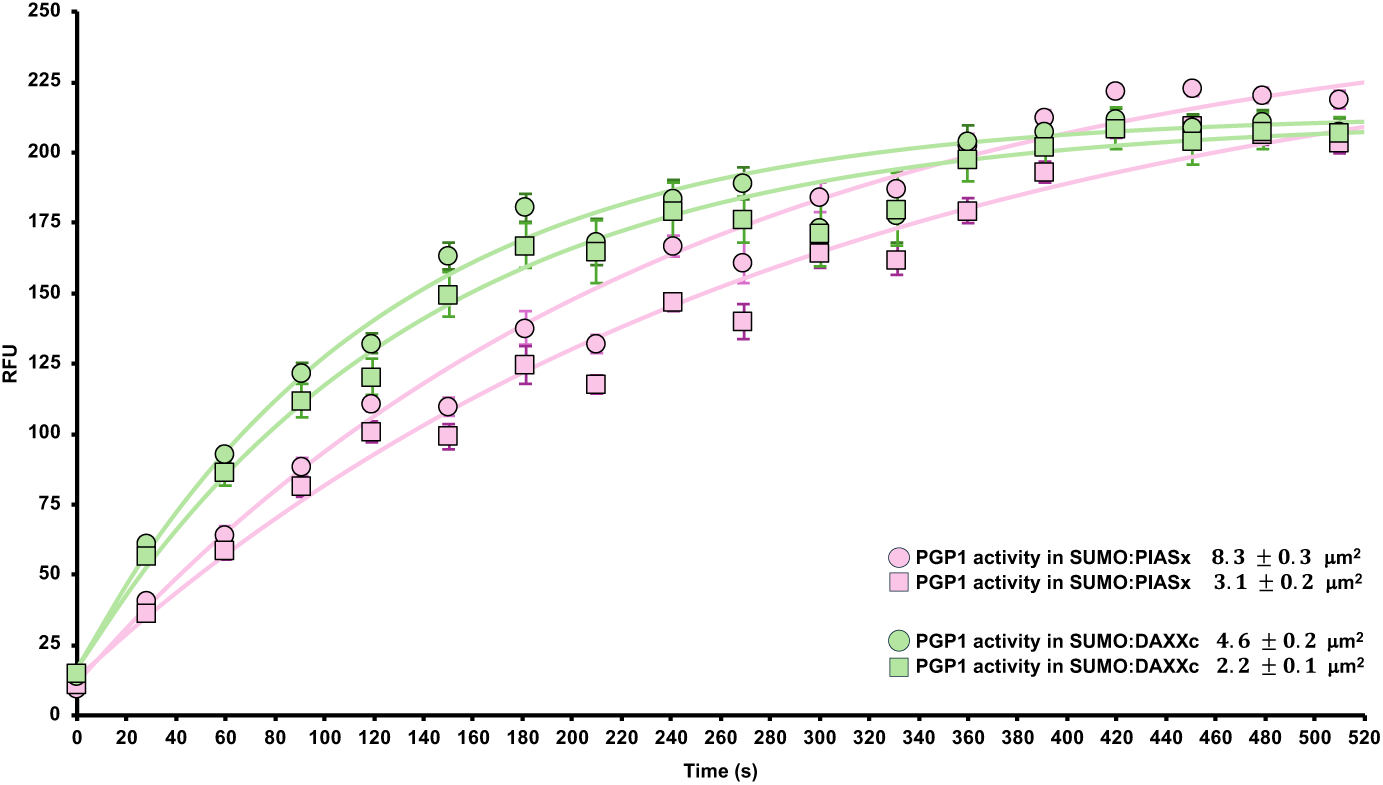
PGP1 activity in different sized condensates. HCA fluorescence in condensates imaged at λ_em_ 700 nm was quantified from time-lapse images of SUMO:DAXXc:AF568-PGP1-(DAXXc)_3_ or SUMO:PIASx:AF568-PGP1-(PIASx)_3_ condensates incubated with 50 nM pGlu-HCA probe and plotted as a function of time. HCA fluorescence from SUMO:PIASx:AF568-PGP1-(PIASx)_3_ was measured in condensates of 8.7 ± 0.3 µm^2^ or 3.1 ± 0.2 µm^2^ area. HCA fluorescence from SUMO:DAXXc:AF568-PGP1-(DAXXc)_3_ was measured in condensates of 4.6 ± 0.2 µm^2^ or 2.2 ± 0.1 µm^2^ area. Markers denote the average fluorescence and error bars indicate s.e.m. from n= 30 similarly sized droplets.

**Supplementary Figure S1.**
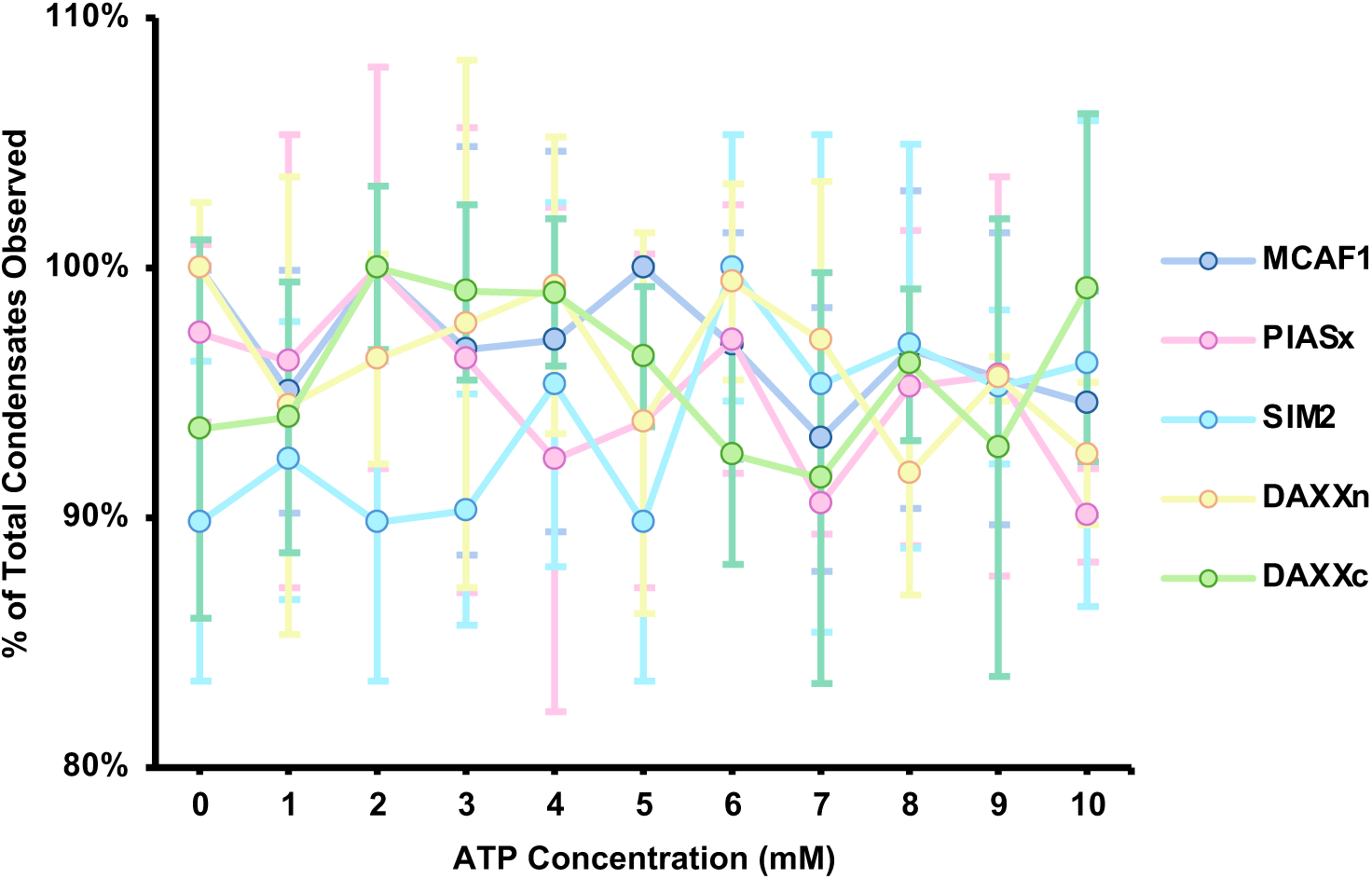
Stability of SUMO-SIM condensates to the biological hydrotrope ATP. Condensates assembled from mEGFP-(SUMO)_5_ and the indicated mEGFP-(SIM)_3_ scaffolds were subjected to increasing final concentrations of ATP. The final number of condensates in the presence of ATP was quantified by confocal fluorescence microscopy within the same sized field of view and values were normalized to the number of condensates observed in the absence of ATP. n =3 independent replicates, filled circles indicate the average number of condensates and error bars show s.d.

**Supplementary Figure S2.**
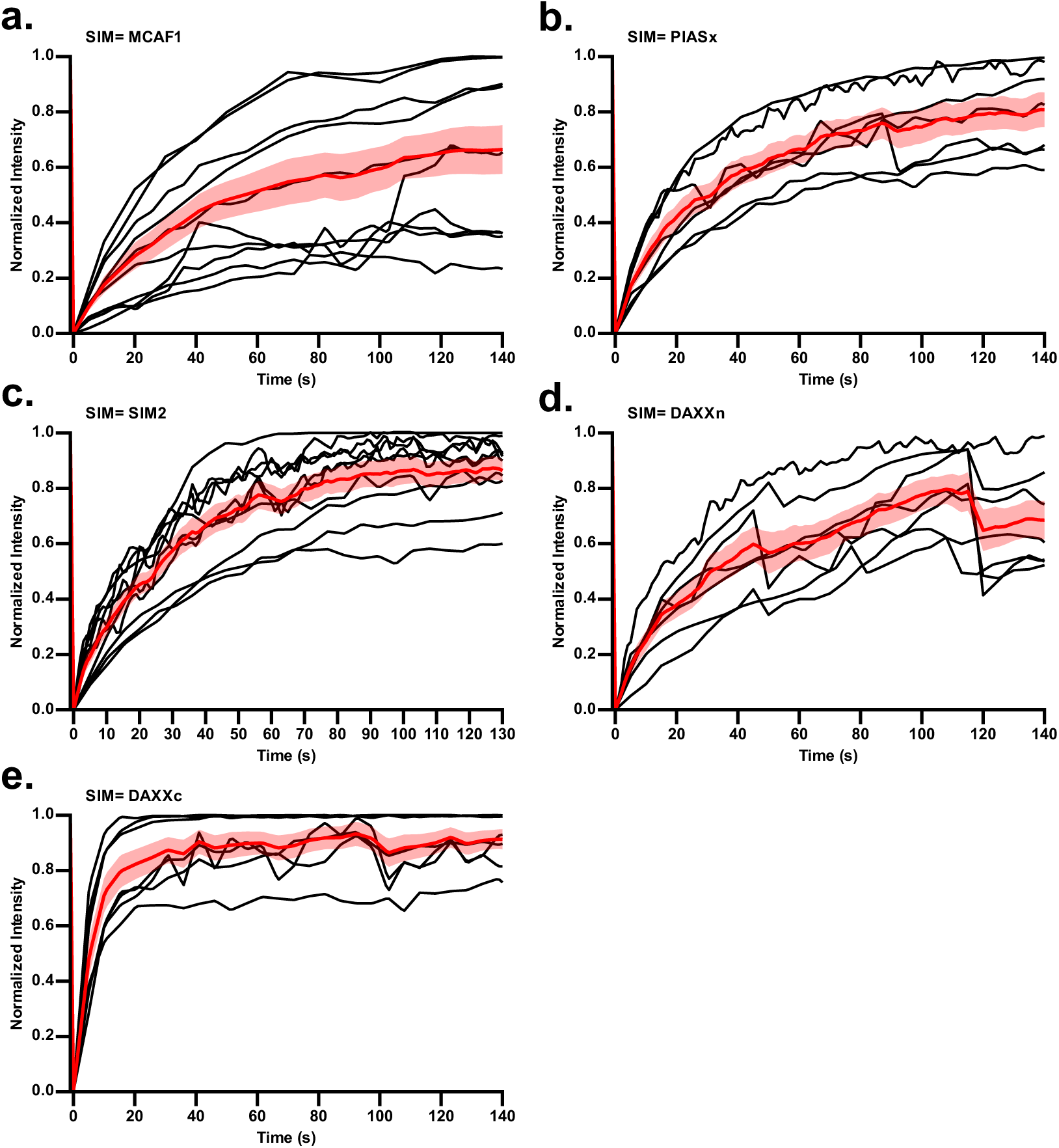
Fluorescence recovery curves after laser-mediated photobleaching of SUMO-SIM scaffolded condensates. A circular region corresponding to ∼50% of the droplet area was photobleached and normalized fluorescence intensity within the bleached region was measured by confocal fluorescence microscopy. Normalized intensity of bleached spots over time for condensates formed by mixing mEGFP-(SUMO)_5_ with **a,** mEGFP-(MCAF1)_3_. **b,** mEGFP-(PIASx)_3_. **c,** mEGFP-(SIM2)_3_. **d,** mEGFP-(DAXXn)_3_. **e,** mEGFP-(DAXXc)_3_. Black traces show individual recovery curves, the red trace shows the mean recovery curve, and the red shaded region indicates s.e.m. across condensates. n≥6 independent replicates and error bars show s.e.m.

**Supplementary Figure S3.**
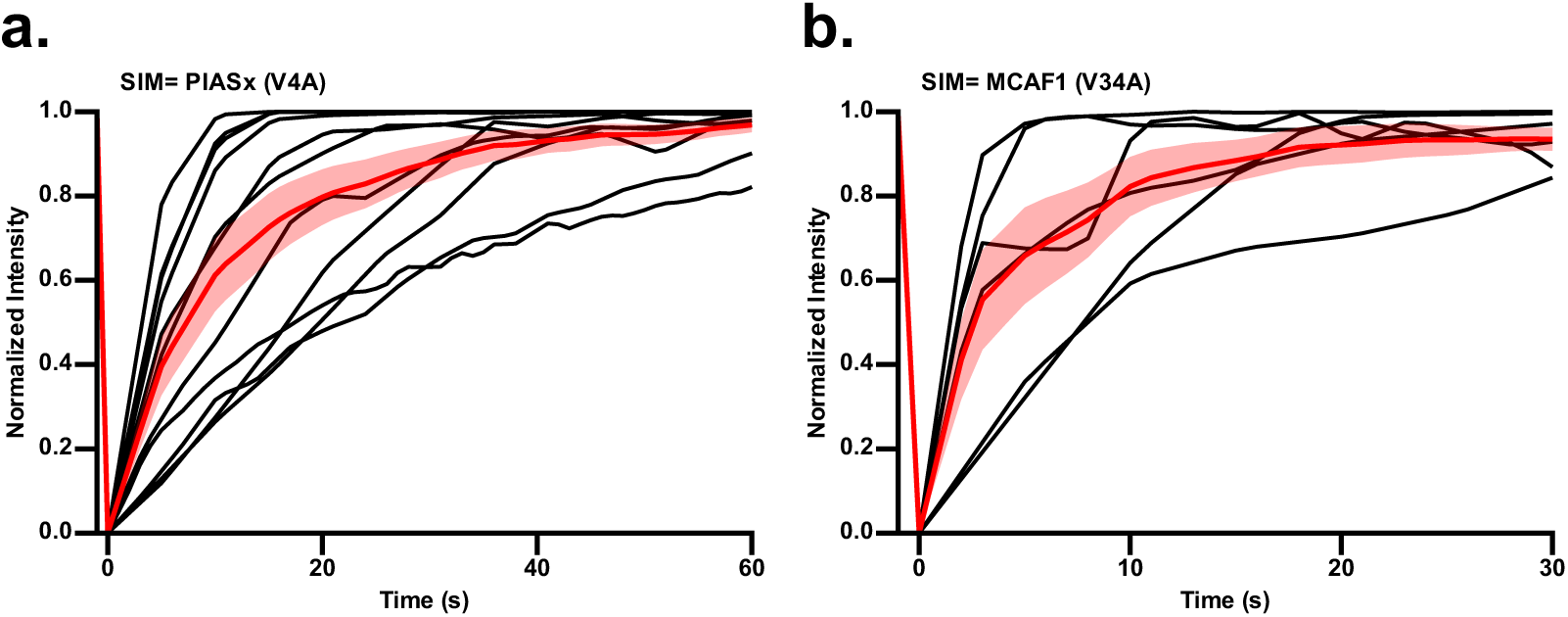
Fluorescence recovery curves after laser-mediated photobleaching of condensates generated with hydrophobic core mutated SIMs. A circular region corresponding to ∼50% of the droplet area was photobleached and normalized fluorescence intensity within the bleached region was measured by confocal fluorescence microscopy. Normalized intensity of bleached spots over time for condensates formed by mixing mEGFP-(SUMO)_5_ with **a,** mEGFP-(PIASx(V4A))_3_. **b,** mEGFP-(MCAF1(V34A))_3_. Black traces show individual recovery curves, the red trace shows the mean recovery curve, and the shaded red region indicates s.e.m. across droplets. n≥4 independent replicates and error bars show s.e.m.

**Supplementary Figure S4.**
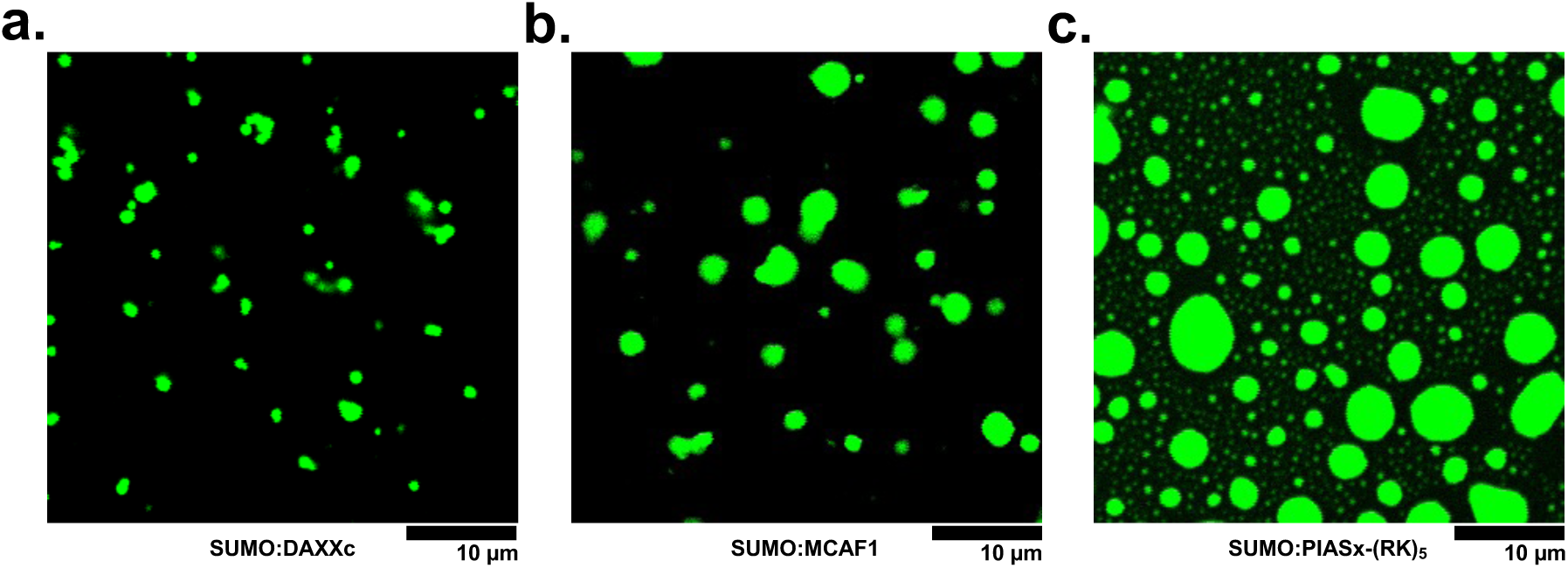
Control condensates formed by binary mixtures of mEGFP-(SUMO)_3_ and mEGFP-(SIM)_5_ scaffolds. Confocal fluorescence microscopy images of condensates formed by equimolar amounts of mEGFP-(SUMO)_5_ and **a,** mEGFP-(DAXXc)_3_ **b,** mEGFP-(MCAF1)_3_ and **c,** mEGFP-(PIASx)_3_-(RK)_5_. Images were acquired under identical imaging settings with the same field-of-view. Scale bar is 10 µm.

**Supplementary Figure S5.**
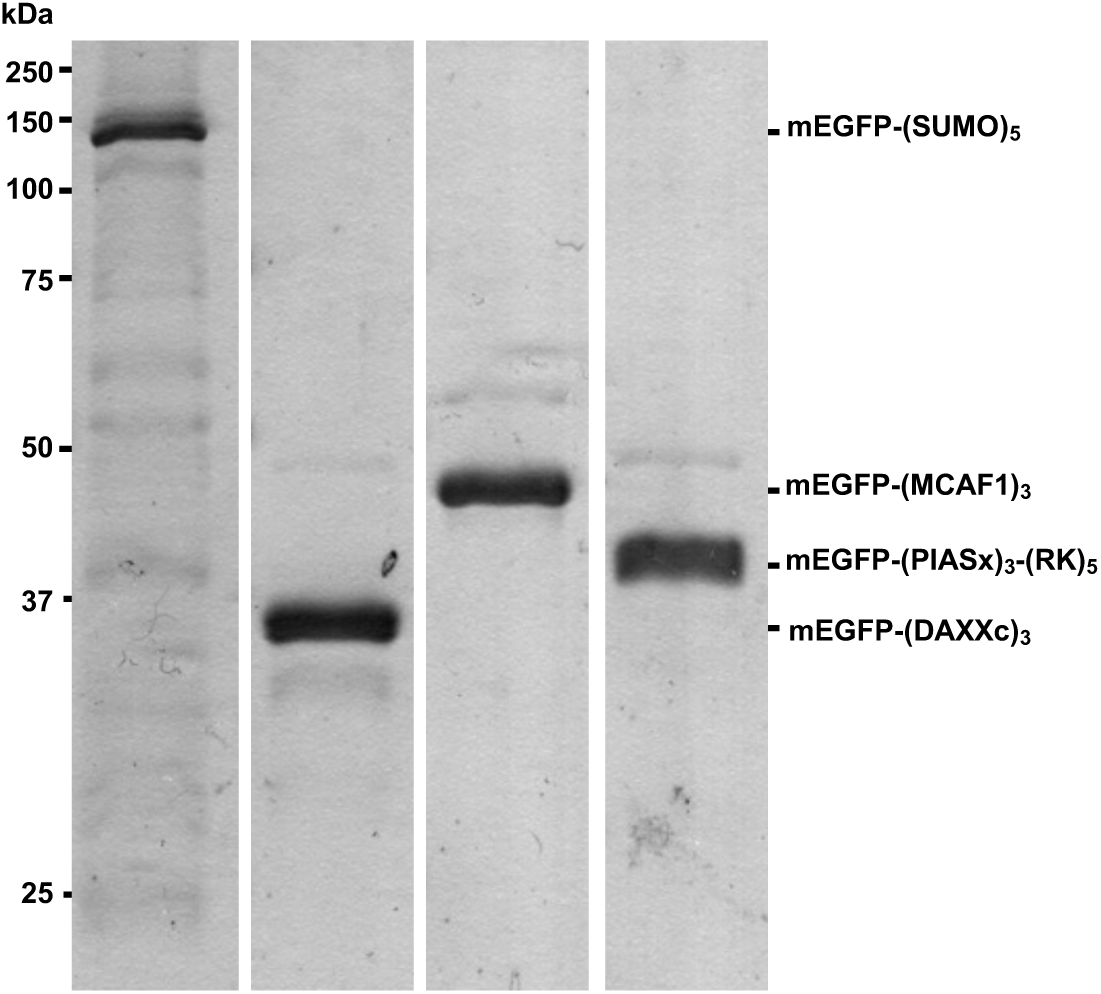
Characterization of protein scaffolds employed in quaternary mixtures for condensate formation. Coomassie-stained 10% SDS-PAGE gels of the four purified proteins, mEGFP-(SUMO)_5_, mEGFP-(DAXXc)_3_, mEGFP-(MCAF1)_3_ and mEGFP-(PIASx)_3_-(RK)_5_ employed in quaternary mixtures for condensate formation.

**Supplementary Figure S6.**
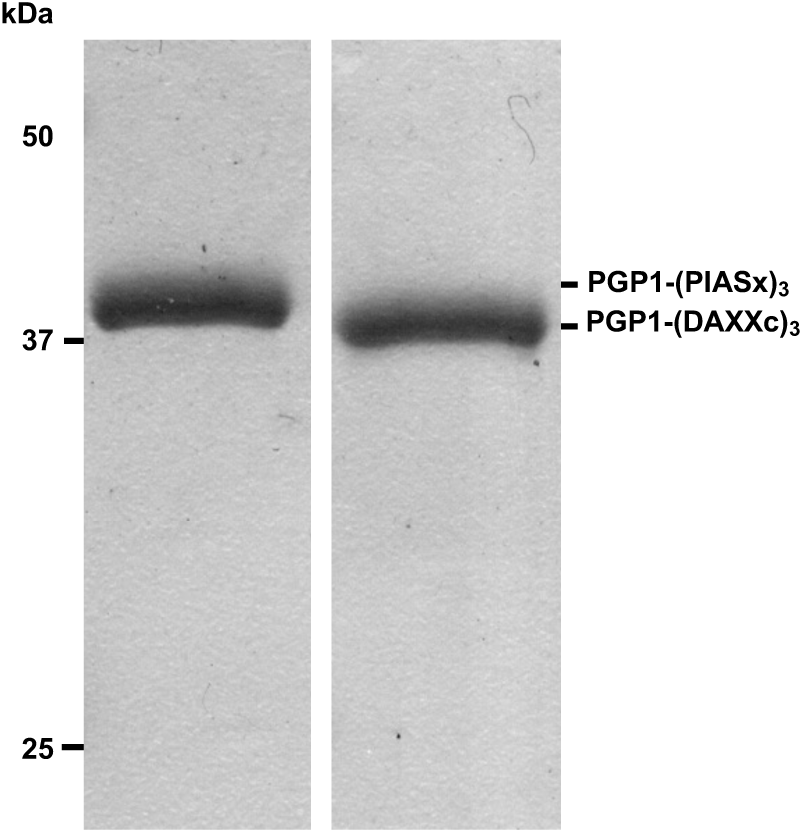
Characterization of PGP1-(SIM)_3_ fusion proteins. Coomassie-stained 10% SDS-PAGE of purified PGP1-(SIM)_3_ fusions. PGP1-(DAXXc)_3_ (left lane) and PGP1-(PIASx)_3_ (right lane) were purified to homogeneity prior to labeling with the Alexa Fluor568 dye.

**Supplementary Figure S7.**
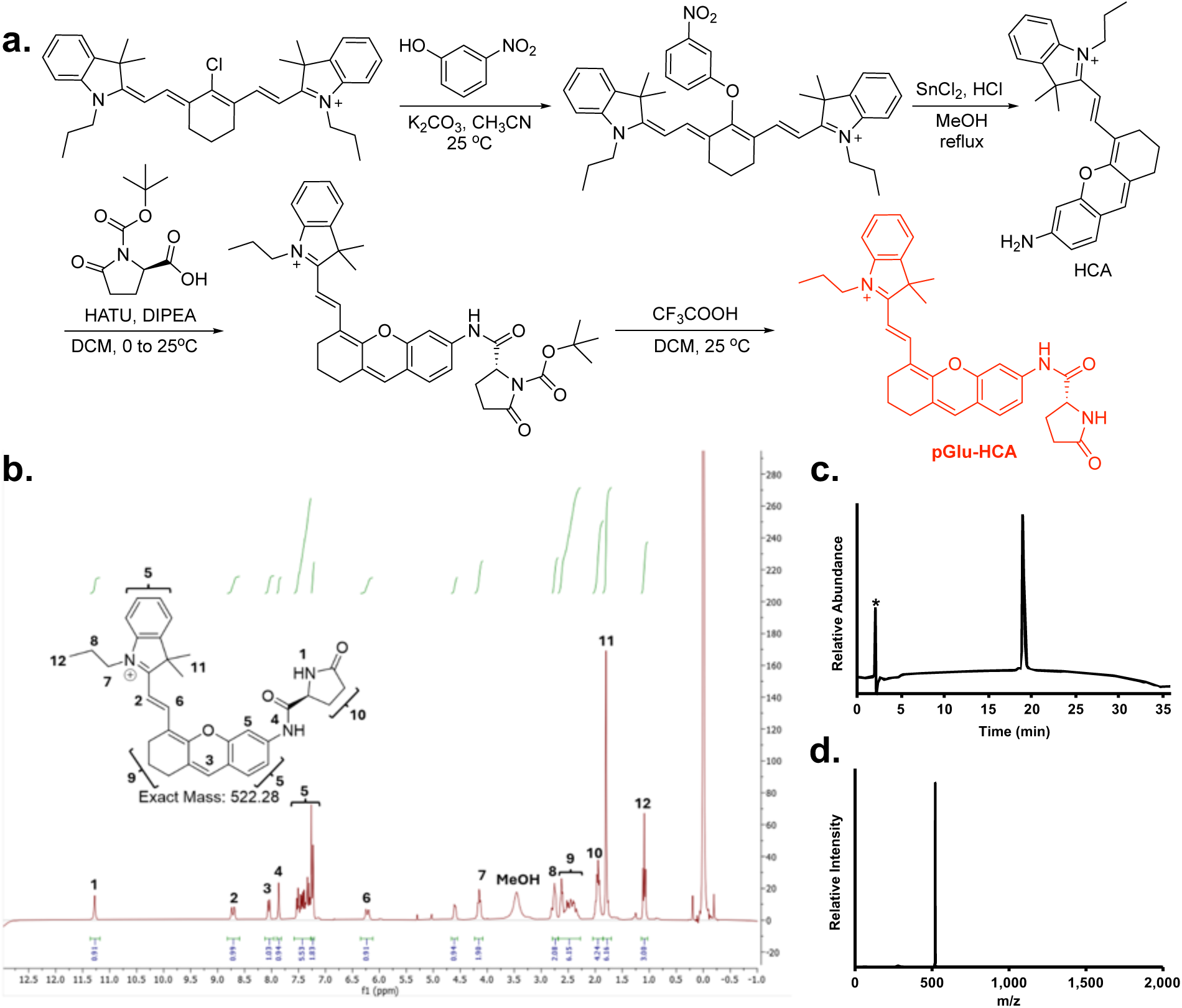
Synthesis and characterization of the near-IR fluorogenic probe pyroglutamyl-hemicyanine (pGlu-HCA). **a**, Synthetic scheme to access pGlu-HCA.^1^ HATU = O-(7-azabenzotriazol-1-yl)-*N,N,N′,N′*-tetramethyluronium hexafluorophosphate, DIPEA = *N,N*-Diisopropylethylamine. **b,** 300 MHz ^1^HNMR spectrum of pGlu-HCA in CDCl_3_. **c,** Analytical C18 HPLC chromatogram of pure pGlu-HCA acquired on a 20-100% B gradient over 30 min. An asterisk indicates the injection peak. **d,** ESI-MS of purified pGlu-HCA. Observed [M]^+^=522.36 Da, calculated [M]^+^=522.28 Da.

**Supplementary Figure S8.**
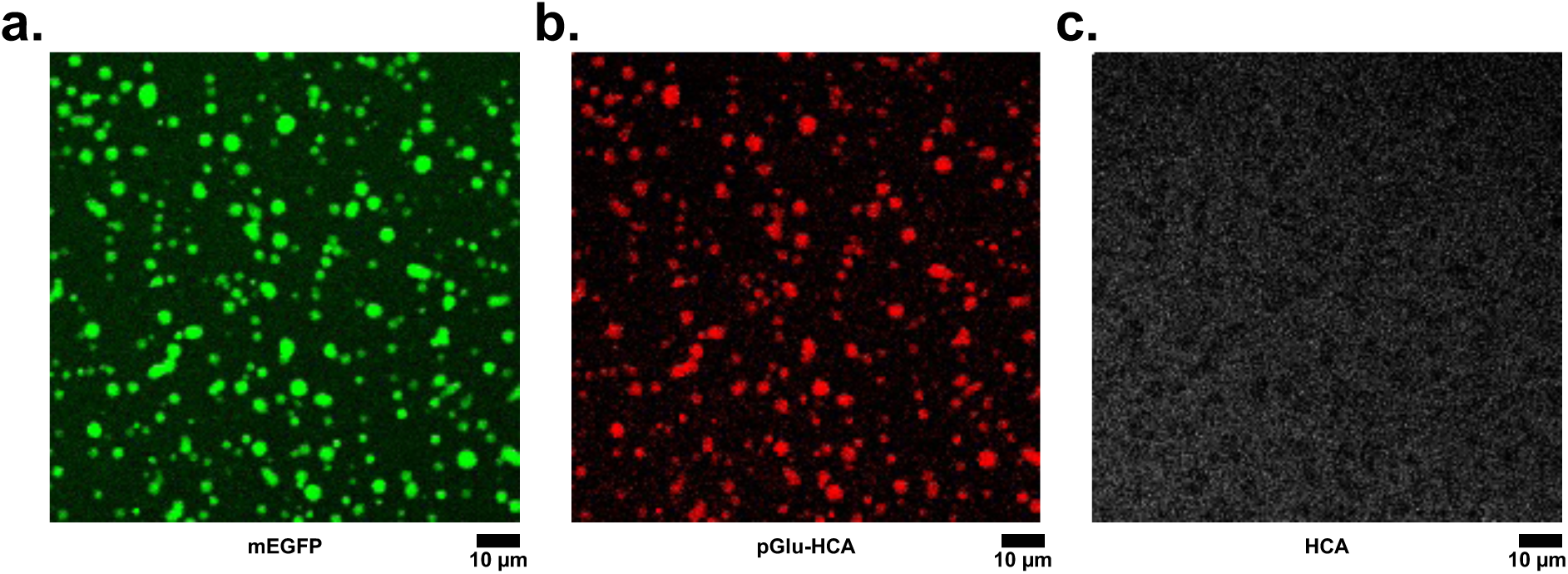
Stability of the pGlu-HCA probe under assay conditions. Background hydrolysis of pGlu-HCA in condensates was monitored by near-IR fluorescence (λ_ex/em_ 670/700 nm) over 20 min under conditions used for measuring PGP1 activity. **a,** mEGFP fluorescence from SUMO:PIASx condensates generated from 2 μm of each protein. **b,** Fluorescence from pGlu-HCA added to bulk solution showing retention of the probe within condensates. **c,** Near-IR fluorescence acquired under HCA-specific imaging conditions showing no detectable hydrolysis of pGlu-HCA in the absence of recruited PGP1. Images were acquired in the same field-of-view under identical settings. Scale bar is 10 μm.

**Supplementary Figure S9.**
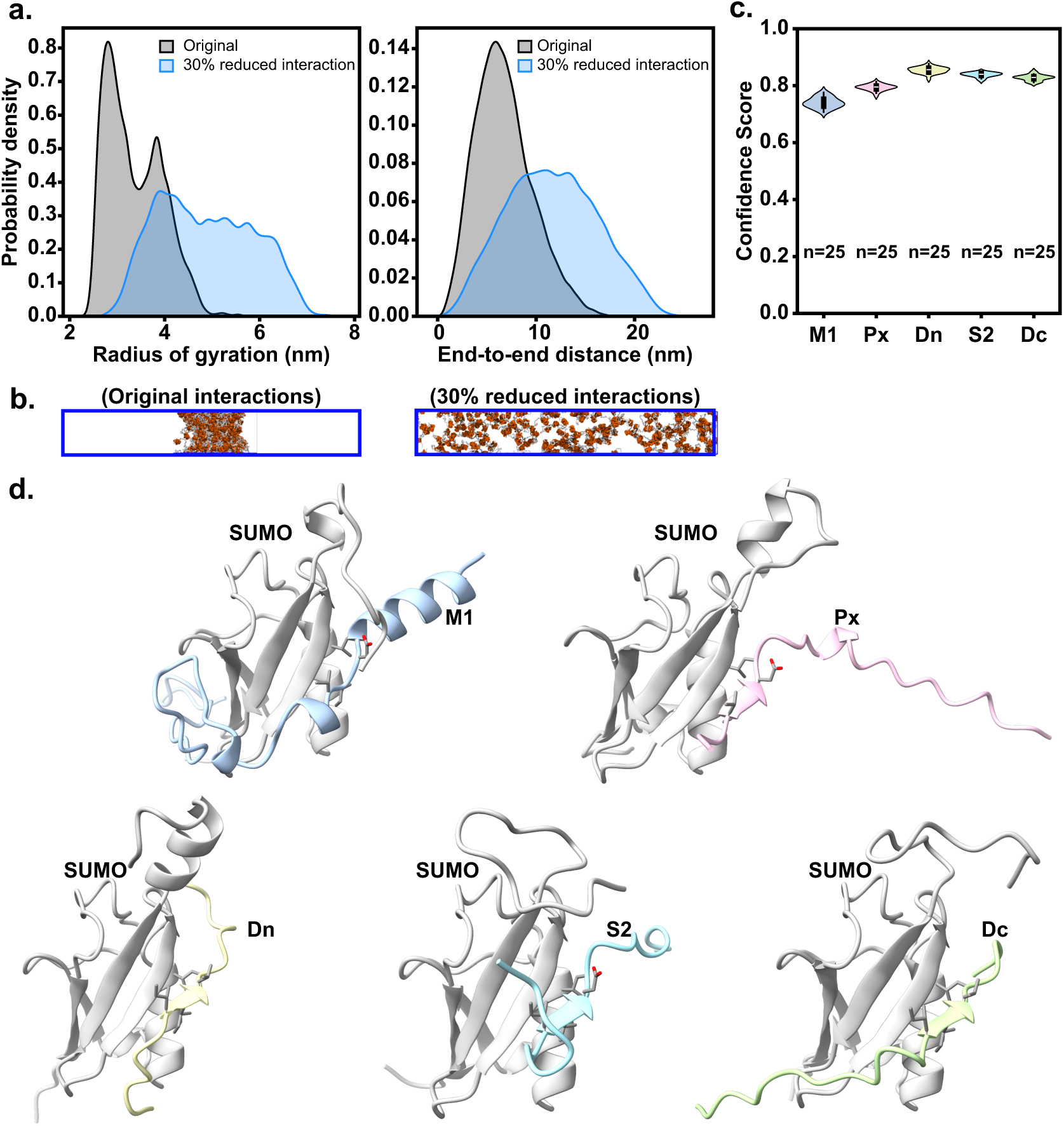
Validation of coarse-grained parametrization and all-atom initial structures for SUMO-SIM simulations. **a**, Probability density distributions of the radius of gyration (Rg, left) and end-to-end distance (Ree, right) from single-chain CG simulations of (SUMO)₅ using original Urry interaction parameters (gray) and with SUMO domain interactions reduced by 30% (blue). **b,** Representative snapshots from CG simulations showing artificial self-condensation with original parameters (left) and dispersed chains upon 30% interaction reduction (right). **c,** Violin plots of Boltz2 prediction confidence scores for 25 structural models of each SUMO-SIM complex (M1, Px, Dn, S2, and Dc). All constructs show consistently high confidence scores (>0.75), supporting the reliability of the predicted binding poses used as starting structures for all-atom simulations. **d,** Representative Boltz2-predicted structures of SUMO (gray) bound to each SIM peptide: M1 (blue), Px (pink), Dn (yellow-green), S2 (cyan), and Dc (light green). Sticks indicate the conserved hydrophobic core residues of each SIM. All SIM peptides dock into the canonical SIM-binding groove of SUMO, consistent with the known binding mode.

**Supplementary Figure S10.**
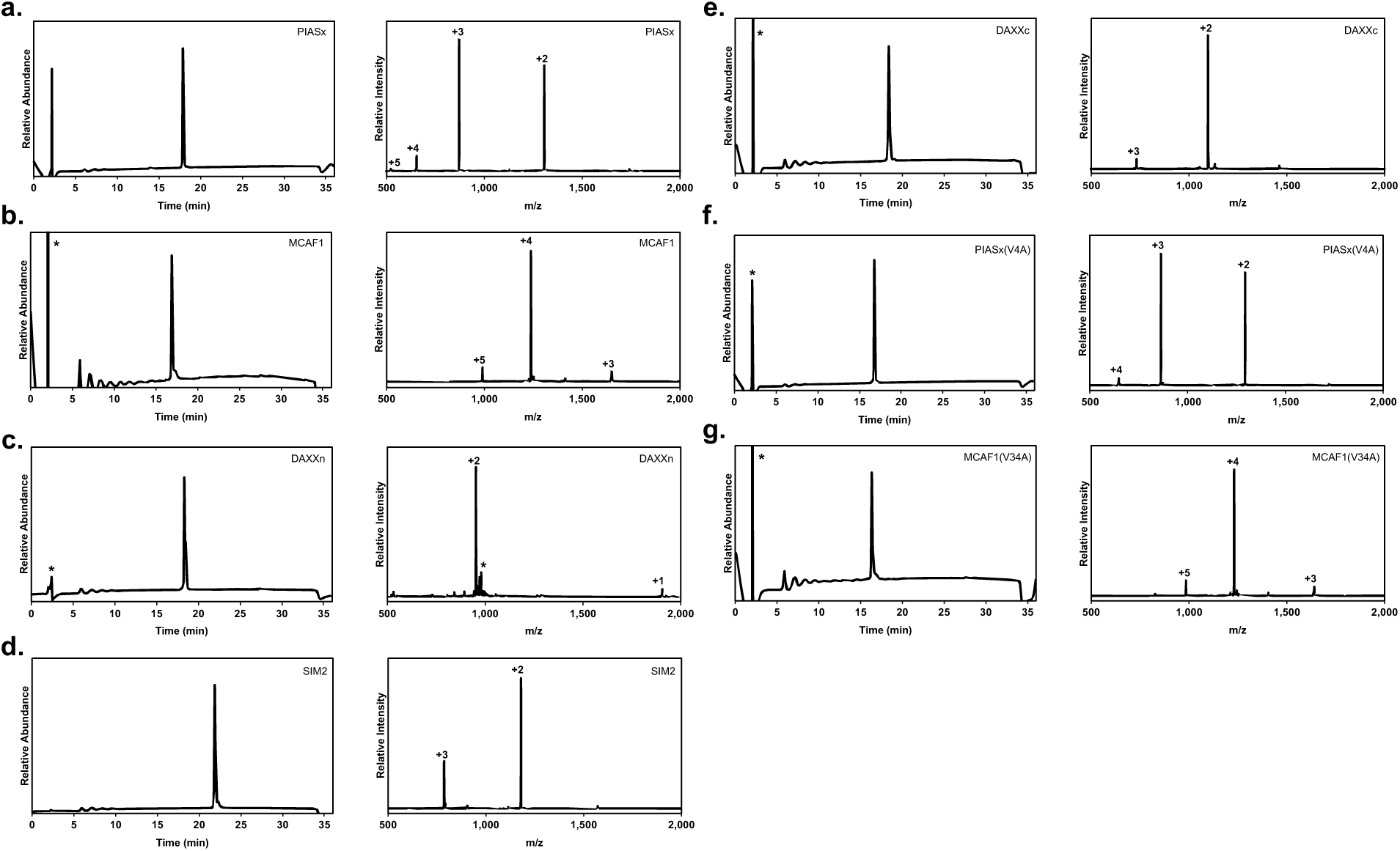
Analytical characterization of synthetic SIM peptides. **a**, Left, C18 analytical HPLC chromatogram of pure PIASx peptide on a gradient of 0-73% B over 30 min. * indicates the injection peak. Right, ESI-MS of pure PIASx peptide. Observed [M]^+^ = 2,613.3 ± 0.4 Da, calculated [M]^+^ = 2,613.3 Da. **b,** Left, C18 analytical HPLC chromatogram of pure MCAF1 peptide on a gradient of 0-73% B over 30 min. * indicates the injection peak. Right, ESI-MS of pure MCAF1 peptide. Observed [M]^+^ = 4,953.1 ± 0.1 Da, calculated [M]^+^ = 4,953.2 Da. **c,** Right, C18 analytical HPLC chromatogram of pure DAXXn peptide on a gradient of 0-73% B over 30 min. * indicates the injection peak. Left, ESI-MS of pure DAXXn peptide. Observed [M]^+^ = 1,907.2 ± 0.3, calculated [M]+ = 1,906.9 Da. * indicates Na^+^ and K^+^ salt adducts. **d,** Left, C18 analytical HPLC chromatogram of pure SIM2 peptide on a gradient of 0-73% B over 30 min. Right, ESI-MS of pure SIM2. Observed [M]^+^ = 2,361.1 ± 0.4 Da, calculated [M]^+^ = 2,360.5 Da. **e.** Left, C18 analytical HPLC chromatogram of pure DAXXc on a gradient of 0-73% B over 30 min. * indicates the injection peak. Right, ESI-MS of pure DAXXc. Observed [M]^+^ = 2,192.2 ± 0.1 Da, calculated [M]^+^ = 2,192.3 Da. **f,** Left, C18 analytical HPLC chromatogram of pure PIASx(V4A) peptide on a gradient of 0-73% B over 30 min. * indicate the buffer peak. Left, ESI-MS of pure PIASx(V4A) peptide. Observed [M]^+^ = 2,585.4 ± 0.4 Da, calculated [M]^+^ = 2,585.7 Da. **g,** Left, C18 analytical HPLC chromatogram of pure MCAF1(V34A) peptide on a gradient of 0-73% B over 30 min. * indicates the injection peak. Right, ESI-MS of pure MCAF1(V34A) peptide. Observed [M]^+^ = 4,925.8 ± 0.9 Da, calculated [M]^+^ = 4,925.1 Da.

**Supplementary Figure S11.**
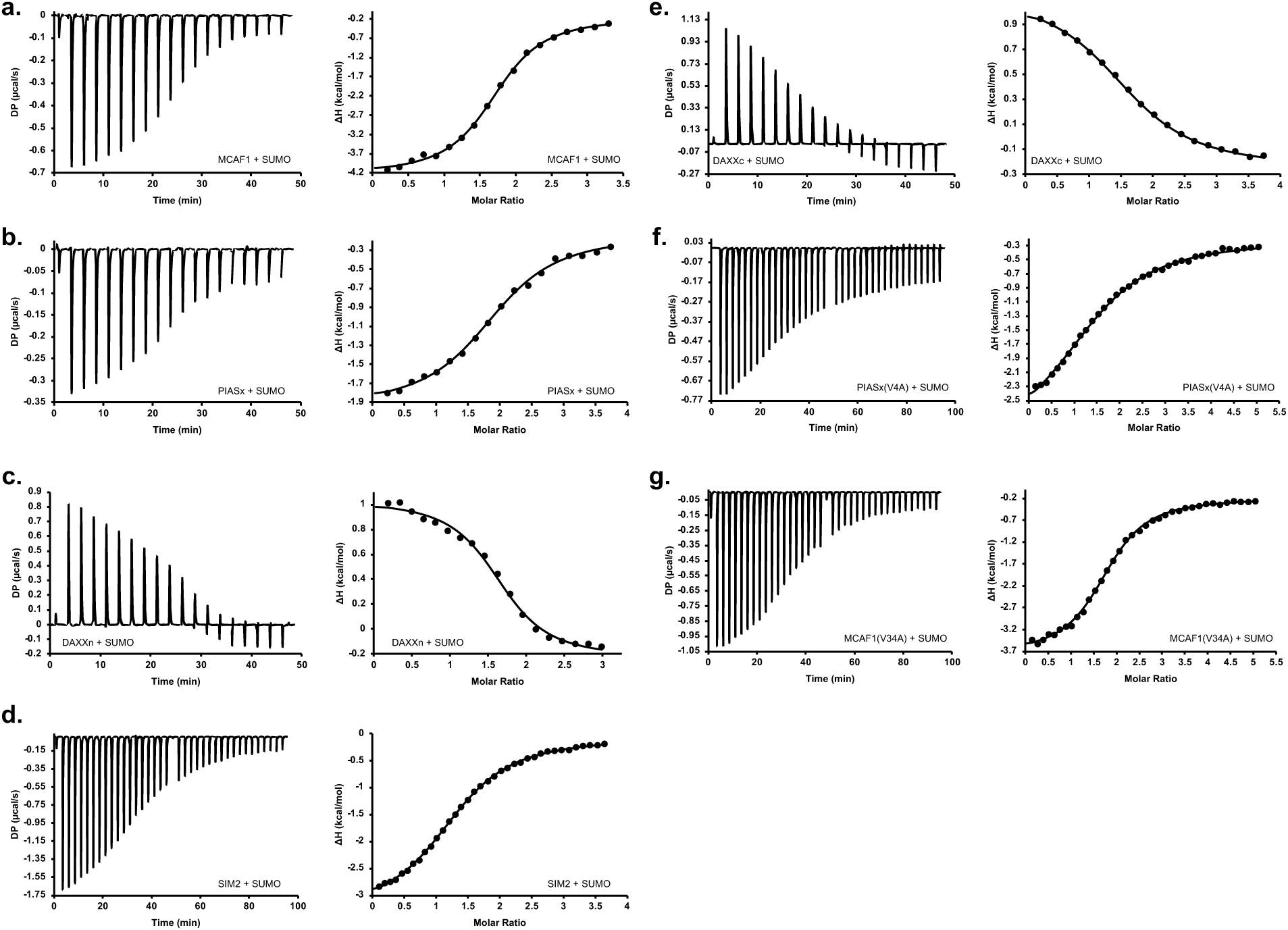
Measurement of SUMO-SIM binding affinity (K_d_) by ITC. **a**, Left, ITC thermogram from titrating the synthetic MCAF1 peptide into SUMO. Right, integrated heats fit to a one-site binding model to obtain K_d_= 3.8 ± 0.2 µM. n= 6 independent replicates and error is s.d. **b,** Left, ITC thermogram from titrating the synthetic PIASx peptide into SUMO. Right, integrated heats fit using a one-site binding model to obtain K_d_= 8.4 ± 0.8 µM. n= 14 independent replicates and error is s.d. **c,** Left, ITC thermogram from titrating the synthetic DAXXn peptide into SUMO. Right, integrated heats fit using a one-site binding model to obtain K_d_= 16.6 ± 1.9 µM. n= 3 independent replicates and error is s.d. **d,** Left, ITC thermogram from titrating the synthetic SIM2 peptide into SUMO. Right, integrated heats fit using a one-site binding model to obtain K_d_= 63.8 ± 1.7 µM. n= 3 independent replicates and error is s.d. **e,** Left, ITC thermogram from titrating the synthetic DAXXc peptide into SUMO. Right, integrated heats fit using a one-site binding model to obtain K_d_= 70.3 ± 3.0 µM. n= 3 independent replicates and error is s.d. **f,** Left, ITC thermogram from titrating the synthetic PIASx(V4A) peptide into SUMO. Right, integrated heats fit using a one-site binding model to obtain K_d_= 57.1 ± 3.5 µM. n= 4 independent replicates and error is s.d. **g,** Left, ITC thermogram from titrating the synthetic MCAF1(V34A) peptide into SUMO. Right, integrated heats fit using a one-site binding model to obtain K_d_= 16.5 ± 0.4 µM. n= 6 independent replicates and error is s.d.

