## Supplementary material for "Scaffold Affinity Tunes Biomolecular Condensate Function": Movie of Fluorescence Recovery after Photobleaching

### Slide 1
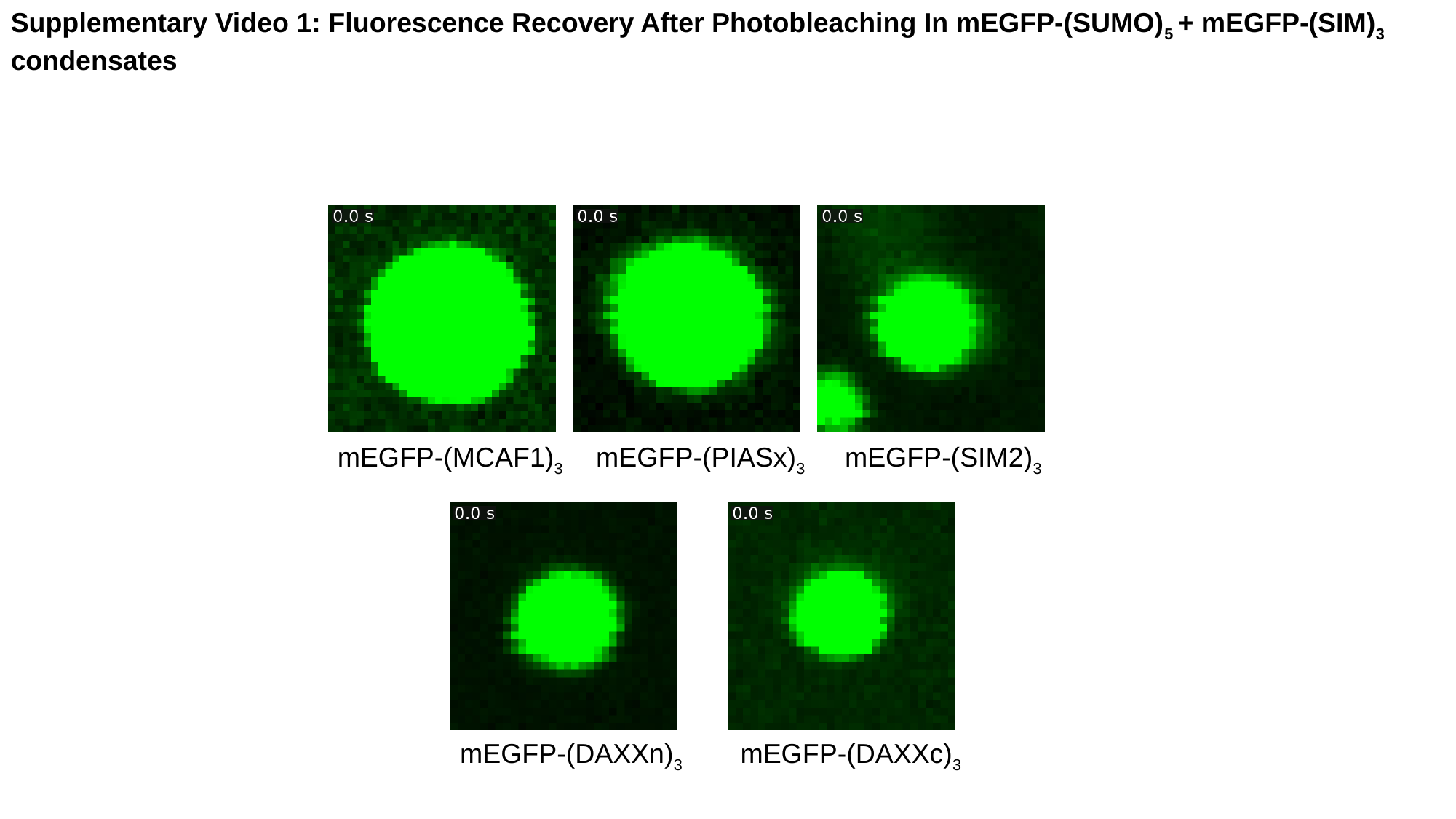

Supplementary Video 1: Fluorescence Recovery After Photobleaching In mEGFP-(SUMO)_5_ + mEGFP-(SIM)_3_ condensates
mEGFP-(MCAF1)_3_
mEGFP-(PIASx)_3_
mEGFP-(SIM2)_3_
mEGFP-(DAXXn)_3_
mEGFP-(DAXXc)_3_
